# The Arabidopsis ATP-Binding Cassette E protein ABCE2 is a conserved component of the translation machinery

**DOI:** 10.1101/2022.05.30.493987

**Authors:** Carla Navarro-Quiles, Eduardo Mateo-Bonmatí, Héctor Candela, Pedro Robles, Antonio Martínez-Laborda, Yolanda Fernández, Jan Šimura, Karin Ljung, Vicente Rubio, María Rosa Ponce, José Luis Micol

**Author notes:** Corresponding author: J.L. Micol.

## Abstract

- ATP-Binding Cassette E (ABCE) proteins dissociate cytoplasmic ribosomes after translation terminates, and contribute to ribosome recycling, thus linking translation termination to initiation. This function has been demonstrated to be essential in animals, fungi, and archaea, but remains unexplored in plants.
- In most species, ABCE is encoded by a single-copy gene; by contrast, *Arabidopsis thaliana* has two *ABCE* paralogs, of which *ABCE2* seems to conserve the ancestral function. We isolated *apiculata7-1 (api7-1)*, a viable, hypomorphic allele of *ABCE2*,which has a pleiotropic morphological phenotype reminiscent of mutations affecting ribosome biogenesis factors and ribosomal proteins. We also studied *api7-2*, a null, recessive lethal allele of *ABCE2*.
- Co-immunoprecipitation experiments showed that ABCE2 physically interacts with components of the translation machinery. An RNA-seq study of the *api7-1* mutant showed increased responses to iron and sulfur starvation. We also found increased transcript levels of genes related to auxin signaling and metabolism.
- Our results support a conserved role for ABCE proteins in ribosome recycling in plants, as previously shown for the animal, fungal, and archaeal lineages. In plants, the ABCE2 protein seems important for general growth and vascular development, likely due to an indirect effect through auxin metabolism.

## INTRODUCTION

Messenger RNA (mRNA) molecules are decoded for protein synthesis by the complex and ancient translation machinery, formed by the ribosome and different sets of translation factors, which function at different translation phases. Translation initiation factors promote the formation of the 70S/80S initiation complex, and the recognition of the mRNA translation start site (Rodnina, 2018; Shirokikh & Preiss, 2018). Translation elongation factors participate in the binding of aminoacyl-tRNAs to the ribosome, the peptide bond formation, and the ulterior release of the deacylated tRNA (Dever *et al*., 2018). Translation termination factors act when the ribosome reaches the translation stop codon and the newly synthesized peptide is released. In this latter phase, the ribosome is dissociated into its 50S/60S and 30S/40S subunits, which are recycled for a new cycle of translation initiation (Hellen, 2018). The ATP-Binding Cassette E (ABCE) proteins are soluble ABC proteins that participate in ribosome recycling and translation initiation, as has been demonstrated for archaea, fungi, and animals, but whose roles in plants remain unexplored (Kashima *et al*., 2014; Young *et al*., 2015; Nürenberg-Goloub *et al*., 2020; Simonetti et al., 2020). Human ABCE1 was first named RNASE L INHIBITOR (RLI) due to its ability to inhibit the activity of RNase L, an enzyme that is only present in mammals (Bisbal et al., 1995).

ABCE proteins contain an iron-sulfur cluster binding domain (FeSD), two nucleotide binding domains (NBD1 and NBD2), and two hinge motifs (Karcher *et al*., 2005; Barthelme *et al*., 2007; Karcher *et al*., 2008). The first hinge motif allows NBD movement to bind and hydrolyze ATP. The second hinge motif and a helix-loop-helix (HLH) mediate the interaction of the ABCE protein with the ribosome after occlusion of two ATP molecules. Once in the ribosome, the ABCE protein displaces its FeSD to split the ribosome, and remains bound to the 30S/40S subunit to prevent a premature recruitment of a 50S/60S subunit during translation initiation. Finally, ATP hydrolysis allows ABCE detachment from the 30S/40S subunit (Barthelme *et al*., 2011; Becker *et al*., 2012; Preis *et al*., 2014; Heuer *et al*., 2017; Nürenberg-Goloub *et al*., 2018; Gouridis *et al*., 2019; Kratzat *et al*., 2021).

In most genomes, the ABCE subfamily is represented by a single-copy gene, usually named *ABCE1*, whose null alleles are lethal, while hypomorphic alleles result in developmental defects and slow-growth phenotypes (Navarro-Quiles *et al*., 2018). *Arabidopsis thaliana* (hereafter referred to as Arabidopsis), however, has two *ABCE* paralogs named *ABCE1* and *ABCE2* (Sánchez-Fernández *et al*., 2001; Verrier *et al*., 2008). Arabidopsis ABCE2 has been studied for its RNA silencing suppression activity (Sarmiento *et al*., 2006; Mõttus *et al*., 2020). In *Cardamine hirsuta*, a close relative of Arabidopsis with compound leaves, only one *ABCE* gene has been identified, *SIMPLE LEAF3 (SIL3)*, which is required for leaflet formation and leaf development. The leaves of homozygotes for the hypomorphic *sil3* mutation are simple and have vascular defects, probably caused by an aberrant auxin transport and homeostasis (Kougioumoutzi *et al*., 2013).

Here, we report a functional analysis of the Arabidopsis *ABCE2* gene. We studied two recessive alleles of *ABCE2:* the hypomorphic and viable *apiculata7-1* (*api7-1*) allele, and the null and lethal *api7-2* allele. The *api7-1* mutant exhibits the typical morphological phenotype caused by mutations in genes encoding ribosome biogenesis factors and ribosomal proteins, which includes aberrant leaf venation patterns. We found by co-immunoprecipitation that ABCE2 physically interacts with components of the translation machinery, and by RNA-seq that its partial loss of function triggers iron and sulfur deficiency responses that might be related to FeS cluster biogenesis, as well as the upregulation of auxin biosynthesis genes. Our observations strongly suggest a conserved role for ABCE proteins in ribosome recycling in plants, as previously shown for the animal, fungal, and archaeal lineages.

## MATERIALS AND METHODS

### Plant materials, growth conditions, and crosses

The *Arabidopsis thaliana* (L.) Heynh. wild-type accessions Landsberg *erecta* (L*er*) and Columbia-0 (Col-0), and the *asymmetric leaves1-1 (as1-1;* N3374; in the Col-1 genetic background) and *as2-1* (N3117; in ER) mutants were initially obtained from the Nottingham Arabidopsis Stock Center (NASC; Nottingham, United Kingdom). We introgressed the *as1-1* and *as2-1* mutations into the Col-0 background by crossing to Col-0 three times. The NASC also provided seeds of the *api7-2* (GABI_509C06; N448798) *(Kleinboelting *et al**.,2012) and *PIN1_pro_:PIN1:GFP DR5_pro_:3XVENUS:N7* (N67931) (Heisler *et al*., 2005) lines. The *ATHB8_pro_:GUS* line (N296) was kindly provided by Simona Baima (Baima *et al*., 1995). The *api7-1* line was isolated in the L*er* background after ethyl methanesulfonate (EMS) mutagenesis in our laboratory, and then backcrossed twice to L*er* (Berná *et al*., 1999). Unless otherwise stated, all the mutants mentioned in this work are homozygous for the mutations indicated. Seed sterilization and sowing, plant culture, crosses, and allelism tests were performed as previously described (Ponce *et al*., 1998; Berná *et al*., 1999; Quesada *et al*., 2000).

### Positional cloning and molecular characterization of *ABCE2* mutant alleles

Genomic DNA was extracted as previously described (Ponce *et al*., 2006). The *ABCE2* gene was cloned as previously described (Mateo-Bonmatí *et al*., 2014). First, we mapped the *api7-1* mutation to a 123.5-kb candidate interval containing 30 genes using a mapping population of 273 F_2_ plants derived from an *api7-1 ×* Col-0 cross, and the primers listed in Table S1, as previously described (Ponce *et al*., 1999; Ponce *et al*., 2006). Then, the whole *api7-1* genome was sequenced by Fasteris (Geneva, Switzerland) using the Illumina HiSeq2000 platform. The bioinformatic analysis of the data was performed as previously described (Mateo-Bonmatí *et al*., 2014).

Discrimination between the wild-type *ABCE2* and *api7-1* mutant alleles was done by PCR with the api7-1_F/R primers (Table S1), followed by restriction with *Eco*57I (Thermo Fisher Scientific), as the *api7-1* mutation (CT<u>C</u>CAG→CT<u>T</u>CAG) creates an *Eco*57I restriction site. The presence and position of the *api7-2* T-DNA insertion in the GABI_509C06 line was confirmed by PCR amplification and Sanger sequencing, respectively, using gene-specific primers and the o8409 primer for the GABI-Kat T-DNA (Table S1).

### Gene constructs and plant transformation

All inserts were PCR amplified from Col-0 genomic DNA using Phusion High Fidelity DNA Polymerase (Thermo Fisher Scientific) and primers that contained *att*B sites at their 5’ ends (Table S1). PCR products were purified using an Illustra GFX PCR DNA and Gel Band Purification Kit (Cytiva), and then cloned into the pGEM-T Easy221 vector, transferred to *Escherichia coli* DH5α, and subcloned into the pEarleyGate 101, pMDC83, or pMDC107 destination vectors (Curtis & Grossniklaus, 2003; Earley *et al*., 2006) as previously described (Mateo-Bonmatí *et al*., 2018).

All constructs were transferred to electrocompetent *Agrobacterium tumefaciens* GV3101 (C58C1 Rif^R^) cells, which were used to transform L*er* or *api7-1* plants by the floral dip method (Clough & Bent, 1998). Putative transgenic plants were selected on plates supplemented with 15 μg·ml^−1^ hygromycin B (Thermo Fisher Scientific, Invitrogen).

To obtain the GSRhino-TAP-tagged ABCE2 fusion, a pGEM-T Easy221 vector harboring the *ABCE2* coding sequence, together with the pEN-L4-2-R1 and pEN-R2-GSRhinotag-L3 entry vectors were subcloned into the pKCTAP destination vector as previously described (Van Leene *et al*., 2015). Transformation of Arabidopsis cell cultures was performed as previously described (Van Leene *et al*., 2015).

### Phenotypic analysis and morphometry

Photographs were taken with a Nikon SMZ1500 stereomicroscope equipped with a Nikon DXM1200F digital camera. For larger specimens, four to five partial images from the same plant were taken and merged using the Photomerge tool of Adobe Photoshop CS3 software. For rosette size, rosette silhouettes were drawn on the screen of a Cintiq 18SX Interactive Pen Display (Wacom) using Adobe Photoshop CS3, and their sizes were measured with the NIS Elements AR 3.1 image analysis package (Nikon). Root length was measured per triplicate from photographs with the Freehand line tool from Fiji software (https://imagej.net/ImageJ) (Schindelin *et al*., 2012). Shoot length was measured *in vivo* with a millimeter ruler, from the soil to the apex of the main shoot. Chlorophyll *a* and *b* and carotenoids were extracted and spectrophotometrically determined as previously described (Wellburn, 1994; Micol-Ponce *et al*., 2020), and normalized to the amount of collected tissue.

### Differential interference contrast and bright-field microscopy, and GUS analyses

For differential interference contrast (DIC) and bright-field microscopy, all samples were cleared, mounted, and photographed as previously described (Candela *et al*., 1999). Micrographs of venation patterns, and leaf primordia expressing *ATHB8_pro_:GUS* were taken under bright field with a Nikon D-Eclipse C1 confocal microscope equipped with a Nikon DS-Ri1 camera, using the NIS-Elements AR 3.1 software (Nikon). Diagrams from leaf cells and venation patterns, and morphometric analysis of leaf cells were obtained as previously described (Pérez-Pérez *et al*., 2011; Mateo-Bonmatí *et al*., 2018). For venation pattern morphometry, the phenoVein (http://www.plant-image-analysis.org) (Bühler *et al*., 2015) software was used. Leaf lamina circularity was calculated as *4 · π · area/perimeter^2^*.Lamina area and perimeter were measured on diagrams from the leaf lamina with the Fiji Wand tool. GUS assays were performed as previously described (Robles *et al*., 2010).

### Confocal microscopy and fluorescence quantification

Confocal laser scanning microscopy images were obtained using a Nikon D-Eclipse C1 confocal microscope equipped with a Nikon DS-Ri1 camera and processed with the operator software EZ-C1 (Nikon). Visualization of the fluorescent proteins and dyes was performed on primary roots mounted with deionized water on glass slides. Fluorescent proteins, 4’,6-diamidino-2-phenylindole (DAPI), and propidium iodide were visualized as described in Table S2. For fluorescence quantification of the *PIN1_pro_:PIN1:GFP* and *DR5_pro_:3XVENUS:N7* protein products, wild-type and *api7-1* seedlings homozygous for these transgenes were grown vertically on the same Petri dishes under identical conditions for 5 days. Image acquisition was performed using a 40× objective with a 0.75 numerical aperture. The dwell time was set at 2.16 and 1.68 μs for PIN1:GFP and 3XVENUS:N7, respectively. Four images were acquired and averaged per optical section. Five optical sections encompassing 4 μm from the innermost root layers were photographed. Acquired images (.ids files) were used to generate flat images (.tiff files) with Fiji, by stacking the optical sections from the fluorescent protein channel. Fluorescence quantification was performed using the Fiji Mean gray value measurement.

### RNA isolation, cDNA synthesis, and quantitative PCR

Samples for RNA extraction were collected on ice and immediately frozen for storage at −80°C until use. RNA was isolated using TRIzol (Thermo Fisher Scientific, Invitrogen). Removal of contaminating DNA, cDNA synthesis, and quantitative PCR (qPCR) were performed as previously described (Mateo-Bonmatí *et al*., 2018). The qPCR was performed as follows: 2 min at 50°C, 10 min at 95°C, followed by 41 cycles of 15 s at 95°C and 1 min at 60°C, and a final step of 15 s at 95°C, and *ACTIN2 (ACT2)* was used as an internal control for relative expression analysis (Moschopoulos *et al*., 2012). Three biological replicates, each with three technical replicates, were analyzed per genotype. Relative quantification of gene expression data was performed using the comparative C_T_ method (2^−δδC_T_^) (Schmittgen & Livak, 2008).

### RNA-seq analysis

Total RNA was isolated from 100 mg of L*er* and *api7-1* rosettes collected 14 days after stratification (das) using TRIzol. RNA concentration and quality were assessed using a 2100 Bioanalyzer (Agilent Genomics) with an RNA 600 Nano Kit (Agilent Technologies) as previously described (Mateo-Bonmatí *et al*., 2018). Three biological replicates per genotype, with more than 14 μg of total RNA per sample, and an RNA integrity number (RIN) higher than 7, were sent to Novogene (Cambridge, United Kingdom) for massive parallel sequencing. Sequencing libraries were generated using NEBNext Ultra RNA Library Prep Kit for Illumina (New England Biolabs) and fed into a NovaSeq 6000 Illumina platform with a S4 Flow Cell type, which produced paired-end reads of 150 bp (Table S3). Read mapping to the Arabidopsis genome (TAIR10) using the 2.0.5 version of HISAT2 (Kim *et al*., 2019), with default parameters, and the identification of differentially expressed genes between L*er* and *api7-1* with the 1.22.2 version of DESeq2 R package (Love *et al*., 2014) were performed by Novogene. Genes with a *P*-value < 0.05 adjusted with the Benjamini and Hochberg’s method, and with a fold change >1.5 were considered differentially expressed. The gene ontology (GO) enrichment analysis of the differentially expressed genes was performed with the online tool DAVID (https://david.ncifcrf.gov/home.jsp) (Huang et al., 2009a; Huang *et al*., 2009b).

### Indol-3-acetic acid metabolite profiling

Shoots, whole roots, and primary root tips (3 mm approximately) were collected 9 das from vertically grown seedlings. These samples were rapidly weighed and frozen in liquid nitrogen. Extraction and purification of the targeted compounds (anthranilate [Ant], tryptophan [Trp], indole-3-acetonitrile [IAN], indol-3-acetic acid [IAA], IAA-glucose [IAA-Glc], IAA-aspartate [IAA-Asp], IAA-glutamate [IAA-Glu], 2-oxindole-3-acetic acid [oxIAA], and oxIAA-glucoside [oxIAA-Glc]) were performed as previously described (Novák *et al*., 2012;Mateo-Bonmatí *et al*., 2021). Ultra-high performance liquid chromatography followed by tandem mass spectrometry (UHPLC-MS/MS) analysis was performed as previously described (Pěnčík *et al*., 2018).

### Co-immunoprecipitation assay

For protein extraction, 700 mg of whole *api7-1 35Spro:ABCE2:YFP* seedlings were collected 10 das per biological replicate. The tissue was crosslinked with 1× phosphate-saline buffer containing 1% (v/v) formaldehyde as previously described (Poza-Viejo *et al*., 2019). For protein extraction, the tissue was ground to a fine powder with liquid nitrogen and then resuspended in a lysis buffer (50 mM Tris-HCl, pH 7.5; 0.1% [v/v] IGEPAL CA-630 [Sigma-Aldrich]; 2 mM phenylmethylsulfonyl fluoride [PMSF; Sigma-Aldrich]; 150 mM NaCl; and a cOmplete protease inhibitor cocktail tablet [Sigma-Aldrich]) using a vortexer. After incubation on ice for 10 min, the samples were centrifuged at 4°C and the supernatants were used as protein extracts. Co-immunoprecipitation was performed with the μMACS GFP Isolation Kit (Milteny Biotec) using protein extracts from three biological replicates. The immunoprecipitation of the ABCE2:YFP fusion protein was checked by western blotting using an anti-GFP-HRP antibody (Milteny Biotec), and the WesternSure chemiluminiscent substrate on a C-DiGit Blot Scanner (LI·COR).

The co-immunoprecipitates were analyzed by liquid chromatography followed by electrospray ionization and MS/MS (LC-ESI-MS/MS) at the Centro Nacional de Biotecnología (CNB) Proteomics facility (Madrid, Spain). Tandem mass spectra were searched against Araport11 using the MASCOT search engine (Matrix Science, http://www.matrixscience.com/). Peptide sequences identified with a false discovery rate (FDR) < 1% were considered statistically valid. Proteins identified with at least 2 peptides without overlapping sequences (unique peptides) in at least 2 biological replicates (namely, at least 4 peptides) were considered identified with high confidence. To search for potential ABCE2:YFP interactors, proteins whose subcellular localization was not predicted to be cytoplasmic by SUBA4 (https://suba.live/) (Hooper *et al*.,2014; Hooper *et al*., 2017) were discarded, with the exception of At2g20830, which is predicted to localize to mitochondria (see Results). To further discard potential false positive interactions, all the proteins identified in three other co-immunoprecipitations of GFP-fused proteins performed in our laboratory under identical conditions to that of ABCE2:YFP, but functionally unrelated, were used to create a subtract list. Proteins identified in ABCE2:YFP samples with at least twice the number of peptides assigned to the same protein in the subtract list were considered enriched. The rest of the proteins, which contained a more similar number of peptides between the ABCE2:YFP list and the subtract list, were considered false positives and discarded. In addition, there were few proteins that were solely identified in ABCE2:YFP samples.

### Tandem affinity purification assay

Tandem affinity purification (TAP) of the GSRhino-TAP-tagged ABCE2 fusion from Arabidopsis cell suspension cultures was performed as previously described (Van Leene *et al*., 2015; García-León *et al*., 2018). Proteins were identified by nano LC-MS/MS at the CNB. Tandem mass spectra were searched against Araport11 using the MASCOT search engine. Proteins identified with at least 1 unique peptide with a MASCOT score higher than 25 (*p* < 0.05) were considered to be valid. Proteins identified with at least 1 unique peptide in the 2 biological replicates or 2 unique peptides in 1 biological replicate were considered identified with high confidence. We discarded as putative ABCE2 interactors those proteins that were not predicted to be cytoplasmic by SUBA4.

### Bioinformatic analyses

The identity and similarity values between conserved proteins were obtained from global pairwise sequence alignments performed with EMBOSS Needle (https://www.ebi.ac.uk/Tools/psa/emboss_needle/) (Madeira *et al*., 2019). The multiple sequence alignment of ABCE orthologs was obtained with Clustal Omega (https://www.ebi.ac.uk/Tools/msa/clustalo/) (Madeira *et al*., 2019).

A TBLASTN search was performed to identify *ABCE* genes within eudicots (taxid:71240) against the sequences contained in the Nucleotide collection database at the National Center for Biotechnology Information BLASTP server (NCBI; https://blast.ncbi.nlm.nih.gov/Blast.cgi) (Altschul *et al*., 1997) using *Arabidopsis thaliana* ABCE2 protein as the query (NP_193656). The phylogenetic analysis was performed using the NCBI accession numbers listed in Table S4 with MEGA X software (Kumar *et al*., 2018): the multiple sequence alignment and the phylogenetic tree were obtained using codon recognition with Muscle (Edgar, 2004b; Edgar, 2004a), and the Neighbor-Joining method (Saitou & Nei, 1987), respectively.

### Accession numbers

Sequence data can be found at The Arabidopsis Information Resource (https://www.arabidopsis.org/) under the following accession numbers: *ABCE1* (At3g13640), *ABCE2* (At4g19210), *ACT2* (At3g18780), *ATHB8* (At4g32880), *OTC* (At1g75330), and *PIN1* (At1g73590).

## RESULTS

### The *apiculata7-1* mutant exhibits a pleiotropic morphological phenotype

The *apiculata7-1 (api7-1)* mutant, which we initially named *api7*, was isolated in a previous large-scale screen for EMS-induced mutations affecting leaf development (Berná *et al*.,1999). Its pleiotropic morphological phenotype includes a small rosette, a short primary root, and a delay in main stem growth (Fig. 1; Fig. S1a). The *api7-1* inflorescences and siliques are seemingly normal (Fig. S1b–g). The rosette leaves are pointed, indented, and pale, and contain a reduced amount of photosynthetic pigments, compared to its wild-type L*er* (Fig. 1; Fig. S1h). *api7-1* first-node leaves show a marked reduction in cell size in the abaxial and adaxial epidermal layers, but not in the palisade mesophyll (Fig. S2).

**Fig. 1.**
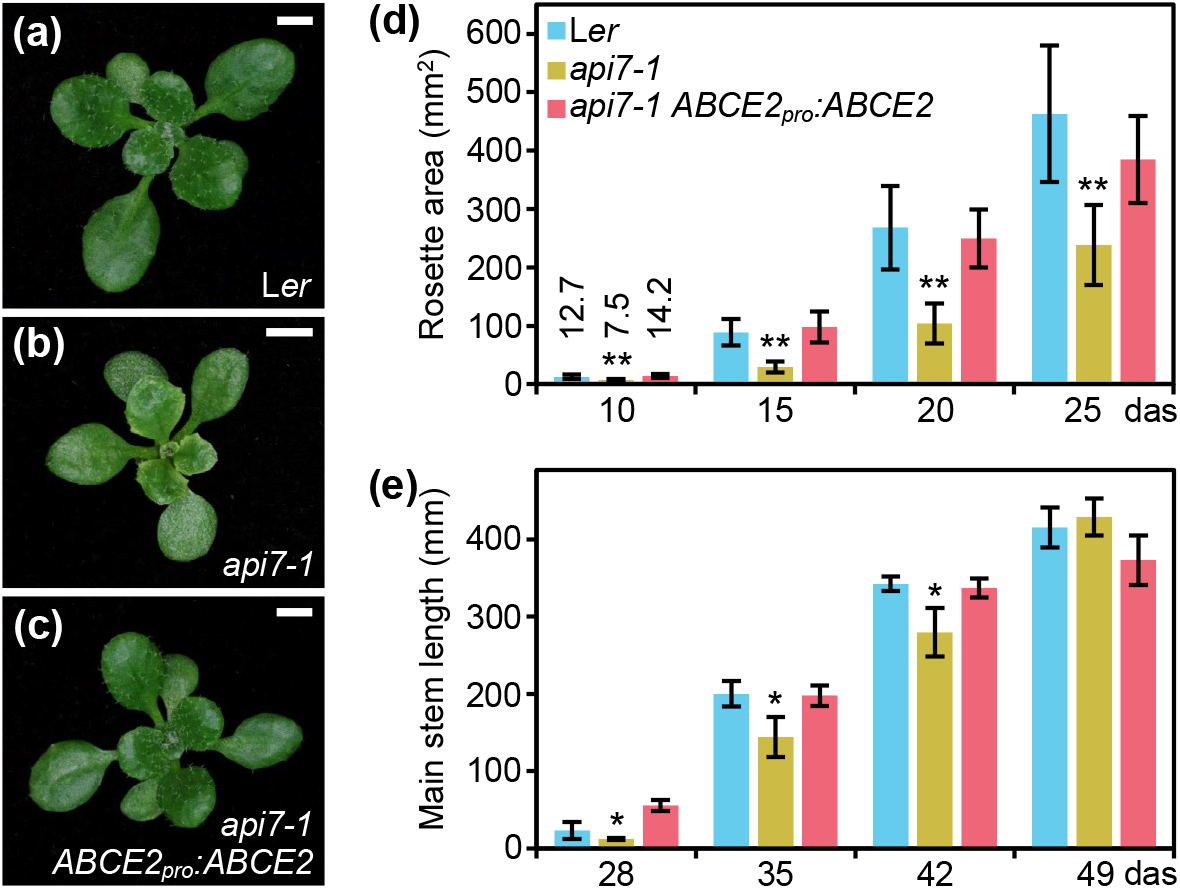
Morphological phenotype of the *api7-1* mutant. (a-c) Rosettes from (a) the wild-type *Ler*,(b) the *api7-1* mutant, and (c) an *api7-1 ABCE2_pro_:ABCE2* mutant and transgenic plant. Pictures were taken 16 days after stratification (das). Scale bars indicate 2 mm. (d,e) Growth progression of (d) rosette area and (e) main stem length. Bars indicate (d) mean and (e) median values. Error bars represent (d) standard deviation and (e) median absolute deviation. Asterisks indicate a significant difference with L*er* in a (d) Student’s *t* test (10 < n < 17) or (e) Mann-Whitney *U* test (n = 8) (\**P*< 0.05, \*\**P*< 0.001).

The pleiotropic phenotype of *api7-1* plants is reminiscent of mutants carrying loss-of-function alleles of genes encoding ribosomal proteins or ribosome biogenesis factors (Byrne, 2009; Horiguchi *et al*., 2011; Rosado *et al*., 2012; Weis *et al*., 2015; Micol-Ponce *et al*., 2018). As these mutations usually alter leaf vascular development, we cleared *api7-1* and L*er* leaves with chloral hydrate, and observed their venation patterns. We confirmed that *api7-1* fully expanded first-node and, to a lesser extent, third-node rosette leaves, contain fewer higher-order veins, and more prominent indentations and vascularized hydathodes, particularly in the leaf apex, than L*er* leaves (Fig. 2; Fig. S3). In contrast, these phenotypic traits seemed to be unaffected on *api7-1* cotyledons, cauline leaves, sepals, and petals (Fig. S4; Table S5). The *ARABIDOPSIS THALIANA HOMEOBOX GENE 8* (*ATHB8*) gene is expressed in pre-procambial cells that will differentiate into veins (Baima *et al*., 1995). To determine the stage at which *api7-1* leaf venation pattern formation diverged from that of L*er*, we crossed *api7-1* plants to an *ATHB8_pro_:GUS* line, and studied the expression of the transgene in cleared first-node rosette leaf primordia of *api7-1 ATHB8_pro_:GUS* plants. Consistent with the slow growth phenotype of *api7-1* plants, we observed a delay in the emergence of first-node leaves (Fig. S5). In addition, *api7-1* primordia retained high GUS activity at their apical region even after the formation of the whole midvein (Fig. S5m,n), suggesting that an increased vascular differentiation in that region is responsible of the vascular phenotype of mature *api7-1* leaves (Fig. S3).

**Fig. 2.**
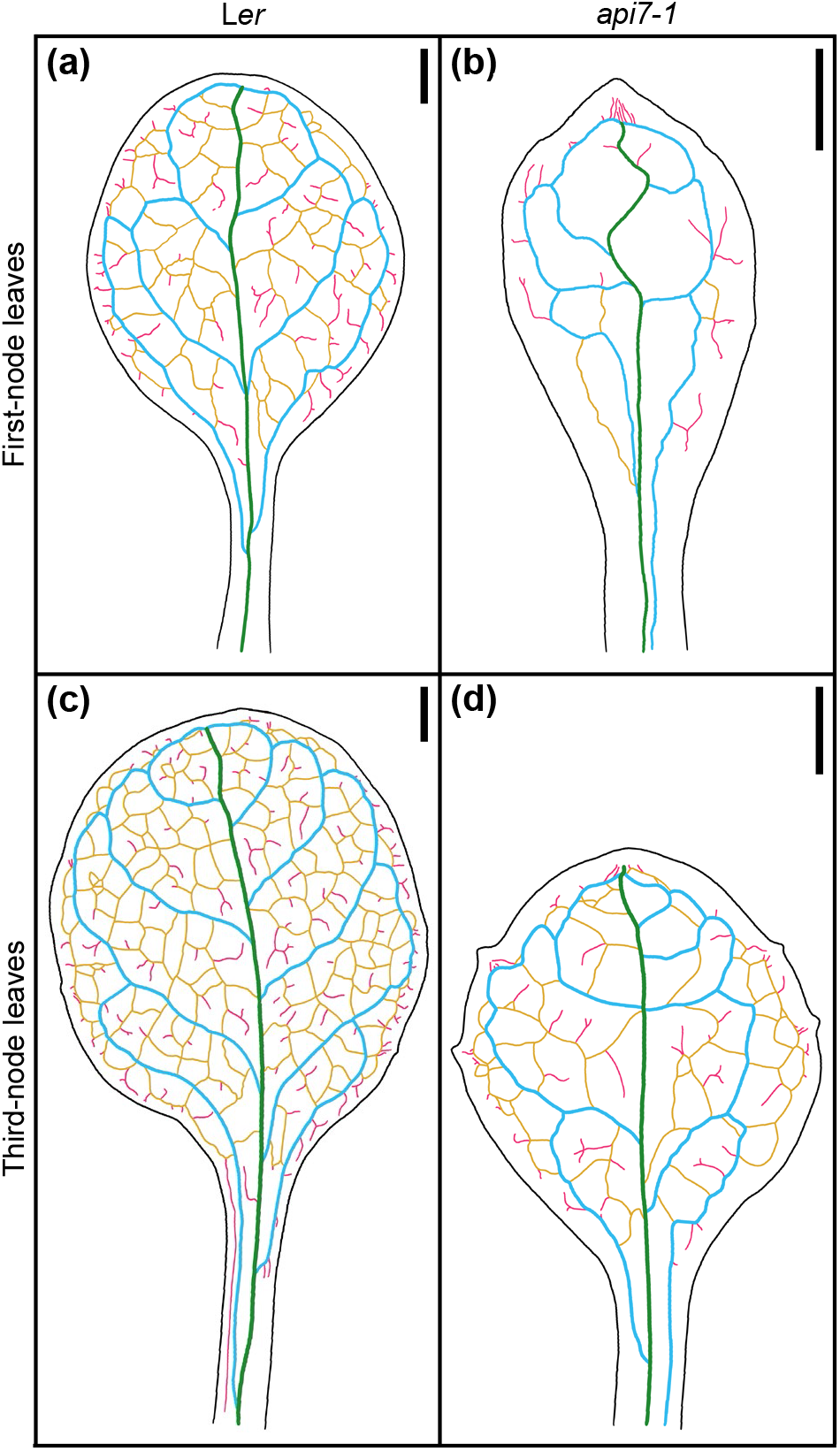
Venation pattern of *api7-1* first- and third-node leaves. Representative diagrams of mature (a,b) first- and (c,d) third-node leaves from (a,c) L*er* and (b,d) *api7-1* plants. Margins were drawn in black, primary veins in green, secondary veins in blue, higher-order connected veins in yellow, and higher-order disconnected veins in pink. Organs were collected 21 das. Scale bars indicate 1 mm.

*ASYMMETRIC LEAVES 1 (AS1)* and *AS2* encode transcription factors involved in leaf dorsoventral patterning. Double mutant combinations of *as1* or *as2* with mutations in genes encoding ribosomal proteins or other components of the translation machinery usually produce synergistic phenotypes. These phenotypes are easily distinguished by the presence of trumpet-shaped (peltate) or radial leaves originated by partial or complete loss of dorsoventrality, respectively (Pinon *et al*., 2008; Yao *et al*., 2008; Horiguchi *et al*., 2011;Moschopoulos *et al*., 2012; Casanova-Sáez *et al*., 2014; Mateo-Bonmatí *et al*., 2015). We obtained *api7-1 as1-1* and *api7-1 as2-1* double mutants in the Col-0 background; these double mutants exhibited additive and synergistic phenotypes, respectively (Fig. S6). The presence of radial leaves in *api7-1 as2-1* plants further supports a role for API7 in translation (Fig. S6f,g).

### *api7-1* is a viable mutant allele of the *ABCE2* gene

The *api7-1* mutation was previously mapped to chromosome 4 (Robles & Micol, 2001). To identify the mutated gene, we combined map-based cloning and next-generation sequencing, as previously described (Mateo-Bonmatí *et al*., 2014). First, we performed linkage analysis of an F_2_ mapping population, which allowed us to delimit a candidate interval encompassing 30 annotated genes (Fig. 3a). We then sequenced the whole *api7-1* genome and identified 4 EMS-type nucleotide substitutions within the candidate interval (Table S6). Only one of these, a C→T transition in At4g19210, was predicted to be a missense mutation causing a Pro138→Ser substitution (Fig. 3b). The At4g19210 gene encodes ABCE2, a protein of 605 amino acids (68.39 kDa). The Pro138 residue, at the beginning of the HLH motif located within NBD1, is conserved across all eukaryotic ABCE proteins tested, except in *Caenorhabditis elegans* (Fig. S7), in which it seems to have evolved more divergently (Chen *et al*., 2006). The conservation of this residue suggests that it is necessary for the proper function of ABCE proteins, probably for the interactions with the ribosome, which mainly occur through the HLH and hinge motifs (Heuer *et al*., 2017;Nürenberg-Goloub *et al*., 2020; Kratzat *et al*., 2021).

**Fig. 3.**
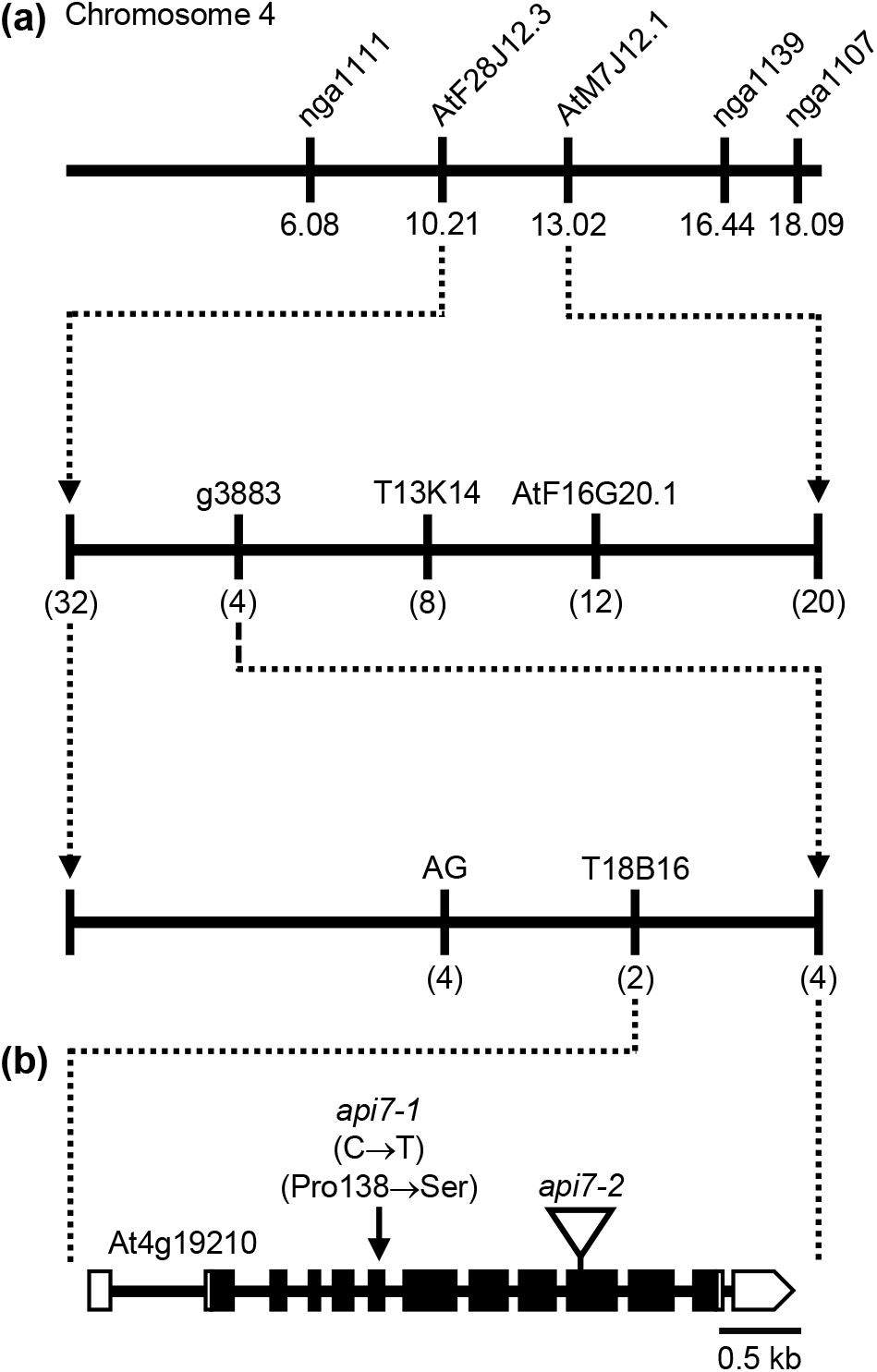
Fine mapping by linkage analysis of the *api7-1* mutation. (a) A mapping population of 273 F_2_ plants derived from an *api7-1 **×*** Col-0 cross allowed us to delimit a candidate region of 123.5 kb in chromosome 4, flanked by the T18B16 and g3883 markers. The names and physical map positions of the molecular markers used for linkage analysis are shown. All values outside parentheses indicate Mb. The number of recombinant chromosomes found (from a total of 546 chromosomes analyzed) is indicated in parentheses. (b) Structure of the At4g19210 *(ABCE2)*gene, located within the candidate region, with indication of the nature and position of the *api7*mutations studied in this work. Boxes and lines indicate exons and introns, respectively. White boxes represent UTRs. The arrow indicates the *api7-1*point mutation. The triangle indicates the *api7-2* T-DNA insertion (GABI_509C06).

To confirm that the mutation found in At4g19210 causes the phenotype of the *api7-1* mutant, we obtained the *ABCE2_pro_:ABCE2* transgene, which was transferred into *api7-1* plants. This transgene completely restored the wild-type rosette leaf shape and stem height (Fig. 1c–e), as well as the photosynthetic pigment content (Fig. S1h). The *ABCE2_pro_:ABCE2* transgene partially restored the root length, and leaf epidermal cell sizes (Fig. S1a, S2). To provide further confirmation that *api7-1* is an allele of *ABCE2*, we performed an allelism test using GABI_509C06 plants (Kleinboelting *et al*., 2012), which were heterozygous for a T-DNA insertion in the 10^th^ exon of At4g19210 (Fig. 3b). We named *api7-2* the insertional allele in GABI_509C06. In the F2 population of this cross, no *api7-2/api7-2* plants were found, and *api7-1/api7-2* and *api7-1/api7-1* plants were phenotypically similar, confirming that these mutations are allelic and that loss of function of *ABCE2* is responsible for the phenotype of the *api7-1* mutant (Fig. S8a–c).

The absence of *api7-2/api7-2* plants derived from GABI_509C06 seeds, and of ungerminated seeds in the F1 progeny of selfed heterozygous *ABCE2/api7-2* plants, suggested an early lethality of this mutant allele. We dissected immature siliques from *ABCE2/api7-2* plants and found 21.95% aborted seeds (*n* = 328), which fits a 1:3 Mendelian segregation ratio (χ^2^= 1.63; *P*-value = 0.202; degrees of freedom = 1). Col-0 siliques showed 1.37% aborted ovules (*n* = 148; Fig. S8d,e). The lethality caused by *api7-2* suggests that it is a null allele of *ABCE2*, while *api7-1* is hypomorphic.

### The Arabidopsis genome contains two partially redundant *ABCE* paralogs

To gather information about the origin of the two Arabidopsis *ABCE* paralogs, we performed a phylogenetic analysis of *ABCE* coding sequences from some Rosidae species (rosids; Fig. S9). Among them, we found that other Brassicaceae genomes also encode ABCE1 and ABCE2 proteins, but only *ABCE2* was identified in *Cardamine hirsuta*. Consistent with the whole-genome triplication in *Brassica rapa* (Zhang *et al*., 2018), we found two and three *Brassica rapa ABCE1* and *ABCE2* sequences, respectively. All Brassicaceae *ABCE1* genes grouped together in the phylogenetic tree, and separately from their *ABCE2* paralogs, which formed other subclade. Although both *ABCE1* and *ABCE2* paralogs have been conserved, *ABCE1* orthologs have evolved more rapidly than their *ABCE2* paralogs, whose short evolutionary distances indicate that they are under strong evolutionary pressure, as expected for an essential gene.

As previously described (Braz *et al*., 2004; Sarmiento *et al*., 2006), we observed that *ABCE2* is highly expressed throughout all Arabidopsis developmental stages. By contrast, the expression levels of its *ABCE1* paralog are very low in all studied organs, in which first-node leaves and flowers show the lowest and highest expression levels, respectively (Fig. S10a,b). The expression level of *ABCE1* in *api7-1* rosettes was the same as in L*er*, showing that ABCE1 cannot compensate for the partial loss of ABCE2 function in rosettes (Fig. S10c). However, *api7-1* flowers, where we observed the highest *ABCE1* expression levels, do not show apparent aberrations (Fig. S1c), suggesting that ABCE1 and ABCE2 might play similar roles during flower development.

ABCE1 and ABCE2 proteins share 80.8% identity, suggesting that ABCE1 and ABCE2 might be functionally equivalent. To test this hypothesis, we performed a promoter swapping assay between the *ABCE1* and *ABCE2* genes (Fig. 4). As expected from the lower expression levels driven by the *ABCE1* promoter, *api7-1 ABCE1_pro_:ABCE2* plants were indistinguishable from *api7-1* mutants, highlighting that correct protein levels are as important as the correct sequence for normal *ABCE2* function. In contrast, the *ABCE2_pro_:ABCE1* transgene partially rescued the *api7-1* phenotype, showing that the ABCE1 and ABCE2 proteins are functionally redundant. Further supporting equivalent functions for ABCE1 and ABCE2, the constitutive expression of *ABCE1* with a *35S_pro_:ABCE1* transgene fully restored a wild-type phenotype in *api7-1* rosettes (Fig. S11).

**Fig. 4.**
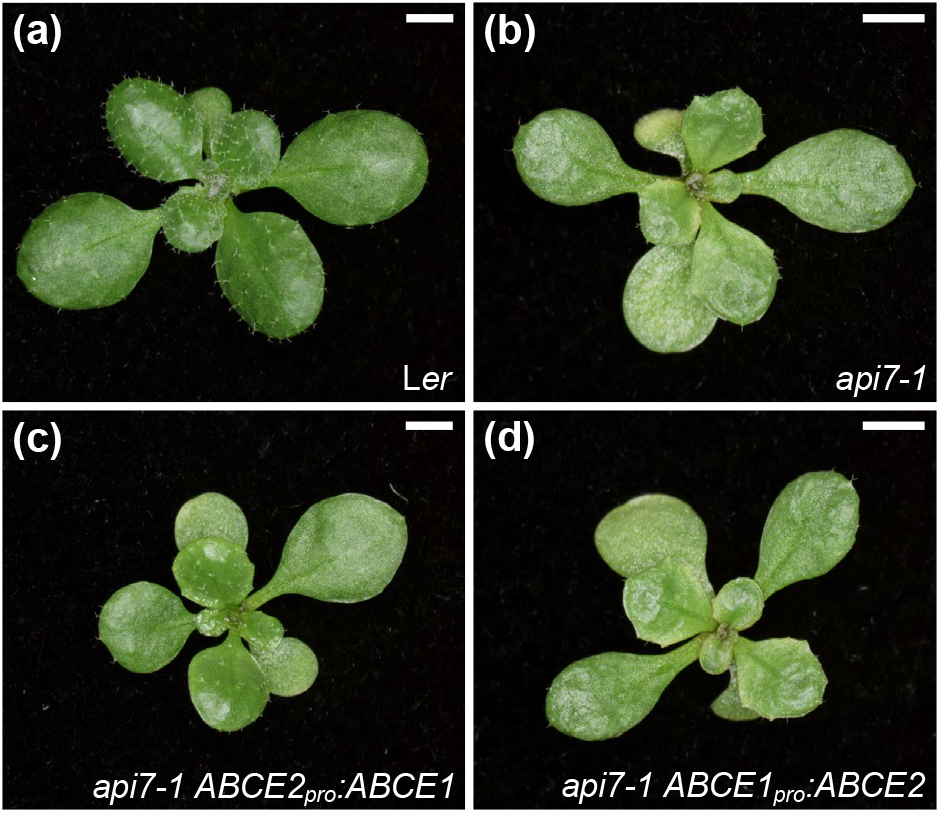
Effects of the *ABCE2_pro_:ABCE1* and *ABCE1_pro_:ABCE2* transgenes on the morphological phenotype of the *api7-1* mutant. Rosettes from (a) L*er*, (b) *api7-1*, (c) *api7-1 ABCE2_pro_:ABCE1*,and (d) *api7-1 ABCE1_pro_:ABCE2* plants. Pictures were taken 14 das. Scale bars indicate 2 mm.

### ABCE2 is a cytoplasmic protein that physically associates with components of the translation machinery

To determine the subcellular localization of Arabidopsis ABCE2, we obtained in-frame translational fusions of ABCE2 to GFP and YFP, driven by the 35S promoter: *35S_pro_:ABCE2:GFP* and *35S_pro_:ABCE2:YFP*. We visualized the ABCE2:GFP fusion protein in the cytoplasm of root cells treated with propidium iodide, which mainly stains cell walls, and the ABCE2:YFP fusion protein in roots stained with the nucleoplasm dye DAPI, and confirmed the nuclear exclusion of ABCE2 (Fig. 5).

**Fig. 5.**
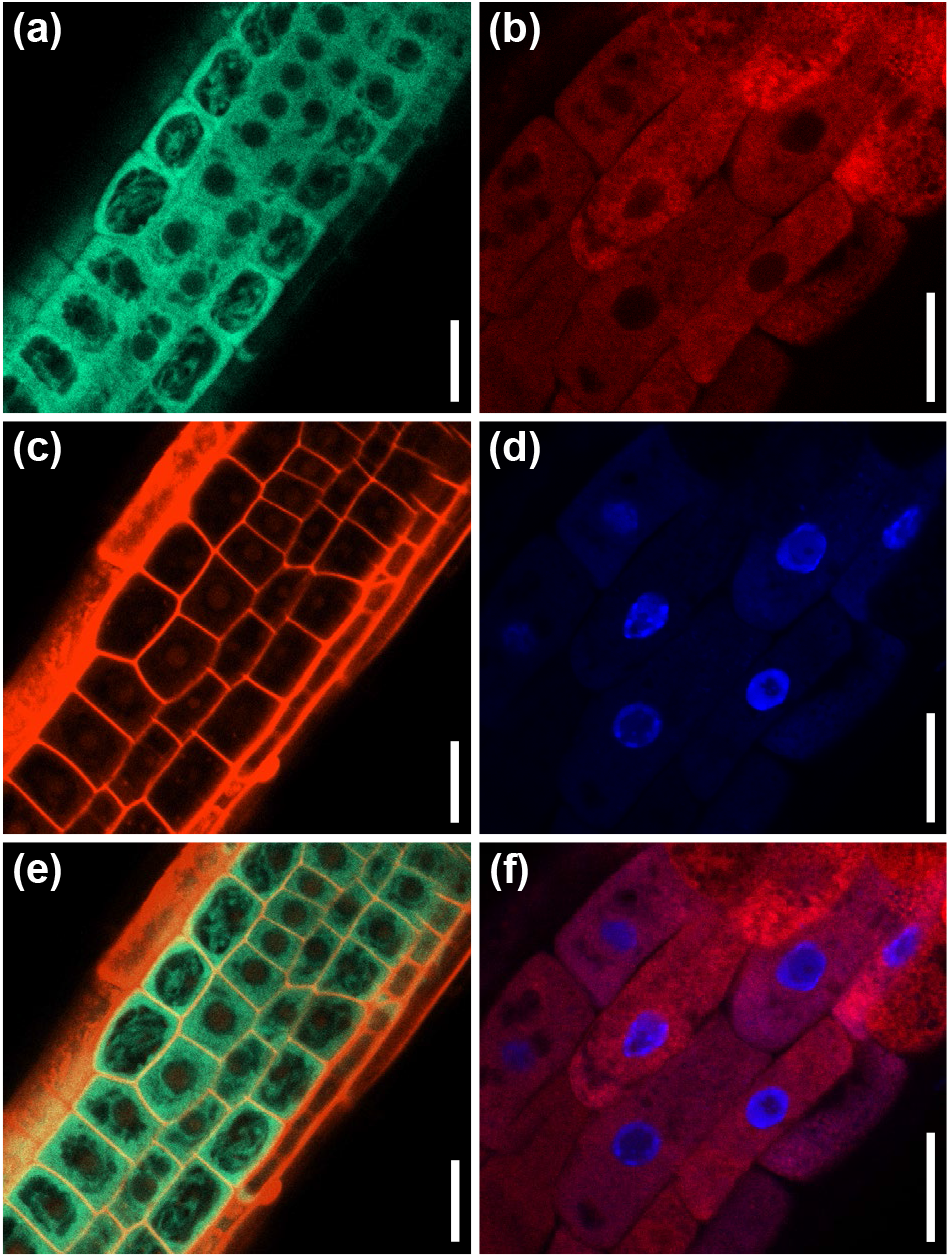
Subcellular localization of the ABCE2 protein in cells from the root elongation zone. Confocal laser scanning micrographs of (a,c,e) L*er 35S_pro_:ABCE2:GFP* and (b,d,f) *api7-1 35S_pro_:ABCE2:YFP* transgenic plants. Fluorescent signals correspond to (a) GFP, (b) YFP, (c) propidium iodide, and (d) DAPI staining, and (e,f) the overlay of (e) GFP and propidium iodide, and (f) YFP and DAPI. Pictures were taken (a,c,e) 14 and (b,d,f) 5 das. Scale bars indicate 20 &#x03BC;m.

To investigate the function of ABCE2, we performed a co-immunoprecipitation assay using the ABCE2:YFP protein from a homozygous T_3_ *api7-1 35S_pro_:ABCE2:YFP* line, which was phenotypically wild-type, confirming that the fusion protein is functional (Fig. S12a–c). We checked the purification of the fusion protein by western blotting using an anti-GFP antibody (Fig. S12d–f). Using LC-ESI-MS/MS, we identified 20 putative interactors of ABCE2, of which 13 participate in translation (6 subunits of the eIF3 complex, eIF5B, RPL3B, and ROTAMASE CYP 1 [ROC1]) or in its regulation (At5g58410,EVOLUTIONARILY CONSERVED C-TERMINAL REGION 2 [ECT2], ILITYHIA [ILA], and REGULATORY-ASSOCIATED PROTEIN OF TOR 1 [RAPTOR1] or RAPTOR2), and two others had previously been shown to interact with ABCE2 orthologs (At2g20830, and EXPORTIN 1A [XPO1A] or XPO1B). The functions of the remaining 5 proteins that co-immunoprecipitated with ABCE2 are unclear and these proteins were therefore set aside for future characterization (Fig. 6; Fig. S13; Tables S7, S8; Data Set 1).

**Fig. 6.**
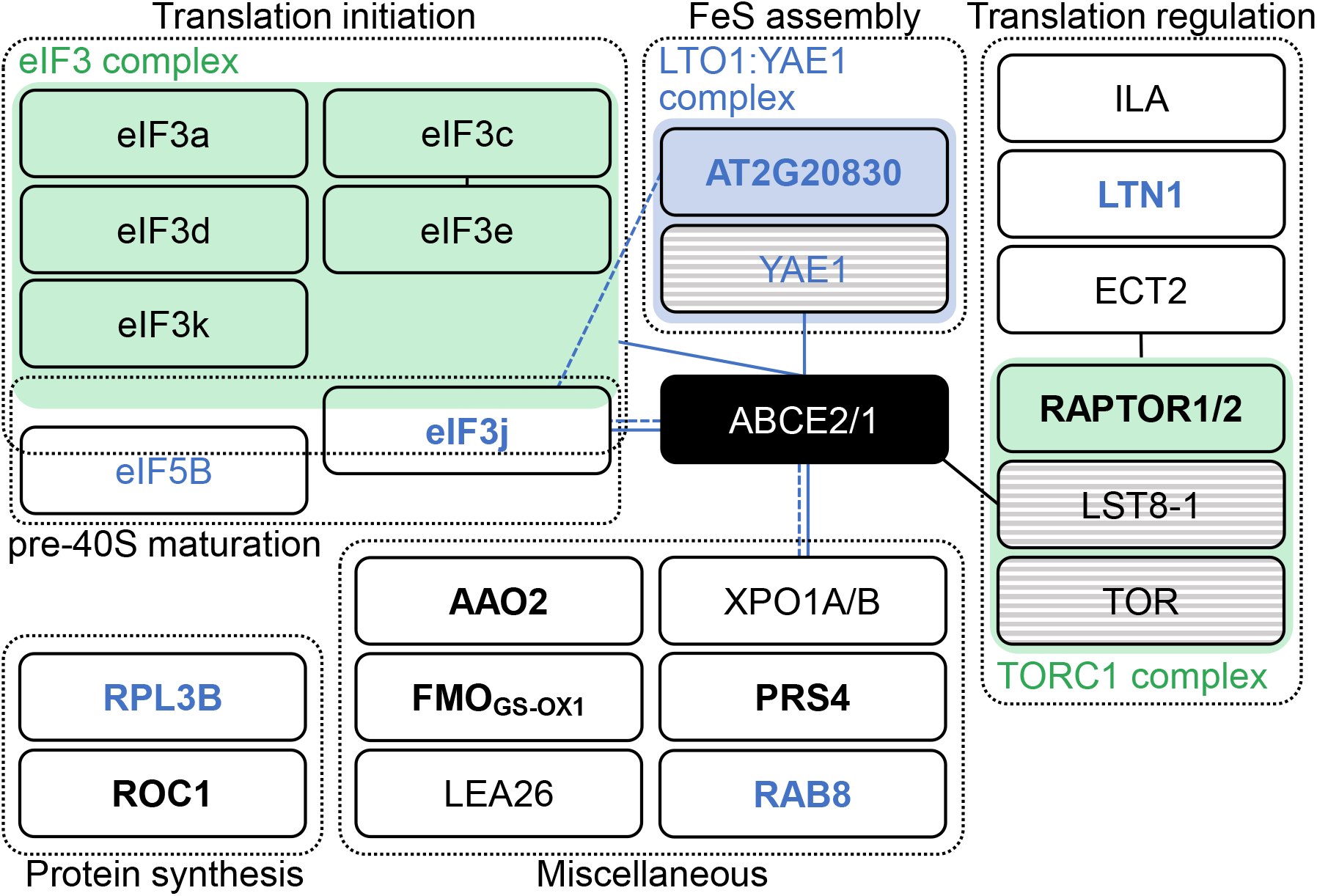
Proteins identified in an ABCE2:YFP co-immunoprecipitation assay. Proteins were grouped within dashed boxes according to their annotated functions for Arabidopsis (names in black letters) or orthologous (names in blue letters) proteins. Green and blue boxes represent complexes that have been described in Arabidopsis and other species, respectively. Proteins in striped boxes were not identified in our assay but have been included in this figure because they are known to belong to a given complex. Continuous and dashed lines connecting boxes indicate physical and genetic interactions described elsewhere for Arabidopsis (black) or other species (blue), respectively. For references, see Table S8. Names in bold and plain letters indicate proteins unique to or enriched in ABCE2:YFP samples, respectively.

At2g20830 encodes a folic acid binding/transferase that shares 30.2% and 26.7% identity with human and *Saccharomyces cerevisiae* Lto1 (named after “required for biogenesis of the large ribosomal subunit and initiation of translation in oxygen”), respectively (human and *S. cerevisiae* Lto1 proteins share 27.8% identity). Lto1, together with Yae1, constitute an essential complex for FeS cluster assembly on ABCE1 (Zhai *et al*.,2014; Paul *et al*., 2015; Zhu *et al*., 2020; Prusty *et al*., 2021). Despite the observation that At2g20830 protein was predicted to localize to mitochondria, the conservation level of this protein with its yeast and human Lto1 orthologs prompted us to consider At2g20830 an ABCE2 interactor. Indeed, At2g20830 may be necessary for FeS cluster assembly on ABCE2.

Our co-immunoprecipitation assay suggested that Arabidopsis ABCE2 interacts with 6 of the 13 eIF3 subunits: eIF3a, c, d, e, k, and j. In *S. cerevisiae*, those interactions have been related to the presence of the ABCE1 protein in the 40S subunit after ribosome dissociation, until late steps of initiation of a new cycle of translation (Heuer *et al*., 2017;Mancera-Martínez *et al*., 2017; Kratzat *et al*., 2021). Interestingly, the interaction between the non-stoichiometric subunit eIF3j and ABCE1 also occurs in humans and *S. cerevisiae*. In these species, eIF3j acts as an accessory factor for ABCE1-mediated ribosome dissociation (Young & Guydosh, 2019; Kratzat *et al*., 2021), a function that seems to be conserved in Arabidopsis.

To corroborate and extend the list of interactions between ABCE2 and components of the translation machinery, we performed a TAP assay of a GSRhino-TAP-tagged ABCE2 bait, obtained from cell suspension cultures, and identified its putative interactors by nano LC-MS/MS (Data Set 2). We found that 81 proteins co-purified with ABCE2, of which 28 were ribosomal proteins (Data Set 2e).

### *api7-1* mutation perturbs auxin metabolism

To gain insight into the biological processes affected in the *api7-1* mutant, we performed an RNA-seq analysis of L*er* and *api7-1* shoots collected 14 das. We identified 3218 downregulated and 2135 upregulated genes in the *api7-1* mutant (Data Set 3a). A GO enrichment analysis performed separately for down- and upregulated genes showed that the downregulated genes were mainly related to responses to abiotic and biotic stresses and protein post-translational modifications. In contrast, upregulated genes grouped into more diverse Biological Process terms (Data Set 3b,c). Among them, we found three terms related to auxin (response to auxin [GO:0009733], auxin-activated signaling pathway [GO:0009734], and auxin polar transport [GO:0009926]).

We observed that four out of the six genes that participate in the main auxin biosynthesis pathway in shoots were upregulated. They included two of the three genes encoding enzymes that convert tryptophan (Trp) into indole-3-pyruvic acid (IPyA), *TRYPTOPHAN AMINOTRANSFERASE OF ARABIDOPSIS 1* (*TAA1*), and *TAA1-RELATED 2 (TAR2)*, and two *YUCCA* genes *(YUC2* and *YUC6)* encoding enzymes that turn IPyA into indole-3-acetic acid (IAA) (Cheng *et al*., 2007; Casanova-Sáez *et al*., 2021;Kneuper *et al*., 2021). However, the expression of two genes involved in a secondary pathway for IAA biosynthesis, *CYTOCHROME P450, FAMILY 79, SUBFAMILY B, POLYPEPTIDE 2 (CYP79B2)* and *IAMHYDROLASE12 (IAMH2)* were downregulated. We also found that four genes involved in auxin inactivation, *IAA CARBOXYLMETHYLTRANSFERASE 1 (IAMT1), GRETCHEN HAGEN 3.17 (GH3.17), DIOXYGENASE FOR AUXIN OXIDATION 2 (DAO2)*, and *UDP-glycosyltransferase 76E5*(*UGT76E5*) were upregulated, and that three genes involved in auxin reactivation, *IAA-LEUCINE RESISTANT (ILR)-LIKE 2 (ILL2), ILL3*, and *ILL4* (Casanova-Sáez *et al*., 2021; Mateo-Bonmatí *et al*., 2021), were downregulated, probably in response to high auxin levels (Fig. 7a). In this manner, our transcriptional data point to an increase in auxin biosynthesis in *api7-1* shoots, which might be partially or fully compensated by reducing the synthesis rate in secondary pathways, and by inactivating and preventing the reactivation of IAA.

**Fig. 7.**
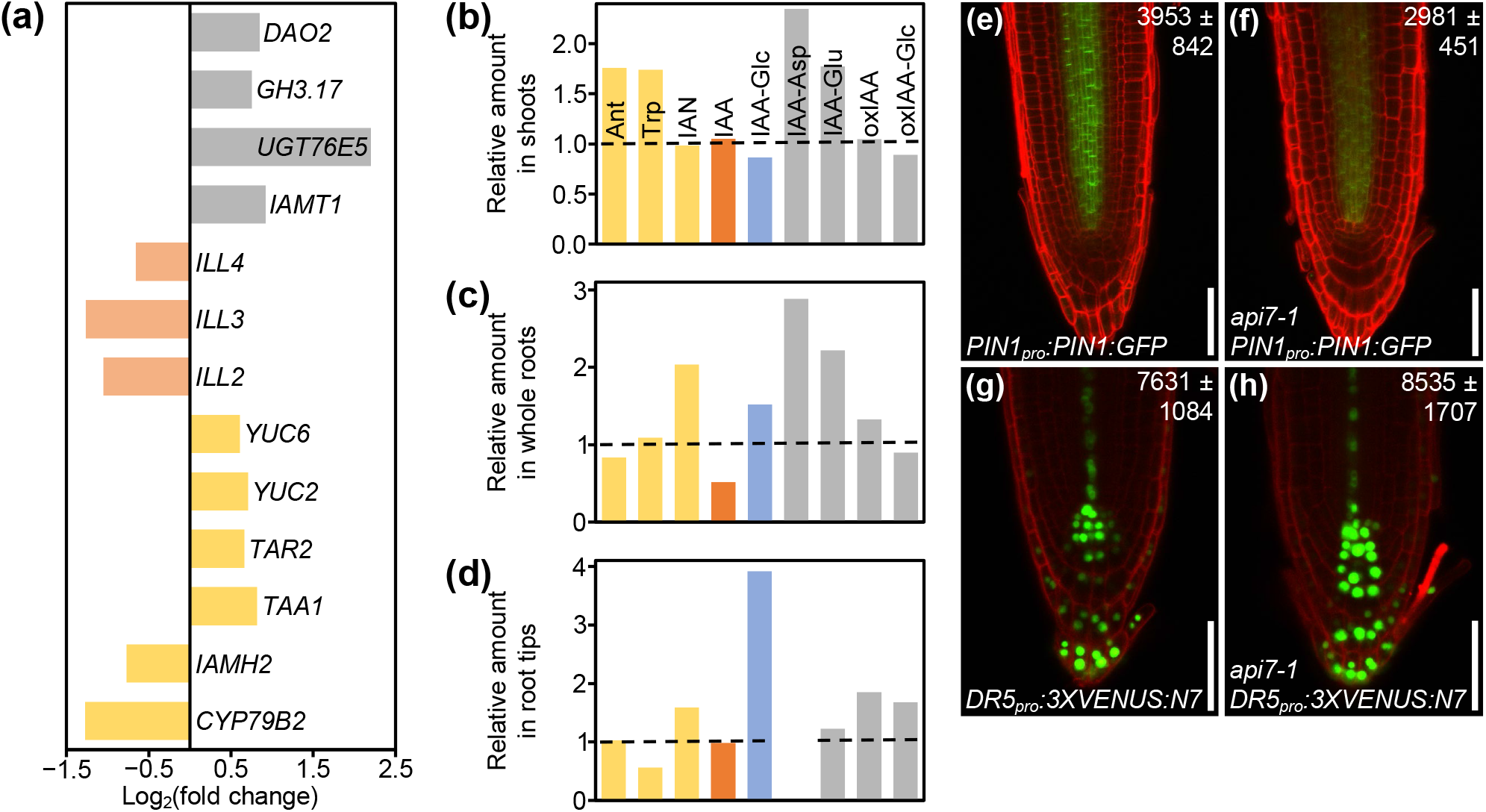
Auxin metabolism, transport and signaling are altered in *api7-1* plants. (a) Expression levels of genes related to auxin metabolism (biosynthesis, yellow; activation, pale orange; storage and catabolism, grey) in *api7-1* shoots 14 das. Values are shown as the binary logarithm of the foldchange between *api7-1* and *Ler* mean reads. Mean reads were calculated from three biological replicates. (b–d) Relative amounts of some IAA precursors (yellow), IAA (orange), the IAA storage molecule IAA-Glc (blue), and IAA catabolites (grey) in *api7-1* (b) shoots, (c) whole roots, and (d) root tips 9 das. IAA-Asp was not detected in root tips. The mean amounts of each metabolite in L*er* were used as the reference value (dashed lines; see Fig. S14). Mean amounts were calculated from four biological replicates. (e–h) Visualization of the expression of reporter transgenes for auxin (e,f) transport and (g,h) perception, in (e,g) wild-type and (f,h) *api7-1* roots. Cell walls were stained with propidium iodide. Values indicate average fluorescence intensities ±standard deviation from (e,f) GFP and (g,h) VENUS, which are significantly different from the wild type in a Student’s *t* test [(e,f) *P <* 0.001, n = 25; (g,h) *P <* 0.05, n = 27]. Pictures were taken 5 das. Scale bars indicate 50 μm.

To directly assess our hypothesis, we checked the content of IAA, the main auxin in most plants, as well as some of its precursors and inactive forms in *api7-1* and L*er* shoots, whole roots and root tips. We found a similar trend within the three tissues: an increase in IAA catabolism, as suggested by the RNA-seq results, and an accumulation of its precursors, when compared to L*er* tissues (Fig. 7b–d; Fig. S14). The inactivation of IAA in *api7-1* shoots and whole roots mainly occurs through glutamate (IAA-Glu) and aspartate (IAA-Asp) conjugation, while in root tips occurs through IAA oxidation (oxIAA), and subsequent glycosylation (oxIAA-glc). *api7-1* shoots accumulate anthranilate (Ant), a substrate for Trp biosynthesis, and Trp itself, the main precursor for IAA biosynthesis. Whole roots and root tips accumulate indole-3-acetonitrile (IAN), another IAA precursor, and store the inactive glycosylated IAA (IAA-glc). The IAA levels were normal in shoots and root tips, suggesting that auxin homeostasis is maintained in *api7-1*. However, the IAA levels in whole roots were decreased by almost 50%, maybe due to its high inactivation levels. Trp levels were low in root tips, suggesting that it might be converted to IAN, which is overaccumulated, or to IAA, which seems to be stored and catabolized to maintain its normal levels.

In agreement with the reduced levels of IAA in *api7-1* whole roots, the levels of a fusion protein between the auxin exporter PIN-FORMED1 (PIN1) and GFP (PIN1:GFP) in *api7-1 PIN1_pro_:PIN1:GFP* roots were lower than in L*er* roots (Fig. 7e,f). In addition, we observed that the expression of the synthetic auxin-responsive promoter *DR5* in *api7-1 DR5_pro_:3XVENUS:N7* root tips, measured as the fluorescence intensity of 3XVENUS:N7, was slightly increased in comparison to L*er DR5_pro_:3XVENUS:N7* root tips, indicating that auxin signaling might be also altered in *api7-1* (Fig. 7g,h).

### Genes related to iron homeostasis are deregulated in *api7-1* plants

Interestingly, we also found in our RNA-seq assay that iron ion homeostasis and transport (GO:0055072 and GO:0006826), and response to iron and sulfur ion starvation (GO:0010106 and GO:0010438) terms were among the most enriched in the analysis of upregulated genes (Data Set 3b). For instance, genes related to iron uptake, such as *IRON-REGULATED TRANSPORTER 1 (IRT1)* and *FERRIC CHELATE REDUCTASE DEFECTIVE 1* (*FRD1*) (Eide *et al*., 1996; Robinson *et al*., 1999), or to iron mobility, such as *NATURAL RESISTANCE-ASSOCIATED MACROPHAGE PROTEIN 3* (*NRAMP3*) and *NRAMP6* (Lanquar *et al*., 2005; Li *et al*., 2019), and several genes encoding transcription factors induced by iron and sulfur deficiencies were upregulated in the *api7-1* mutant (Fig. S15). These pathways might be activated in *api7-1* plants to provide iron and sulfur for FeS cluster biogenesis, probably to compensate for the depletion in ABCE2 protein. Indeed, the gene that encodes the Arabidopsis NEET protein (termed after its conserved Asn-Glu-Glu-Thr sequence near its C-terminus) (Colca *et al*., 2004), which participates in FeS cluster transference during its biogenesis (Nechushtai *et al*., 2012; Zandalinas *et al*., 2020), was also upregulated (Fig. S15).

Consequently, the iron content in *api7-1* cells might be higher than in the wild type, and might be inducing the formation of reactive oxygen species (ROS), as occurs in mutants affected in free iron storage (Briat *et al*., 2010). In agreement with this assumption, several terms related to oxidative stress responses were also enriched. Specifically, we found that *FERRITIN 2 (FER2)* and *FER3*, which encode iron storage proteins in response to high iron levels to avoid oxidative damage (Briat *et al*., 2010; Reyt *et al*., 2015), were upregulated (Fig. S15). In addition, previous studies have shown that ROS prevent FeS cluster assembly into ABCE proteins, which is necessary for their activity in ribosome recycling (Alhebshi *et al*., 2012; Sudmant *et al*., 2018; Zhu *et al*., 2020). In this manner, *api7-1* plants might experience a positive feedback loop where a response to iron starvation due to reduced activity of ABCE2 increases iron levels, inducing the production of ROS which, in turn, further disturbs ABCE2 activity. Nevertheless, further studies are needed to ascertain a potential relation among ABCE2 activity, iron homeostasis, and oxidative stress, which were beyond the scope of this work.

## DISCUSSION

### Plant ABCE proteins participate in translation in a cross-kingdom conserved manner

In this work, we studied Arabidopsis ABCE2, one of the most conserved proteins among archaea and eukaryotes (Hopfner, 2012). Archaea, fungi, and animal ABCE proteins dissociate cytoplasmic ribosomes into their 30S/40S and 50S/60S subunits at different translation-related events (Nürenberg-Goloub & Tampé, 2019). After ribosome dissociation, an ABCE escorts the 30S/40S subunit until the late steps of translation initiation, preventing premature joining of the 50S/60S subunit into the preinitiation complex (Heuer *et al*., 2017; Nürenberg-Goloub *et al*., 2020).

The crosslinking performed on the tissue used for the ABCE2 co-immunoprecipitation assay did not allow us to discern direct from indirect ABCE2 interactors. However, the interactions with XPO1A/B, eIF3j, and the protein encoded by At2g20830 are very likely to be direct, in agreement with previous studies in non-plant species (Kirli *et al*., 2015; Paul *et al*., 2015; Young & Guydosh, 2019; Kratzat *et al*., 2021). In contrast, the interactions observed with other eIF3 subunits, RPL3B, ROC1, ECT2, ILA, RAPTOR1/2, and the protein encoded by At5g58410, which is annotated as E3 ubiquitin-protein ligase listerin (LTN1; UniProt code: Q9FGI1) might occur indirectly as they are part of or interact with the translation machinery (Coaker *et al*., 2006; Shao *et al*., 2013; Kashima *et al*., 2014; Sesma *et al*., 2017; Wang *et al*., 2017; Arribas-Hernández *et al*., 2018; Faus *et al*., 2018; Izquierdo *et al*., 2018; Scutenaire *et al*., 2018; Wei *et al*., 2018). However, the interaction between ABCE2 and eIF5B, which does not seem to occur in *S. cerevisiae* and mammals (Heuer *et al*., 2017; Mancera-Martínez *et al*., 2017), will require further exploration.

Further supporting a role for ABCE2 in translation, we observed a synergistic interaction in the *api7-1 as2-1* double mutant, which show radial leaves, as previously described for double mutant combinations of loss-of-function alleles of *AS1* or *AS2* and other components of the translation machinery (Pinon *et al*., 2008; Yao *et al*., 2008;Horiguchi *et al*., 2011; Moschopoulos *et al*., 2012; Casanova-Sáez *et al*., 2014; Mateo-Bonmatí *et al*., 2015). In this manner, our results strongly suggest that Arabidopsis and, by extension, all plant ABCEs, probably dissociate cytoplasmic ribosomes, as has been reported for species of other kingdoms (Nürenberg-Goloub & Tampé, 2019). In addition, previous works also support a conserved role for the Arabidopsis ABCE2 and human ABCE1 proteins as suppressors of RNA silencing (Braz *et al*., 2004; Sarmiento et al., 2006;Kärblane *et al*., 2015; Mõttus *et al*., 2020). However, we did not find any ABCE2 interactor potentially involved in this process, nor any enriched ontology term related to gene silencing in our RNA-seq assay. This might be due to the need for a cellular environment that triggers RNA silencing and exposes this novel function of ABCE proteins. Further research will help to assess a potential relationship between ribosome recycling and RNA silencing.

### The developmental defects of the *api7-1* mutant have different causes

The essential function of ABCEs has been confirmed in several species: null alleles of *ABCE* genes in all studied organisms are lethal, while hypomorphic alleles cause severe growth aberrations (Navarro-Quiles *et al*., 2018). In this work, we describe the first hypomorphic and null alleles of the Arabidopsis *ABCE2* gene, *api7-1* and *api7-2*, respectively. We showed that the *api7-2* mutation is lethal and that *api7-1* plants share developmental defects with other mutants affected in genes encoding ribosomal proteins or ribosome biogenesis factors. These phenotypic traits include an aberrant leaf venation pattern (Horiguchi *et al*., 2011), as is the case for the *api7-1* mutant. Indeed, a mutant allele of *SIMPLE LEAF3*, the *Cardamine hirsuta ABCE* ortholog, also causes venation pattern defects which may be related to an aberrant auxin transport and signaling (Kougioumoutzi *et al*., 2013).

In agreement with the involvement of local auxin biosynthesis, polar transport and signalling in vascular development (Verna *et al*., 2019; Kneuper *et al*., 2021), we observed that auxin metabolism and auxin-induced genes were upregulated in the *api7-1* mutant. In addition, a previous study found that the IAA content in *api7-1* seedlings was slightly reduced when compared to L*er* (Pěnčík *et al*., 2018). In our experimental conditions, IAA levels in *api7-1* shoots and root tips were normal, but reduced in whole roots. However, the general accumulation of IAA precursors and catabolites in *api7-1* seedlings suggests that, despite auxin metabolism in *api7-1* is perturbed, auxin homeostasis is maintained through different compensation mechanisms, like occurs in other mutants affected in IAA metabolism (Mellor *et al*., 2016; Porco *et al*., 2016; Zhang *et al*., 2016). In this sense, the altered levels of IAA precursors and catabolites, and the deregulation of auxin signalling might contribute to the aberrant phenotype of *api7-1* plants. Our transcriptomic results also point to the deregulation of additional biological pathways as potential contributors to the *api7-1* phenotype: one of them might be an increased production of ROS, caused by a potential deregulation of iron and sulfur homeostasis.

### The Arabidopsis ABCE1 and ABCE2 proteins are functionally redundant

Arabidopsis has two ABCE paralogs, *ABCE1* and *ABCE2* (Sánchez-Fernández *et al*., 2001;Verrier *et al*., 2008). In agreement with previous literature (Braz *et al*., 2004; Sarmiento *et al*., 2006), we observed that *ABCE1* expression levels are low in all studied organs and throughout development, in contrast to the high expression of *ABCE2*. We also showed that *ABCE1* is unable to complement *ABCE2* dysfunction in *api7-1* rosettes *per se*.

However, the wild-type phenotype of *api7-1* flowers, where we found the highest expression levels of *ABCE1*, and the ability of *ABCE2_pro_:ABCE1* and *35S_pro_:ABCE1* to complement the *api7-1* mutant phenotype, indicate that the ABCE1 protein is functional and that it may contribute to translation in the reproductive tissues of wild-type plants. In addition, our phylogenetic analysis showed that the *ABCE* duplication event occurred early during the evolution of Brassicaceae, and that at least five species from this clade conserved an *ABCE1* gene that evolved more rapidly than its *ABCE2* paralog, suggesting that *ABCE2* conserved the ancestral function, whereas *ABCE1* underwent hypofunctionalization (Veitia,2017).

ABCE proteins are encoded by a single gene in most species, and they are essential for archaea and eukaryotes (Navarro-Quiles *et al*., 2018). Due to their importance, the molecular mechanisms by which they participate in ribosome recycling have been deeply studied, and remain a subject of intense research (Heuer *et al*., 2017; Mancera-Martínez *et al*., 2017; Nürenberg-Goloub *et al*., 2018; Nürenberg-Goloub *et al*., 2020; Kratzat *et al*.,2021). Nevertheless, the biological consequences of ABCE depletion or disruption are poorly understood in all organisms. In this sense, future research linking the molecular function of ABCEs with the phenotypic output of their dysfunction will contribute to determining the pathways through which translation modulates development, as we show here with the isolation and study of the hypomorphic and viable *api7-1* allele of the Arabidopsis *ABCE2* gene.

## Supporting information

Supporting Data Set 1

Supporting Data Set 2

Supporting Data Set 3

## ACKNOWLEDGEMENTS

We thank J. Castelló, J.M. Serrano, and M.J. Ñíguez for their excellent technical assistance, and M. Sendra-Ortolà and I.C. Pomares-Bri for helping in the phenotypic analysis of *api7-1* and some gene constructs.

## AUTHOR CONTRIBUTIONS

J.L.M. conceived, designed, and supervised the research, provided resources, and obtained funding. Several experiments were codesigned by C.N.-Q., E.M.-B., and J.L.M. C.N.-Q. performed most of the experiments. E.M.-B. obtained the *ABCE2_pro_:ABCE2* and *35S_pro_:ABCE2:GFP* transgenes, and contributed to the phenotypic analysis of *api7-1*. E.M.-B. and H.C. obtained the *api7-1 as* double mutants. C.N.-Q. and H.C. performed the phylogenetic analysis. H.C. and A.M.L. screened the Micol collection of leaf mutants for abnormal leaf venation patterns. P.R. performed preliminary morphometric analysis of cells and venation from *api7-1* leaves. J.Š. and K.L. performed the IAA metabolite profiling. Y.F. and V.R. performed the TAP assay. M.R.P., H.C., and E.M-B. performed the mapping and cloning of the *api7-1* mutation. C.N.-Q. and J.L.M. wrote the manuscript. All authors revised and approved the manuscript.

## DATA AVAILABILITY

The raw data from genome resequencing and RNA-seq were deposited in the Sequence Read Archive (https://www.ncbi.nlm.nih.gov/sra/) database under accession numbers SRP065876 and PRJNA719000, respectively.

## SUPPORTING INFORMATION

Additional supporting information may be found in the online version of this article.

**Fig. S1** Primary root length, inflorescence and silique morphological phenotypes, and pigment content in leaves of L*er*, *api7-1*, and *api7-1 ABCE2_pro_:ABCE2* plants.

**Fig. S2** Leaf cell phenotypes of L*er*, *api7-1*, and *api7-1 ABCE2_pro_:ABCE2* plants.

**Fig. S3** Some details of the vascular phenotype of first- and third-node leaves from L*er* and *api7-1* plants.

**Fig. S4** Venation pattern of *api7-1* cotyledons, cauline leaves, sepals, and petals.

**Fig. S5** Vascularization in *api7-1* leaf primordia.

**Fig. S6** Genetic interactions of *api7-1* with *as1-1* and *as2-1*.

**Fig. S7** Sequence conservation among ABCE orthologs.

**Fig. S8** *api7-2* is a lethal allele of *ABCE2*.

**Fig. S9** Phylogenetic analysis of some Rosidae *ABCE* genes.

**Fig. S10** *ABCE1* and *ABCE2* expression analyses.

**Fig. S11** The *35S_pro_:ABCE1* transgene restores the wild-type phenotype in *api7-1* plants.

**Fig. S12** The *35S_pro_:ABCE2:YFP* transgene fully restores the wild-type phenotype in *api7-1* plants.

**Fig. S13** Amino acid sequences of proteins identified by LC-ESI-MS/MS in co-immunoprecipitated ABCE2:YFP protein.

**Fig. S14** Tissue profiling of IAA metabolites in *api7-1* seedlings.

**Fig. S15** Expression levels of some genes deregulated in *api7-1* plants.

**Table S1** Primer sets used in this work.

**Table S2** Excitation and detection parameters of fluorophores.

**Table S3** Quality control summary of the RNA-seq assay.

**Table S4** NCBI accession numbers of the sequences used for phylogenetic analysis.

**Table S5** Morphometry of the leaf venation pattern of the *api7-1* mutant.

**Table S6** Mutations identified in the *api7-1* candidate interval.

**Table S7** ABCE2 interactors identified in a co-immunoprecipitation assay.

**Table S8** Conservation level and described functions of putative ABCE2 interactors.

**Data Set 1** Proteins identified in an ABCE2:YFP co-immunoprecipitation assay.

**Data Set 2** Proteins identified in a GSRhino-TAP-tagged ABCE2 fusion tandem affinity purification assay.

**Data Set 3** Genes deregulated in an RNA-seq analysis of *api7-1* plants and gene ontology term enrichment analysis.

## REFERENCES

Alhebshi A, Sideri TC, Holland SL, Avery SV. 2012. The essential iron-sulfur protein Rli1 is an important target accounting for inhibition of cell growth by reactive oxygen species. Molecular Biology of the Cell 23: 3582–3590.

Altschul SF, Madden TL, Schäffer AA, Zhang J, Zhang Z, Miller W, Lipman DJ. 1997. Gapped BLAST and PSI-BLAST: a new generation of protein database search programs. Nucleic Acids Research 25: 3389–3402.

Arribas-Hernández L, Bressendorff S, Hansen MH, Poulsen C, Erdmann S, Brodersen P. 2018. An m6A-YTH module controls developmental timing and morphogenesis in Arabidopsis. Plant Cell 30: 952–967.

Baima S, Nobili F, Sessa G, Lucchetti S, Ruberti I, Morelli G. 1995. The expression of the *Athb-8* homeobox gene is restricted to provascular cells in *Arabidopsis thaliana*. Development 121: 4171–4182.

Barthelme D, Dinkelaker S, Albers SV, Londei P, Ermler U, Tampé R. 2011. Ribosome recycling depends on a mechanistic link between the FeS cluster domain and a conformational switch of the twin-ATPase ABCE1. Proceedings of the National Academy of Sciences USA 108: 3228–3233.

Barthelme D, Scheele U, Dinkelaker S, Janoschka A, MacMillan F, Albers SV, Driessen AJ, Stagni MS, Bill E, Meyer-Klaucke W, et al. 2007. Structural organization of essential iron-sulfur clusters in the evolutionarily highly conserved ATP-binding cassette protein ABCE1. Journal of Biological Chemistry 282: 14598–14607.

Becker T, Franckenberg S, Wickles S, Shoemaker CJ, Anger AM, Armache JP, Sieber H, Ungewickell C, Berninghausen O, Daberkow I, et al. 2012. Structural basis of highly conserved ribosome recycling in eukaryotes and archaea. Nature 482: 501–506.

Berná G, Robles P, Micol JL. 1999. A mutational analysis of leaf morphogenesis in *Arabidopsis thaliana*. Genetics 152: 729–742.

Bisbal C, Martinand C, Silhol M, Lebleu B, Salehzada T. 1995. Cloning and characterization of a RNase L inhibitor. A new component of the interferon-regulated 2-5A pathway. Journal of Biological Chemistry 270: 13308–13317.

Braz AS, Finnegan J, Waterhouse P, Margis R. 2004. A plant orthologue of RNase L inhibitor (RLI) is induced in plants showing RNA interference. Journal of Molecular Evolution 59: 20–30.

Briat JF, Duc C, Ravet K, Gaymard F. 2010. Ferritins and iron storage in plants. Biochimica et Biophysica Acta 1800: 806–814.

Bühler J, Rishmawi L, Pflugfelder D, Huber G, Scharr H, Hülskamp M, Koornneef M, Schurr U, Jahnke S. 2015. phenoVein-A tool for leaf vein segmentation and analysis. Plant Physiology 169: 2359–2370.

Byrne ME. 2009. A role for the ribosome in development. Trends in Plant Science 14: 512–519.

Candela H, Martínez-Laborda A, Micol JL. 1999. Venation pattern formation in *Arabidopsis thaliana* vegetative leaves. Developmental Biology 205: 205–216.

Casanova-Sáez R, Candela H, Micol JL. 2014. Combined haploinsufficiency and purifying selection drive retention of *RPL36a* paralogs in Arabidopsis. Scientific Reports 4: 4122.

Casanova-Sáez R, Mateo-Bonmatí E, Ljung K. 2021. Auxin metabolism in plants. Cold Spring Harbor Perspectives in Biology 13: a039867.

Clough SJ, Bent AF. 1998. Floral dip: a simplified method for *Agrobacterium-mediated* transformation of Arabidopsis thaliana. Plant Journal 16: 735–743.

Coaker G, Zhu G, Ding Z, Van Doren SR, Staskawicz B. 2006. Eukaryotic cyclophilin as a molecular switch for effector activation. Molecular Microbiology 61: 1485–1496.

Colca JR, McDonald WG, Waldon DJ, Leone JW, Lull JM, Bannow CA, Lund ET, Mathews WR. 2004. Identification of a novel mitochondrial protein (“mitoNEET”) crosslinked specifically by a thiazolidinedione photoprobe. American Journal of Physiology: Endocrinology and Metabolism 286: 252–260.

Curtis MD, Grossniklaus U. 2003. A Gateway cloning vector set for high-throughput functional analysis of genes in planta. Plant Physiology 133: 462–469.

Chen ZQ, Dong J, Ishimura A, Daar I, Hinnebusch AG, Dean M. 2006. The essential vertebrate ABCE1 protein interacts with eukaryotic initiation factors. Journal of Biological Chemistry 281: 7452–7457.

Cheng Y, Dai X, Zhao Y. 2007. Auxin synthesized by the YUCCA flavin monooxygenases is essential for embryogenesis and leaf formation in Arabidopsis. Plant Cell 19: 2430–2439.

Dever TE, Dinman JD, Green R. 2018. Translation elongation and recoding in eukaryotes. Cold Spring Harbor Perspectives in Biology 10: a032649.

Earley KW, Haag JR, Pontes O, Opper K, Juehne T, Song K, Pikaard CS. 2006. Gateway-compatible vectors for plant functional genomics and proteomics. Plant Journal 45: 616–629.

Edgar RC. 2004a. MUSCLE: a multiple sequence alignment method with reduced time and space complexity. BMC Bioinformatics 5: 113.

Edgar RC. 2004b. MUSCLE: multiple sequence alignment with high accuracy and high throughput. Nucleic Acids Research 32: 1792–1797.

Eide D, Broderius M, Fett J, Guerinot ML. 1996. A novel iron-regulated metal transporter from plants identified by functional expression in yeast. Proceedings of the National Academy of Sciences USA 93: 5624–5628.

Faus I, Niñoles R, Kesari V, Llabata P, Tam E, Nebauer SG, Santiago J, Hauser MT, Gadea J. 2018. Arabidopsis ILITHYIA protein is necessary for proper chloroplast biogenesis and root development independent of eIF2α phosphorylation. Journal of Plant Physiology 224-225: 173–182.

García-León M, Iniesto E, Rubio V. 2018. Tandem affinity purification of protein complexes from Arabidopsis cell cultures. Methods in Molecular Biology 1794: 297–309.

Gouridis G, Hetzert B, Kiosze-Becker K, de Boer M, Heinemann H, Nürenberg-Goloub E, Cordes T, Tampé R. 2019. ABCE1 controls ribosome recycling by an asymmetric dynamic conformational equilibrium. Cell Reports 28: 723–734.

Heisler MG, Ohno C, Das P, Sieber P, Reddy GV, Long JA, Meyerowitz EM. 2005. Patterns of auxin transport and gene expression during primordium development revealed by live imaging of the Arabidopsis inflorescence meristem. Current Biology 15: 1899–1911.

Hellen CUT. 2018. Translation termination and ribosome recycling in eukaryotes. Cold Spring Harbor Perspectives in Biology 10: a032656.

Heuer A, Gerovac M, Schmidt C, Trowitzsch S, Preis A, Kötter P, Berninghausen O, Becker T, Beckmann R, Tampé R. 2017. Structure of the 40S-ABCE1 post-splitting complex in ribosome recycling and translation initiation. Nature Structural and Molecular Biology 24: 453–460.

Hooper CM, Castleden IR, Tanz SK, Aryamanesh N, Millar AH. 2017. SUBA4: the interactive data analysis centre for Arabidopsis subcellular protein locations. Nucleic Acids Research 45: 1064–1074.

Hooper CM, Tanz SK, Castleden IR, Vacher MA, Small ID, Millar AH. 2014. SUBAcon: a consensus algorithm for unifying the subcellular localization data of the *Arabidopsis* proteome. Bioinformatics 30: 3356–3364.

Hopfner KP. 2012. Rustless translation. Biological Chemistry 393: 1079–1088.

Horiguchi G, Mollá-Morales A, Pérez-Pérez JM, Kojima K, Robles P, Ponce MR, Micol JL, Tsukaya H. 2011. Differential contributions of ribosomal protein genes to *Arabidopsis thaliana* leaf development. Plant Journal 65: 724–736.

Huang DW, Sherman BT, Lempicki RA. 2009a. Bioinformatics enrichment tools: paths toward the comprehensive functional analysis of large gene lists. Nucleic Acids Research 37: 1–13.

Huang DW, Sherman BT, Lempicki RA. 2009b. Systematic and integrative analysis of large gene lists using DAVID bioinformatics resources. Nature Protocols 4: 44–57.

Izquierdo Y, Kulasekaran S, Benito P, López B, Marcos R, Cascón T, Hamberg M, Castresana C. 2018. Arabidopsis *nonresponding to oxylipins* locus *NOXY7* encodes a yeast GCN1 homolog that mediates noncanonical translation regulation and stress adaptation. Plant, Cell and Environment 41: 1438–1452.

Kärblane K, Gerassimenko J, Nigul L, Piirsoo A, Smialowska A, Vinkel K, Kylsten P, Ekwall K, Swoboda P, Truve E, et al. 2015. ABCE1 is a highly conserved RNA silencing suppressor. PLOS ONE 10: e0116702.

Karcher A, Büttner K, Märtens B, Jansen RP, Hopfner KP. 2005. X-ray structure of RLI, an essential twin cassette ABC ATPase involved in ribosome biogenesis and HIV capsid assembly. Structure 13: 649–659.

Karcher A, Schele A, Hopfner KP. 2008. X-ray structure of the complete ABC enzyme ABCE1 from Pyrococcus abyssi. Journal of Biological Chemistry 283: 7962–7971.

Kashima I, Takahashi M, Hashimoto Y, Sakota E, Nakamura Y, Inada T. 2014. A functional involvement of ABCE1, eukaryotic ribosome recycling factor, in nonstop mRNA decay in *Drosophila melanogaster* cells. Biochimie 106: 10–16.

Kim D, Paggi JM, Park C, Bennett C, Salzberg SL. 2019. Graph-based genome alignment and genotyping with HISAT2 and HISAT-genotype. Nature Biotechnology 37: 907–915.

Kirli K, Karaca S, Dehne HJ, Samwer M, Pan KT, Lenz C, Urlaub H, Görlich D. 2015. A deep proteomics perspective on CRM1-mediated nuclear export and nucleocytoplasmic partitioning. eLIFE 4: e11466.

Kleinboelting N, Huep G, Kloetgen A, Viehoever P, Weisshaar B. 2012. GABI-Kat SimpleSearch: new features of the *Arabidopsis thaliana* T-DNA mutant database. Nucleic Acids Research 40: D1211–1215.

Kneuper I, Teale W, Dawson JE, Tsugeki R, Katifori E, Palme K, Ditengou FA. 2021. Auxin biosynthesis and cellular efflux act together to regulate leaf vein patterning. Journal of Experimental Botany 72: 1151–1165.

Kougioumoutzi E, Cartolano M, Canales C, Dupré M, Bramsiepe J, Vlad D, Rast M, Ioio RD, Tattersall A, Schnittger A, et al. 2013. *SIMPLE LEAF3* encodes a ribosome-associated protein required for leaflet development in Cardamine hirsuta. Plant Journal 73: 533–545.

Kratzat H, Mackens-Kiani T, Ameismeier M, Potocnjak M, Cheng J, Dacheux E, Namane A, Berninghausen O, Herzog F, Fromont-Racine M, et al. 2021. A structural inventory of native ribosomal ABCE1-43S pre-initiation complexes. EMBO Journal 40: e105179.

Kumar S, Stecher G, Li M, Knyaz C, Tamura K. 2018. MEGA X: Molecular Evolutionary Genetics Analysis across computing platforms. Molecular Biology and Evolution 35: 1547–1549.

Lanquar V, Lelièvre F, Bolte S, Hamès C, Alcon C, Neumann D, Vansuyt G, Curie C, Schröder A, Krämer U, et al. 2005. Mobilization of vacuolar iron by AtNRAMP3 and AtNRAMP4 is essential for seed germination on low iron. EMBO Journal 24: 4041–4051.

Li J, Wang Y, Zheng L, Li Y, Zhou X, Li J, Gu D, Xu E, Lu Y, Chen X, et al. 2019. The intracellular transporter AtNRAMP6 is involved in Fe homeostasis in Arabidopsis. Frontiers in Plant Science 10: 1124.

Love MI, Huber W, Anders S. 2014. Moderated estimation of fold change and dispersion for RNA-seq data with DESeq2. Genome Biology 15: 550.

Madeira F, Park YM, Lee J, Buso N, Gur T, Madhusoodanan N, Basutkar P, Tivey ARN, Potter SC, Finn RD, et al. 2019. The EMBL-EBI search and sequence analysis tools APIs in 2019. Nucleic Acids Research 47: 636–641.

Mancera-Martínez E, Brito Querido J, Valasek LS, Simonetti A, Hashem Y. 2017. ABCE1: A special factor that orchestrates translation at the crossroad between recycling and initiation. RNA Biology 14: 1279–1285.

Mateo-Bonmatí E, Casanova-Sáez R, Candela H, Micol JL. 2014. Rapid identification of *angulata* leaf mutations using next-generation sequencing. Planta 240: 1113–1122.

Mateo-Bonmatí E, Casanova-Sáez R, Quesada V, Hricová A, Candela H, Micol JL. 2015. Plastid control of abaxial-adaxial patterning. Scientific Reports 5: 15975.

Mateo-Bonmatí E, Casanova-Sáez R, Šimura J, Ljung K. 2021. Broadening the roles of UDP-glycosyltransferases in auxin homeostasis and plant development. New Phytologist 232: 642–654.

Mateo-Bonmatí E, Esteve-Bruna D, Juan-Vicente L, Nadi R, Candela H, Lozano FM, Ponce MR, Pérez-Pérez JM, Micol JL. 2018. *INCURVATA11* and *CUPULIFORMIS2* are redundant genes that encode epigenetic machinery components in Arabidopsis. Plant Cell 30: 1596–1616.

Mellor N, Band LR, Pěnčík A, Novák O, Rashed A, Holman T, Wilson MH, Voβ U, Bishopp A, King JR, et al. 2016. Dynamic regulation of auxin oxidase and conjugating enzymes *AtDAO1* and *GH3* modulates auxin homeostasis. Proceedings of the National Academy of Sciences USA 113: 11022–11027.

Micol-Ponce R, Sarmiento-Mañús R, Fontcuberta-Cervera S, Cabezas-Fuster A, de Bures A, Sáez-Vásquez J, Ponce MR. 2020. SMALL ORGAN4 is a ribosome biogenesis factor involved in 5.8S ribosomal RNA maturation. Plant Physiology 184: 2022–2039.

Micol-Ponce R, Sarmiento-Mañús R, Ruiz-Bayón A, Montacié C, Sáez-Vasquez J, Ponce MR. 2018. Arabidopsis RIBOSOMAL RNA PROCESSING7 is required for 18S rRNA maturation. Plant Cell 30: 2855–2872.

Moschopoulos A, Derbyshire P, Byrne ME. 2012. The *Arabidopsis* organelle-localized glycyl-tRNA synthetase encoded by *EMBRYO DEFECTIVE DEVELOPMENT1* is required for organ patterning. Journal of Experimental Botany 63: 5233–5243.

Mõttus J, Maiste S, Eek P, Truve E, Sarmiento C. 2020. Mutational analysis of *Arabidopsis thaliana* ABCE2 identifies important motifs for its RNA silencing suppressor function. Plant Biology 23: 21–31.

Navarro-Quiles C, Mateo-Bonmatí E, Micol JL. 2018. ABCE proteins: from molecules to development. Frontiers in Plant Science 9: 1125.

Nechushtai R, Conlan AR, Harir Y, Song L, Yogev O, Eisenberg-Domovich Y, Livnah O, Michaeli D, Rosen R, Ma V, et al. 2012. Characterization of *Arabidopsis* NEET reveals an ancient role for NEET proteins in iron metabolism. Plant Cell 24: 2139–2154.

Novák O, Hényková E, Sairanen I, Kowalczyk M, Pospíšil T, Ljung K. 2012. Tissue-specific profiling of the *Arabidopsis thaliana* auxin metabolome. Plant Journal 72: 523–536.

Nürenberg-Goloub E, Heinemann H, Gerovac M, Tampé R. 2018. Ribosome recycling is coordinated by processive events in two asymmetric ATP sites of ABCE1. Life Science Alliance 1: e201800095.

Nürenberg-Goloub E, Kratzat H, Heinemann H, Heuer A, Kötter P, Berninghausen O, Becker T, Tampé R, Beckmann R. 2020. Molecular analysis of the ribosome recycling factor ABCE1 bound to the 30S post-splitting complex. EMBO Journal 39: e103788.

Nürenberg-Goloub E, Tampé R. 2019. Ribosome recycling in mRNA translation, quality control, and homeostasis. Biological Chemistry 401: 47–61.

Paul VD, Mühlenhoff U, Stümpfig M, Seebacher J, Kugler KG, Renicke C, Taxis C, Gavin AC, Pierik AJ, Lill R. 2015. The deca-GX3 proteins Yae1-Lto1 function as adaptors recruiting the ABC protein Rli1 for iron-sulfur cluster insertion. eLIFE 4: e08231.

Pěnčík A, Casanova-Sáez R, Pilařová V, Žukauskaitė A, Pinto R, Micol JL, Ljung K, Novák O. 2018. Ultra-rapid auxin metabolite profiling for high-throughput mutant screening in Arabidopsis. Journal of Experimental Botany 69: 2569–2579.

Pérez-Pérez JM, Rubio-Díaz S, Dhondt S, Hernández-Romero D, Sánchez-Soriano J, Beemster GT, Ponce MR, Micol JL. 2011. Whole organ, venation and epidermal cell morphological variations are correlated in the leaves of *Arabidopsis* mutants. Plant, Cell and Environment 34: 2200–2211.

Pinon V, Etchells JP, Rossignol P, Collier SA, Arroyo JM, Martienssen RA, Byrne ME. 2008. Three *PIGGYBACK* genes that specifically influence leaf patterning encode ribosomal proteins. Development 135: 1315–1324.

Ponce MR, Quesada V, Micol JL. 1998. Rapid discrimination of sequences flanking and within T-DNA insertions in the *Arabidopsis* genome. Plant Journal 14: 497–501.

Ponce MR, Robles P, Lozano FM, Brotóns MA, Micol JL. 2006. Low-resolution mapping of untagged mutations. Methods in Molecular Biology 323: 105–113.

Ponce MR, Robles P, Micol JL. 1999. High-throughput genetic mapping in *Arabidopsis thaliana*. Molecular and General Genetics 261: 408–415.

Porco S, Pěnčík A, Rashed A, Voβ U, Casanova-Sáez R, Bishopp A, Golebiowska A, Bhosale R, Swarup R, Swarup K, et al. 2016. Dioxygenase-encoding AtDAO1 gene controls IAA oxidation and homeostasis in Arabidopsis. Proceedings of the National Academy of Sciences USA 113: 11016–11021.

Poza-Viejo L, del Olmo I, Crevillén P. 2019. Plant chromatin immunoprecipitation v.2. protocols.io: 25468.

Preis A, Heuer A, Barrio-Garcia C, Hauser A, Eyler DE, Berninghausen O, Green R, Becker T, Beckmann R. 2014. Cryoelectron microscopic structures of eukaryotic translation termination complexes containing eRF1-eRF3 or eRF1-ABCE1. Cell Reports 8: 59–65.

Prusty NR, Camponeschi F, Ciofi-Baffoni S, Banci L. 2021. The human YAE1-ORAOV1 complex of the cytosolic iron-sulfur protein assembly machinery binds a [4Fe-4S] cluster. Inorganica Chimica Acta 518: 120252.

Quesada V, Ponce MR, Micol JL. 2000. Genetic analysis of salt-tolerant mutants in *Arabidopsis thaliana*. Genetics 154: 421–436.

Reyt G, Boudouf S, Boucherez J, Gaymard F, Briat JF. 2015. Iron-and ferritin-dependent reactive oxygen species distribution: impact on *Arabidopsis* root system architecture. Molecular Plant 8: 439–453.

Robinson NJ, Procter CM, Connolly EL, Guerinot ML. 1999. A ferric-chelate reductase for iron uptake from soils. Nature 397: 694–697.

Robles P, Fleury D, Candela H, Cnops G, Alonso-Peral MM, Anami S, Falcone A, Caldana C, Willmitzer L, Ponce MR, et al. 2010. The *RON1/FRY1/SAL1* gene is required for leaf morphogenesis and venation patterning in Arabidopsis. Plant Physiology 152: 1357–1372.

Robles P, Micol JL. 2001. Genome-wide linkage analysis of *Arabidopsis* genes required for leaf development. Molecular Genetics and Genomics 266: 12–19.

Rodnina MV. 2018. Translation in prokaryotes. Cold Spring Harbor Perspectives in Biology 10: a032664.

Rosado A, Li R, van de Ven W, Hsu E, Raikhel NV. 2012. *Arabidopsis* ribosomal proteins control developmental programs through translational regulation of auxin response factors. Proceedings of the National Academy of Sciences USA 109: 19537–19544.

Saitou N, Nei M. 1987. The neighbor-joining method: a new method for reconstructing phylogenetic trees. Molecular Biology and Evolution 4: 406–425.

Sánchez-Fernández R, Davies TG, Coleman JO, Rea PA. 2001. The *Arabidopsis thaliana* ABC protein superfamily, a complete inventory. Journal of Biological Chemistry 276: 30231–30244.

Sarmiento C, Nigul L, Kazantseva J, Buschmann M, Truve E. 2006. AtRLI2 is an endogenous suppressor of RNA silencing. Plant Molecular Biology 61: 153–163.

Scutenaire J, Deragon JM, Jean V, Benhamed M, Raynaud C, Favory JJ, Merret R, Bousquet-Antonelli C. 2018. The YTH domain protein ECT2 is an m6A reader required for normal trichome branching in Arabidopsis. Plant Cell 30: 986–1005.

Schindelin J, Arganda-Carreras I, Frise E, Kaynig V, Longair M, Pietzsch T, Preibisch S, Rueden C, Saalfeld S, Schmid B, et al. 2012. Fiji: an open-source platform for biological-image analysis. Nature Methods 9: 676–682.

Schmittgen TD, Livak KJ. 2008. Analyzing real-time PCR data by the comparative C_T_ method. Nature Protocols 3: 1101–1108.

Sesma A, Castresana C, Castellano MM. 2017. Regulation of translation by TOR, eIF4E and eIF2α in plants: current knowledge, challenges and future perspectives. Frontiers in Plant Science 8: 644.

Shao S, von der Malsburg K, Hegde RS. 2013. Listerin-dependent nascent protein ubiquitination relies on ribosome subunit dissociation. Molecular Cell 50: 637–648.

Shirokikh NE, Preiss T. 2018. Translation initiation by cap-dependent ribosome recruitment: Recent insights and open questions. Wiley Interdisciplinary Reviews: RNA 9: e1473.

Simonetti A, Guca E, Bochler A, Kuhn L, Hashem Y. 2020. Structural insights into the mammalian late-stage initiation complexes. Cell Reports 31: 107497.

Sudmant PH, Lee H, Dominguez D, Heiman M, Burge CB. 2018. Widespread accumulation of ribosome-associated isolated 3’ UTRs in neuronal cell populations of the aging brain. Cell Reports 25: 2447–2456.

Van Leene J, Eeckhout D, Cannoot B, De Winne N, Persiau G, Van De Slijke E, Vercruysse L, Dedecker M, Verkest A, Vandepoele K, et al. 2015. An improved toolbox to unravel the plant cellular machinery by tandem affinity purification of *Arabidopsis* protein complexes. Nature Protocols 10: 169–187.

Veitia RA. 2017. Gene duplicates: agents of robustness or fragility? Trends in Genetics 33: 377–379.

Verna C, Ravichandran SJ, Sawchuk MG, Linh NM, Scarpella E. 2019. Coordination of tissue cell polarity by auxin transport and signaling. eLIFE 8: e51061.

Verrier PJ, Bird D, Burla B, Dassa E, Forestier C, Geisler M, Klein M, Kolukisaoglu Ü, Lee Y, Martinoia E, et al. 2008. Plant ABC proteins - a unified nomenclature and updated inventory. Trends in Plant Science 13: 151–159.

Wang L, Li H, Zhao C, Li S, Kong L, Wu W, Kong W, Liu Y, Wei Y, Zhu JK, et al. 2017. The inhibition of protein translation mediated by AtGCN1 is essential for cold tolerance in *Arabidopsis thaliana*. Plant, Cell and Environment 40: 56–68.

Wei LH, Song P, Wang Y, Lu Z, Tang Q, Yu Q, Xiao Y, Zhang X, Duan HC, Jia G. 2018. The m6A reader ECT2 controls trichome morphology by affecting mRNA stability in Arabidopsis. Plant Cell 30: 968–985.

Weis BL, Kovacevic J, Missbach S, Schleiff E. 2015. Plant-specific features of ribosome biogenesis. Trends in Plant Science 20: 729–740.

Wellburn AR. 1994. The spectral determination of chlorophylls *a* and *b*, as well as total carotenoids, using various solvents with spectrophotometers of different resolution. Journal of Plant Physiology 144: 307–313.

Yao Y, Ling Q, Wang H, Huang H. 2008. Ribosomal proteins promote leaf adaxial identity. Development 135: 1325–1334.

Young DJ, Guydosh NR. 2019. Hcr1/eIF3j is a 60S ribosomal subunit recycling accessory factor *in vivo*. Cell Reports 28: 39–50.

Young DJ, Guydosh NR, Zhang F, Hinnebusch AG, Green R. 2015. Rli1/ABCE1 recycles terminating ribosomes and controls translation reinitiation in 3’UTRs in vivo. Cell 162: 872–884.

Zandalinas SI, Song L, Sengupta S, McInturf SA, Grant DG, Marjault HB, Castro-Guerrero NA, Burks D, Azad RK, Mendoza-Cozatl DG, et al. 2020. Expression of a dominant-negative AtNEET-H89C protein disrupts iron-sulfur metabolism and iron homeostasis in Arabidopsis. Plant Journal 101: 1152–1169.

Zhai C, Li Y, Mascarenhas C, Lin Q, Li K, Vyrides I, Grant CM, Panaretou B. 2014. The function of ORAOV1/LTO1, a gene that is overexpressed frequently in cancer: essential roles in the function and biogenesis of the ribosome. Oncogene 33: 484–494.

Zhang J, Lin JE, Harris C, Campos Mastrotti Pereira F, Wu F, Blakeslee JJ, Peer WA. 2016. DAO1 catalyzes temporal and tissue-specific oxidative inactivation of auxin in *Arabidopsis thaliana*. Proceedings of the National Academy of Sciences USA 113: 11010–11015.

Zhang L, Cai X, Wu J, Liu M, Grob S, Cheng F, Liang J, Cai C, Liu Z, Liu B, et al. 2018. Improved *Brassica rapa* reference genome by single-molecule sequencing and chromosome conformation capture technologies. Horticulture Research 5: 50.

Zhu X, Zhang H, Mendell JT. 2020. Ribosome recycling by ABCE1 links lysosomal function and iron homeostasis to 3’ UTR-directed regulation and nonsense-mediated decay. Cell Reports 32: 107895.

