## Supporting Figures and Tables for "The Arabidopsis ATP-Binding Cassette E protein ABCE2 is a conserved component of the translation machinery"

Supporting Information not included in this file

- Navarro-Quiles et al_Data Set 1.xlsx
- Navarro-Quiles et al_Data Set 2.xlsx
- Navarro-Quiles et al_Data Set 2.xlsx

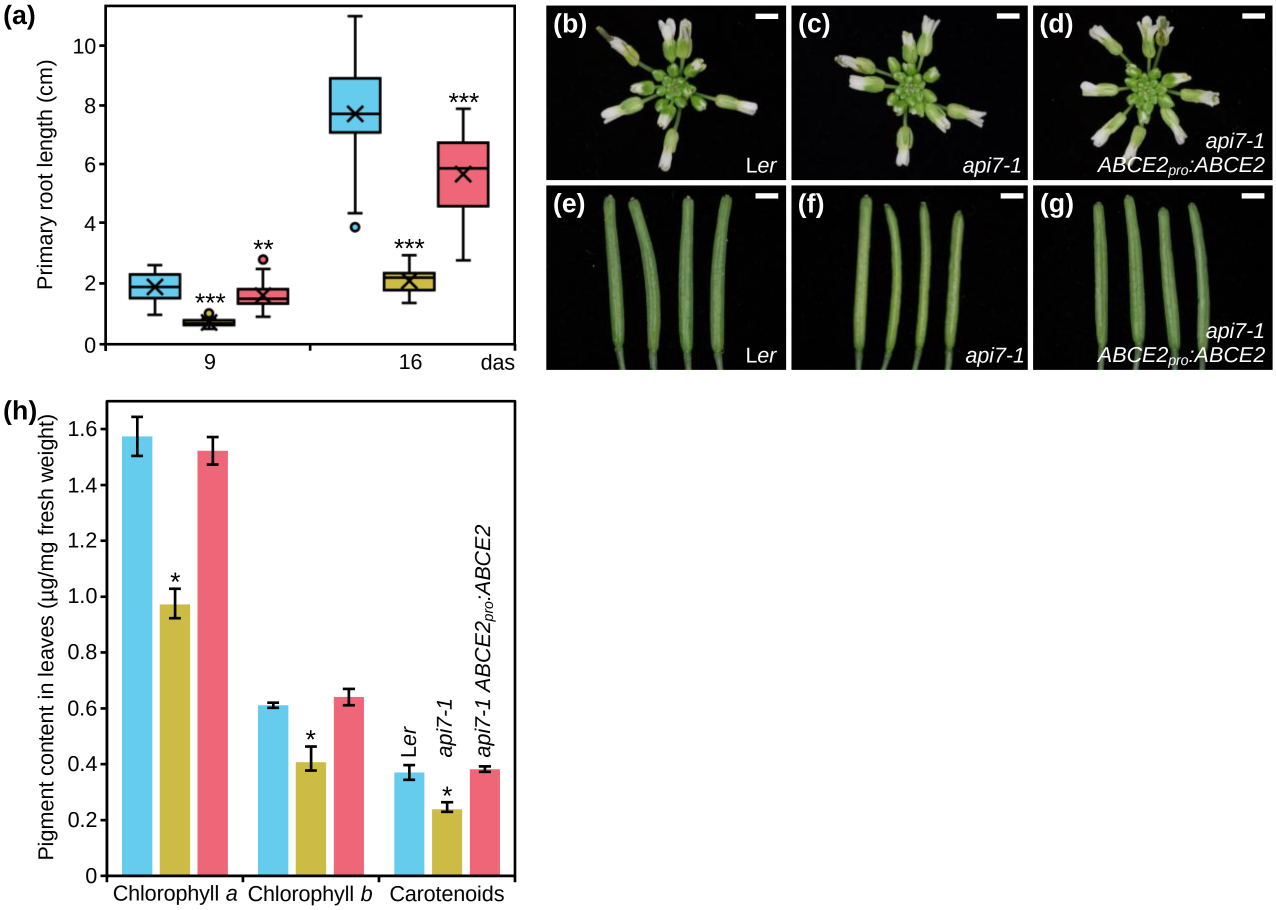


**Fig. S1** Primary root length, inflorescence and silique morphological phenotypes, and pigment content in leaves of L*er*, *api7-1*, and *api7-1 ABCE2$_{pro}$:ABCE2* plants. (a) Primary root growth progression between 9 and 16 das in wild-type L*er*, *api7-1* mutant, and *api7-1 ABCE2$_{pro}$:ABCE2* mutant and transgenic rosettes. Boxes are delimited by the first (Q1, lower hinge) and third (Q3, upper hinge) quartiles. Whiskers represent the most extreme data points that are no more than Q3 + 1.5 × IQR or no less than Q1 − 1.5 × IQR, where the interquartile range (IQR) is Q3 − Q1. ×: Mean. —: Median. ○: Outlier. (b–d) Inflorescences and (e–g) siliques of (b,e) L*er*, (c,f) *api7-1*, and (d,g) *api7-1 ABCE2$_{pro}$:ABCE2* plants. Pictures were taken 40 das. Scale bars indicate 2 mm. (h) Chlorophyll *a* and *b*, and carotenoid content in plants of the genotypes mentioned in (a) collected 16 das. Median values are shown. Error bars represent median absolute deviation. Asterisks indicate a significant difference with L*er* in a (a) Student's *t* test (28 < n < 35) or a (h) Mann-Whitney *U* (n = 5) test (*$P$ < 0.05, **$P$ < 0.01, ***$P$ < 0.001).

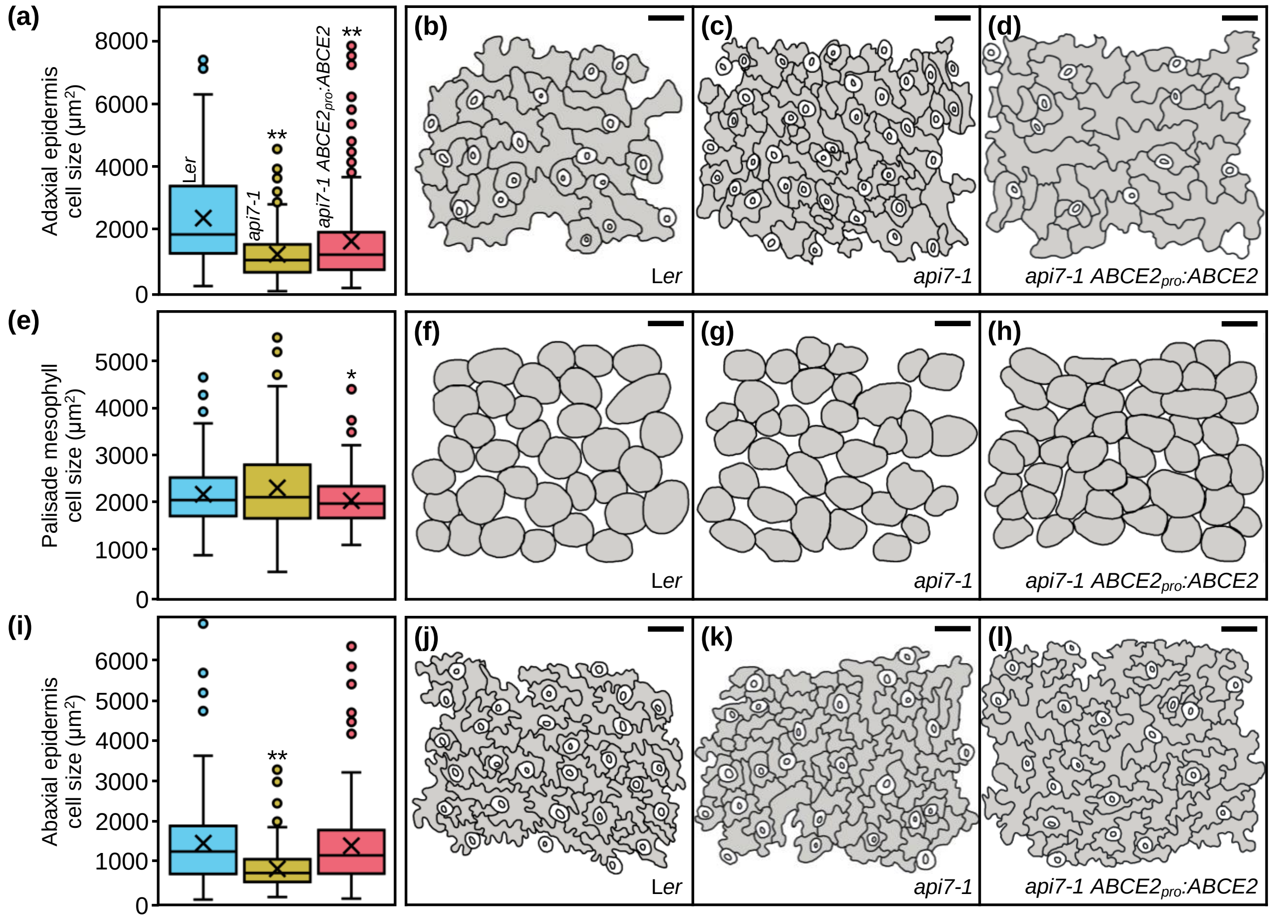


**Fig. S2** Leaf cell phenotypes of L*er*, *api7-1*, and *api7-1 ABCE2$_{pro}$:ABCE2* plants. (a,e,i) Boxplot distributions of cell area in the (a) adaxial epidermis, (e) subepidermal layer of palisade mesophyll, and (i) abaxial epidermis, from first-node leaves collected 21 das. Other details as described in the legend of Fig. S1 for its (a) section. Between 144 and 351 cells were analyzed from at least 4 different samples. Asterisks indicate a significant difference with the L*er* wild-type in a Student's *t* test (*$P$ < 0.05, **$P$ < 0.001). (b–d, f–h, j–l) Representative diagrams of the (b–d) adaxial epidermis, (f–h) subepidermal layer of palisade mesophyll, and (j–l) abaxial epidermis, from (b,f,j) L*er*, (c,g,k) *api7-1*, and (d,h,l) *api7-1 ABCE2$_{pro}$:ABCE2* plants.

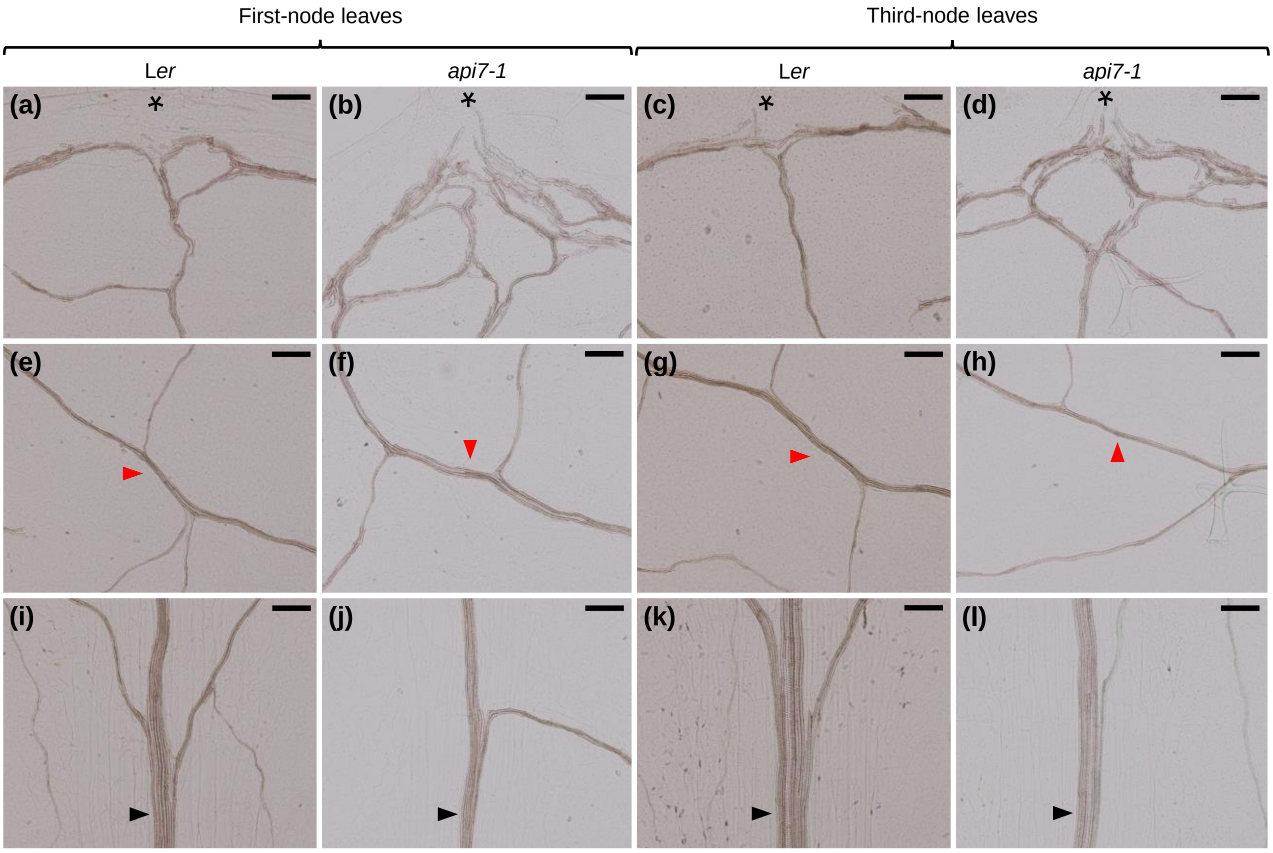


**Fig. S3** Some details of the vascular phenotype of first- and third-node leaves from L*er* and *api7-1* plants. Veins from (a,c,e,g,i,k) L*er* and (b,d,f,h,j,l) *api7-1* (a,b,e,f,i,j) first- and (c,d,g,h,k,l) third-node leaves. (a–d) Venation on the apical region (an asterisk indicates the most apical region) of the lamina, (e–h) a secondary vein (red arrowheads) bifurcating to render tertiary veins, and (i–l) the primary vein (black arrowheads), close to the base of the lamina. We observed 6 first- and third-node leaves from L*er* and 18 from *api7-1* with similar vascular phenotypes to the ones shown. Pictures were taken 21 das. Scale bars indicate 100 µm.

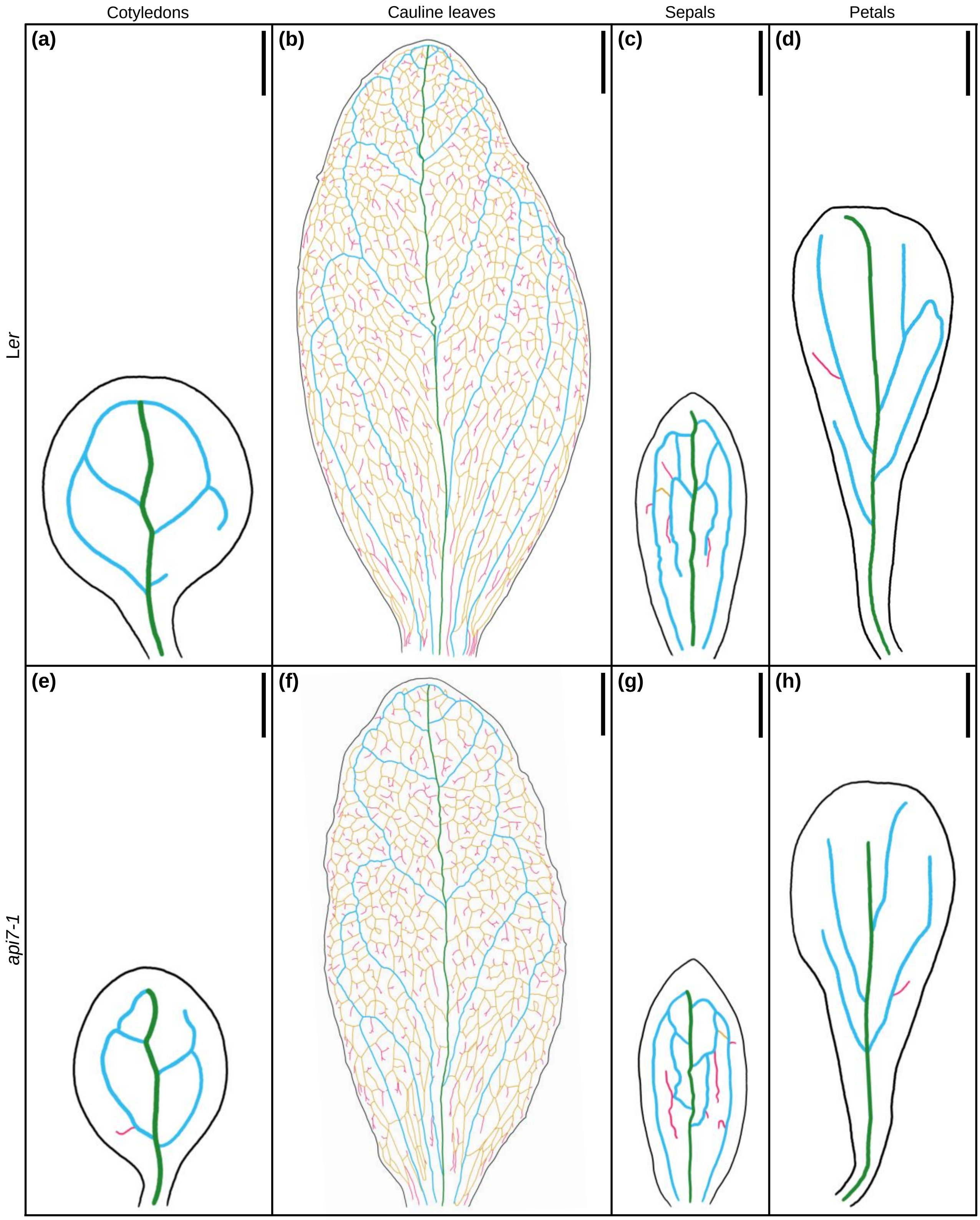


**Fig. S4** Venation pattern of *api7-1* cotyledons, cauline leaves, sepals, and petals. Representative diagrams of (a,e) cotyledons, (b,f) cauline leaves, (c,g) sepals, and (d,h) petals from (a–d) L*er* and (e–h) *api7-1* plants. Margins and veins were drawn as described in Fig. 2. Organs were collected (a,e) 6 and (b–d,f–h) 35 das. Scale bars indicate (a,c–e,g,h) 0.5 and (b,f) 2.5 mm.

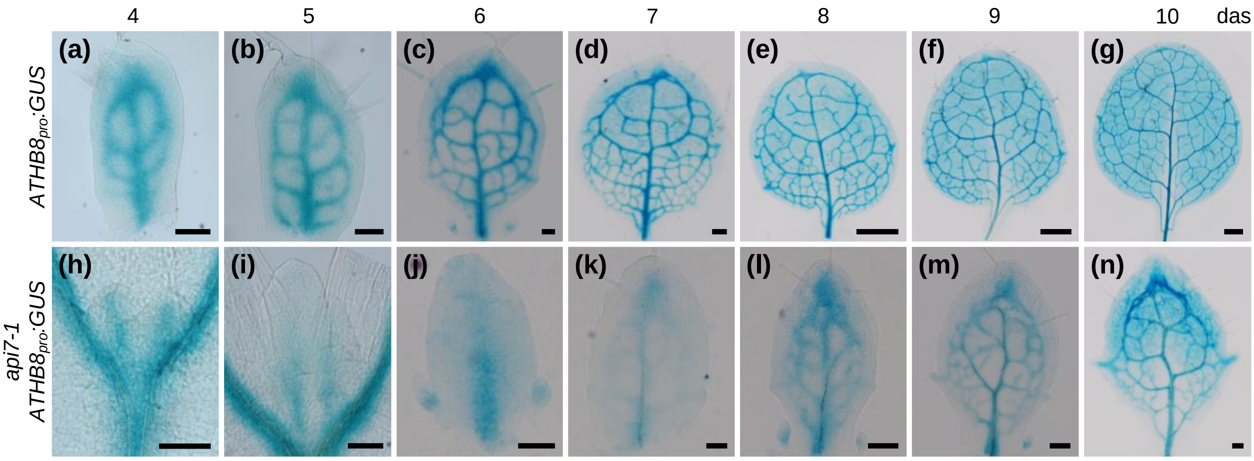


**Fig. S5** Vascularization in *api7-1* leaf primordia. Vascular fate specification is shown as *ATHB8*$_{pro}$*:GUS* activity at expanding first-node leaf primordia in (a–g) L*er* and (h–n) *api7-1* backgrounds. Pictures were taken (a,h) 4, (b,i) 5, (c,j) 6, (c,k) 7, (e,l) 8, (f,m) 9, and (g,n) 10 das. Scale bars indicate (a–c,h–j) 50, (d,k–n) 100, and (e–g) 500 µm.

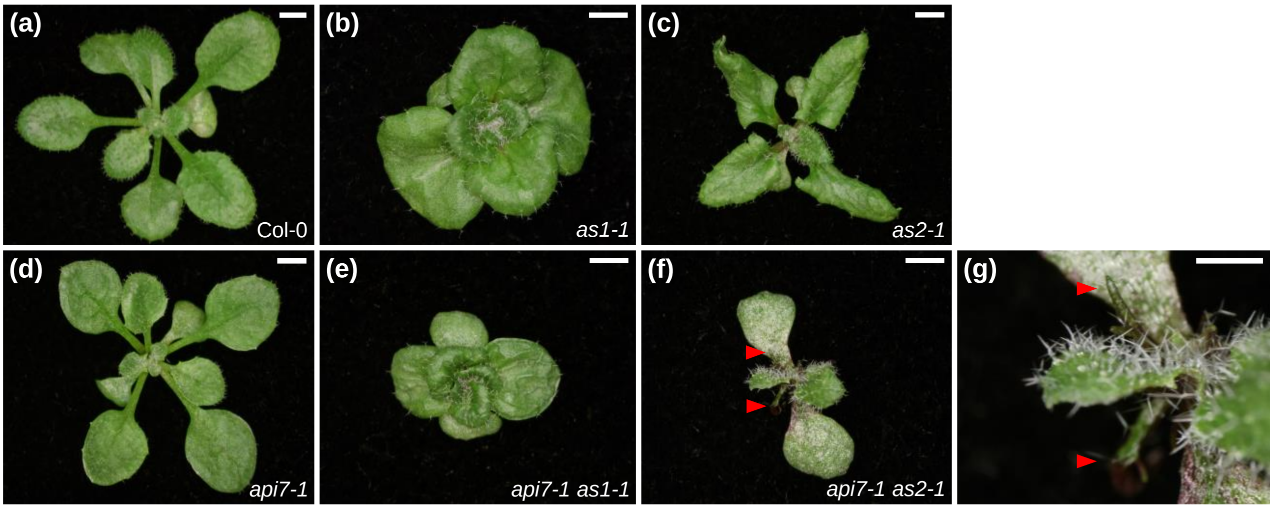


**Fig. S6** Genetic interactions of *api7-1* with *as1-1* and *as2-1*. (a–f) Rosettes from (a) the wild-type L*er*, the (b) *as1-1*, (c) *as2-1*, and (d) *api7-1* single mutants, and the (e) *api7-1 as1-1* and (f) *api7-1 as2-1* double mutants. (g) Close up of (f). Red arrowheads indicate radial leaves. Pictures were taken 16 das. Scale bars indicate (a–f) 2 mm and (g) 1 mm.

```
S. solfataricus     1 ----------------MRVAVINYDFCKPDKCNL-ECINFCPVDRSGGKAIELSEIVKGK
P. furiosus         1 ----------------MRIAVIDYDKCNPDKCGHFLCERVCPVNRMGGEAIIIDEE-NYK
C. elegans          1 MSRKGPLIKNNETDVPLRIAIVEKDRCKPKNCGL-ACKRACPVNRQGKQCIVVEAT-STI
S. cerevisiae       1 -----------MSDKNSRIAIVSADKCKPKKCRQ-ECKRSCPVVKTGKLCIEVTPT-SKI
O. sativa           1 -----------MAERLTRIAIVSEDKCKPKKCRQ-ECKKSCPVVKTGKLCIEVTPA-SKL
S. lycopersicum     1 -----------MSDRLTRIAIVSDDKCKPKKCRQ-ECKKSCPVVKTGKLCIEVTVG-SKI
A. thaliana         1 -----------MADRLTRIAIVSSDRCKPKKCRQ-ECKKSCPVVKTGKLCIEVTVG-SKL
C. hirsuta          1 -----------MADRLTRIAIVSSDRCKPKKCRQ-ECKKSCPVVKTGKLCIEVTVA-SKI
D. melanogaster     1 MSR---RKENEEVDKQTRIAIVSDDKCKPKRCRQ-ECKKTCPVVRMGKLCIEVTPT-SKI
H. sapiens          1 -----------MADKLTRIAIVNHDKCKPKKCRQ-ECKKSCPVVRMGKLCIEVTPQ-SKI
O. cuniculus        1 -----------MADKLTRIAIVNHDKCKPKKCRQ-ECKKSCPVVRMGKLCIEVTPQ-SKI
D. rerio            1 -----------MADKNTRIAIVNHDKCKPKKCRQ-ECKKSCPVVRMGKLCIEVSPQ-SKI
consensus           1            . .. .*.*..  *.*.*..*.. .*...***.. *...*...   ...

S. solfataricus    44 PVIYEETCIGCGICVKKCPYEAISIVNLPDELEGEVIHRYKVNGFKLFGLPTPKNNTILG
P. furiosus        44 PIIQEASCTGCGICVHKCPFKAISIVNLPEQLEEDCVHRYGVNAFVLYRLPVVKEGMVVG
C. elegans         59 SQISEILCIGCGICVKKCPYDAIKIINLPANLANETTHRYSQNSFKLHRLPTPRCGEVLG
S. cerevisiae      48 AFISEILCIGCGICVKKCPFDAIQIINLPTNLEAHVTHRYSANSFKLHRLPTPRPGQVLG
O. sativa          48 AFISEELCIGCGICVKKCPFDAIEIINLPKDLEKDTTHRYGPNTFKLHRLPVPRPGQVLG
S. lycopersicum    48 AFISEELCIGCGICVKKCPFEAIQIINLPKDLDKDTTHRYGPNTFKLHRLPVPRPGQVLG
A. thaliana        48 AFISEELCIGCGICVKKCPFEAIQIINLPRDLEKDTTHRYGANTFKLHRLPVPRPGQVLG
C. hirsuta         48 AFISEELCIGCGICVKKCPFEAIQIINLPRDLEKDTTHRYGPNTFKLHRLPVPRPGQVLG
D. melanogaster    56 ASLSEELCIGCGICVKKCPFEAITIINLPSNLEKHTTHRYSKNSFKLHRLPIPRPGEVLG
H. sapiens         48 AWISETLCIGCGICIKKCPFGALSIVNLPSNLEKETTHRYCANAFKLHRLPIPRPGEVLG
O. cuniculus       48 AWISETLCIGCGICIKKCPFGALSIVNLPSNLEKETTHRYCANAFKLHRLPIPRPGEVLG
D. rerio           48 VWISESLCIGCGICIKKCPFGALSIVNLPSNLEKETTHRYCANSFKLHRLPIPRPGEVLG
consensus          61 ....* .*.*****..***..*. *.***  *.....***  *.*.*..**..... ..*

S. solfataricus   104 VLGKNGVGKTTVLKILAGEIIPNFGDPNSKVGKDEVLKRFRGKEIYNYFKELYSNELKIV
P. furiosus       104 IVGPNGTGKSTAVKILAGQLIPNLCG--DNDSWDGVIRAFRGNELQNYFEKLKNGEIRPV
C. elegans        119 LVGTNGIGKSTALKILAGKQKPNLGNFQKEQEWTTIINHFRGSELQNYFTRILEDTLKCV
S. cerevisiae     108 LVGTNGIGKSTALKILAGKQKPNLGRFDDPPEWQEIIKYFRGSELQNYFTKMLEDDIKAI
O. sativa         108 LVGTNGIGKSTALKVLAGKLKPNLGRFKNPPDWQEILTYFRGSELQNYFTRILEDNLKAI
S. lycopersicum   108 LVGTNGIGKSTALKVLAGKLKPNLGRFTNPPDWQEILTHFRGSELQNYFTRILEDNLKAI
A. thaliana       108 LVGTNGIGKSTALKILAGKLKPNLGRFTSPPDWQEILTHFRGSELQNYFTRILEDNLKAI
C. hirsuta        108 LVGTNGIGKSTALKVLAGKLKPNLGRFTNPPDWQEILTHFRGSELQNYFTRILEDNLKAI
D. melanogaster   116 LVGQNGIGKSTALKILAGKQKPNLGKYANPPDWTEILSYFRGSELQNYFTKILEDNLKAL
H. sapiens        108 LVGTNGIGKSTALKILAGKQKPNLGKYDDPPDWQEILTYFRGSELQNYFTKILEDDLKAI
O. cuniculus      108 LVGTNGIGKSTALKILAGKQKPNLGKYDDPPDWQEILTYFRGSELQNYFTKILEDDLKAI
D. rerio          108 LVGTNGIGKSTALKILAGKQKPNLGRFDAPPDWQEILAYFRGSELQNYFTKILEDDLKAI
consensus         121 ..*.**.**.*..*.***. .**....  ........  ***.*..***...... ....

S. solfataricus   164 HKIQYVEYASKFLKGTVNEI---LTKIDER--GKKDEVKELLNMTNLWNKDANILSGGGL
P. furiosus       162 VKPQYVDLIPKAVKGKVIEL---LKKADET--GKLEEVVKALELENVLDREIQHLSGGEL
C. elegans        179 IKPQYVDQIPRAAKGTVEKN---LTRKHDN--DNLNSVIDQMELRGLLDREIDQLSGGEL
S. cerevisiae     168 IKPQYVDNIPRAIKGPVQKVGELLKLRMEKSPEDVKRYIKILQLENVLKRDIEKLSGGEL
O. sativa         168 IKPQYVDHIPKAVQGNVGQV---LDQKDER--GVKAELCVDLELNQVIDRNVGDLSGGEL
S. lycopersicum   168 IKPQYVDHIPKAVQGNVGQV---LDQKDER--DVKEKLCVDLELNQVLDRNVGDLSGGEL
A. thaliana       168 IKPQYVDHIPRAVKGNVGEV---LDQKDER--DKKAELCADLELNQVIDRDVENLSGGEL
C. hirsuta        168 IKPQYVDHIPKAVRGNVGEV---LDQKDER--NKKAELCADLELNQVIDRDVENLSGGEL
D. melanogaster   176 VKPQYVDQIPKAVRGAVGDL---LDKKDER--ELQTKICEMLDLSHIRDREIAQLSGGEL
H. sapiens        168 IKPQYVDQIPKAAKGTVGSI---LDRKDET--KTQAIVCQQLDLTHLKERNVEDLSGGEL
O. cuniculus      168 IKPQYVDQIPKAAKGTVGSI---LDRKDET--KTQAIVCQQLDLTHLKERNVEDLSGGEL
D. rerio          168 VKPQYVDQIPKTVKGSVGAI---LSRKDDT--KTEELVCEQLDLLHLRDRNVEDLSGGEL
consensus         181 .*.***. ......* *. .   *  ...        ..  ...  . .. .  ****.*
```

```
S. solfataricus   219 QRLLVAASLLREADVYIFDEPSSYLDVRERMNMAKAIRELLK-NKYVIVVDHDLIVLDYL
P. furiosus       217 QRVAIAAALLRNATFYFFDEPSSYLDIRQRLNAARAIRRLSEEGKSVLVVEHDLAVLDYL
C. elegans        234 QRFAIAMCCVQKADVYMFDEPSSYLDVKQRLKAAAIIRERVSDTNYVVVVEHDLAVLDYL
S. cerevisiae     228 QRFAIGMSCVQEADVYMFDEPSSYLDVKQRLNAAQIIRSLLAPTKYVICVEHDLSVLDYL
O. sativa         223 QRFAIAVVAVQNAEIFMFDEPSSYLDVKQRLKAAQVVRSLIRPNSYVIVVEHDLSVLDYL
S. lycopersicum   223 QRFAIAVVAIQNAEIYMFDEPSSYLDVKQRLKAAQVVRSLIRPNSYVIVVEHDLSVLDYL
A. thaliana       223 QRFAIAVVAIQNAEIYMFDEPSSYLDVKQRLKAAQVVRSLIRPNSYVIVVEHDLSVLDYL
C. hirsuta        223 QRFAIAVVAIQNAEIYMFDEPSSYLDVKQRLKAAQVVRSLIRPNSYVIVVEHDLSVLDYL
D. melanogaster   231 QRFAIAMVCIQNADIFMFDEPSSYLDVKQRLNAALTIRSLLHPTKFIIVVEHDLSVLDYL
H. sapiens        223 QRFACAVVCIQKADIFMFDEPSSYLDVKQRLKAAITIRSLINPDRYIIVVEHDLSVLDYL
O. cuniculus      223 QRFACAVVCIQKADIFMFDEPSSYLDVKQRLKAAITIRSLINPDRYIIVVEHDLSVLDYL
D. rerio          223 QRFACAVVCIQRADIFMFDEPSSYLDVKQRLRAAITIRSLISPDRYIIVVEHDLSVLDYL
consensus         241 **...... .. *....*********...*...*  .*... .  ....*.***.*****

S. solfataricus   278 TDLIHIIYGESSVYGRVSKSYAARVGINNFLKGYLPAENMKIRPDEIKFMLKEVSD-LDL
P. furiosus       277 SDIIHVVYGEPGVYGIFSQPKGTRNGINEFLRGYLKDENVRFRPYEIKFTKTGER----V
C. elegans        294 SDFICCLYGVPGVYGVVTLPSGVREGINMFLEGFIRTENMRFRESKLSFKTSEQQ----E
S. cerevisiae     288 SDFVCIIYGVPSVYGVVTLPASVREGINIFLDGHIPAENLRFRTEALQFRIADATEDL-Q
O. sativa         283 SDFICCLYGKPGAYGVVTLPFSVREGINIFLAGFVPTENLRFRDESLTFKIAET-QESAE
S. lycopersicum   283 SDFICCLYGKPGAYGVVTLPFSVREGINIFLAGFVPTENLRFRDESLTFKVAETPQEAAE
A. thaliana       283 SDFICCLYGKPGAYGVVTLPFSVREGINIFLAGFVPTENLRFRDESLTFKVAETPQESAE
C. hirsuta        283 SDFICCLYGKPGAYGVVTLPFSVREGINIFLAGFVPTENLRFRDESLTFKVAETPQESAE
D. melanogaster   291 SDFICCLYGVPGCYGVVTMPFSVREGINIFLDGFVPTENMRFRTESLTFKVSESAT--EE
H. sapiens        283 SDFICCLYGVPSAYGVVTMPFSVREGINIFLDGYVPTENLRFRDASLVFKVAETAN--EE
O. cuniculus      283 SDFICCLYGVPSAYGVVTMPFSVREGINIFLDGYVPTENLRFRDASLVFKVAETAN--EE
D. rerio          283 SDFICCLYGVPSAYGVVTMPFSVREGINIFLDGYVPTENLRFRETSLVFKVAETAA--EE
consensus         301 .*.....** .  **........*.***.** *....**...*. .. *.....     .

S. solfataricus   337 SKDLKTKMKWTKIIKKLGDEQLVVDNGEAKEGEIIGILGPNGIGKTTFARILVGEITADE
P. furiosus       333 EIERETLVTYPRLVKDYGSFRLEVEPGEIKKGEVIGIVGPNGIGKTTFVKMLAGVEEPTE
C. elegans        350 DIKRTGNIKYPSMSKTLGNFHLDVEAGDFSDSEIIVMLGENGTGKTTMIKMMAGSLKPED
S. cerevisiae     347 NDSASRAFSYPSLKKTQGDFVLNVEEGEFSDSEILVMMGENGTGKTTLIKLLAGALKPDE
O. sativa         342 EIETYQRYKYPTMSKTQGNFKLTVVEGEFTDSQIVVMLGENGTGKTTFIRMLAGLLKPDT
S. lycopersicum   343 EIESYARYRYPTMTKTQGNFKLKVSEGEFTDSQIVVMLGENGTGKTTFIRMLAGLLKPDV
A. thaliana       343 EIQSYARYKYPTMTKTQGNFRLRVSEGEFTDSQIIVMLGENGTGKTTFIRMLAGLLKPDD
C. hirsuta        343 EVQSYARYKYPTMSKTQGNFRLRVSEGEFTDSQIIVMLGENGTGKTTFIRMLAGLLKPDE
D. melanogaster   349 EIKRMNHYVYPAMVKTLGKFELTVEKGHFSDSEILVLLGENGTGKTTFIRMLAGNLQPDG
H. sapiens        341 EVKKMCMYKYPGMKKKMGEFELAIVAGEFTDSEIMVMLGENGTGKTTFIRMLAGRLKPDE
O. cuniculus      341 EVKKMCMYKYPGMKKKMGEFELAIVAGEFTDSEIMVMLGENGTGKTTFIRMLAGRLKPDE
D. rerio          341 EVKKLCRYQYPNMKKSMGEFTLTITEGEFTDSEIMVMLGENGTGKTTFIRMLAGGLKPDG
consensus         361 ..      .... . *. * * *  .  *..........*.**.****......*  .....

S. solfataricus   397 ----GSVTPEKQILSYKPQRIFPNYDGTVQQYLENASKDALSTSSWFFEEVTKRLNLHRL
P. furiosus       393 ----GKIEW-DLTVAYKPQYIKADYEGTVYELLSKIDASKLN-SNFYKTELLKPLGIIDL
C. elegans        410 E---NTELP-HVSISYKPQKISPKSETTVRFMLHDKIQNMYE-HPQFKTDVMNPLMMEQL
S. cerevisiae     407 ----GQDIP-KLNVSMKPQKIAPKFPGTVRQLFFKKIRGQFL-NPQFQTDVVKPLRIDDI
O. sativa         402 VEGTEIEIP-EFNVSYKPQKISPKFQHSVRHLLHQKIRDSYT-HPQFVSDVMKPLQIEQL
S. lycopersicum   403 VEGSDIEMP-EFNVSYKPQKISPKFQSTVRHLLHQKIRDSYQ-HPQFCSDVMKPLQIEQL
A. thaliana       403 TEGPDREIP-EFNVSYKPQKISPKFQNSVRHLLHQKIRDSYM-HPQFMSDVMKPLQIEQL
C. hirsuta        403 TEGPDREIP-EFNVSYKPQKISPKFQNSVRHLLHQKIRDSYM-HPQFVSDVMKPLQIEQL
D. melanogaster   409 ----EVELP-MLNISYKPQKISPKFQNHVRHLLHDKIRDAYV-HPQFIADVMKPMKIEEI
H. sapiens        401 ----GGEVP-VLNVSYKPQKISPKSTGSVRQLLHEKIRDAYT-HPQFVTDVMKPLQIENI
O. cuniculus      401 ----GGEVP-VLNVSYKPQKISPKSTGSVRQLLHEKIRDAYT-HPQFVTDVMKPLQIENI
D. rerio          401 ----GGDVP-ILNVSYKPQKISPKFKGSVRALLHDKIRDAYT-HPQFVTDVMKPMQIESI
consensus         421        ...    ....***.*....  .*. ... .... .  .... ......... .
```

```
S. solfataricus  453 LESNVNDLSGGELQKLYIAATLAKEADLYVLDEPSSYLDVEERYIVAKAIKRVTRERKAV
P. furiosus      447 YDREVNELSGGELQRVAIAATLLRDADIYLLDEPSAYLDVEQRLAVSRAIRHLMEKNEKT
C. elegans       465 LDRNVKELSGGELQRVALALCLGKTASLYLIDEPSAYLDSEQRLHAAKVIKRFIMHAKKT
S. cerevisiae    461 IDQEVQHLSGGELQRVAIVLALGIPADIYLIDEPSAYLDSEQRIICSKVIRRFILHNKKT
O. sativa        460 MDQEVINLSGGELQRVAICLCLGKPADIYLIDEPSAYLDSEQRIVASKVIKRFILHAKKT
S. lycopersicum  461 MDQEVVNLSGGELQRVALALCLGKPADIYLIDEPSAYLDSEQRIVASKVIKRFILHAKKT
A. thaliana      461 MDQEVVNLSGGELQRVALTLCLGKPADIYLIDEPSAYLDSEQRIVASKVIKRFILHAKKT
C. hirsuta       461 MDQEVINLSGGELQRVALALCLGKPADIYLIDEPSAYLDSEQRIVASKVIKRFILHAKKT
D. melanogaster  463 MDQEVQNLSGGELQRVALVLCLGKPADVYLIDEPSAYLDSEQRLVAAKVIKRYILHAKKT
H. sapiens       455 IDQEVQTLSGGELQRVALALCLGKPADVYLIDEPSAYLDSEQRLMAARVVKRFILHAKKT
O. cuniculus     455 IDQEVQTLSGGELQRVALALCLGKPADVYLIDEPSAYLDSEQRLMAARVVKRFILHAKKT
D. rerio         455 IDQDVQNLSGGELQRVALALCLGKPADVYLIDEPSAYLDSEQRLMAARVIKRFILHAKKT
consensus        481 ....*  *******.......*...*..*..****.***.*.*... .............

S. solfataricus  513 TFIIDHDLSIHDYIADRIIVFKGEPEKAGLATSPVTLKTGMNEFLRELEVTFRRDAETGR
P. furiosus      507 ALVVEHDVLMIDYVSDRLMVFEGEPGKYGRALPPMGMREGMNRFLASIGITFRRDPDTGR
C. elegans       525 AFVVEHDFIMATYLADRVVVFEGQPSVKCTACKPQSLLEGMNRFLKMLDITFRRDQETYR
S. cerevisiae    521 AFIVEHDFIMATYLADKVIVFEGIPSKNAHARAPESLLTGCNRFLKNLNVTFRRDPNSFR
O. sativa        520 AFIVEHDFIMATYLADKVIVYEGRPSIDCTANAPQSLVSGMNKFLSHLDITFRRDPTNYR
S. lycopersicum  521 AFVVEHDFIMATYLADRVIVYEGTPSIDCVANAPQSLLTGMNLFLSHLNITFRRDPTNYR
A. thaliana      521 AFVVEHDFIMATYLADRVIVYEGQPSIDCTANCPQSLLSGMNLFLSHLNITFRRDPTNFR
C. hirsuta       521 AFVVEHDFIMATYLADRVIVYEGQPSIDCTANCPQSLLSGMNLFLSHLNITFRRDPTNFR
D. melanogaster  523 GFVVEHDFIMATYLADRVIVIEGQPSVKTTAFSPQSLLNGMNRFLELLGITFRRDPNNFR
H. sapiens       515 AFVVEHDFIMATYLADRVIVFDGVPSKNTVANSPQTLLAGMNKFLSQLEITFRRDPNNYR
O. cuniculus     515 AFVVEHDFIMATYLADRVIVFDGVPSKNTVANSPQTLLAGMNKFLSQLEITFRRDPNNYR
D. rerio         515 AFVVEHDFIMATYLADRVIVFDGIPSRTTNANAPQTLLAGMNKFLAQLEITFRRDPNNFR
consensus        541 .....**.....*..*...*..* *.    *  *.... *.*.**  . .*****. ..*

S. solfataricus  573 PRVNKIGSYLDRVQKERGDYYSMVLSTQ-
P. furiosus      567 PRANKEGSVKDREQKEKGEYYYIA-----
C. elegans       585 PRINKLDSVKDVDQKKSGQFFFLDDN---
S. cerevisiae    581 PRINKLDSQMDKEQKSSGNYFFLDNTGI-
O. sativa        580 PRINKLDSTKDREQKSAGSYYYLDD----
S. lycopersicum  581 PRINKLESTKDREQKSAGSYYYLDD----
A. thaliana      581 PRINKLESTKDREQKSAGSYYYLDD----
C. hirsuta       581 PRINKLESTKDREQKSAGSYYYLDD----
D. melanogaster  583 PRINKNNSVKDTEQKRSGQFFFLEDEACN
H. sapiens       575 PRINKLNSIKDVEQKKSGNYFFLDD----
O. cuniculus     575 PRINKLNSIKDVEQKKSGNYFFLDD----
D. rerio         575 PRINKLNSIKDVEQKKSGNYFFLDD----
consensus        601 **.**. * .* .**  * ......
```

**Fig. S7** Sequence conservation among ABCE orthologs. Multiple sequence alignment of full-length ABCE proteins from the archaea *Saccharolobus solfataricus* (Q980K5) and *Pyrococcus furiosus* (I6V0C7), and the eukaryotes *Caenorhabditis elegans* (Q9U2K8), *Saccharomyces cerevisiae* (Q03195), *Oryza sativa* (A0A0P0Y344), *Solanum lycopersicum* (A0A3Q7H7H5), *Arabidopsis thaliana* (At4g19210), *Cardamine hirsuta* (L7VNS9), *Drosophila melanogaster* (Q9VSS1), *Homo sapiens* (P61221), *Oryctolagus cuniculus* (G1SG72), and *Danio rerio* (Q6TNW3). Full-length protein sequences were obtained from UniProt (https://www.uniprot.org/), except that of *Arabidopsis thaliana*, which was obtained from The Arabidopsis Information Resource (TAIR; https://www.arabidopsis.org/). The alignment was obtained with Clustal Omega 1.2.4 with default settings, and shaded with BOXSHADE with output format RTF_new. Identical and similar residues across at least eight out of the twelve sequences are shaded in black and gray, respectively. Asterisks and dots indicate identical and similar residues, respectively. Numbers indicate residue positions. The conserved Pro138 residue, which is replaced by Ser in the *api7-1* mutant, is highlighted in red.

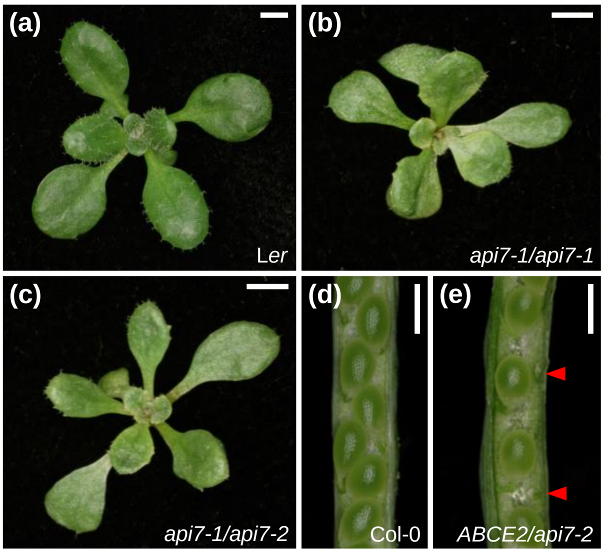


**Fig. S8** *api7-2* is a lethal allele of *ABCE2*. (a–c) Rosettes from (a) L*er*, (b) *api7-1/api7-1*, and (c) *api7-1/api7-2* plants. (d,e) Dissected immature siliques from (d) Col-0 and (e) *ABCE2/api7-2* plants. Red arrowheads indicate aborted seeds. Pictures were taken (a–c) 16 and (d,e) 57 das. Scale bars indicate (a–c) 2 mm, and (d,e) 500 µm.

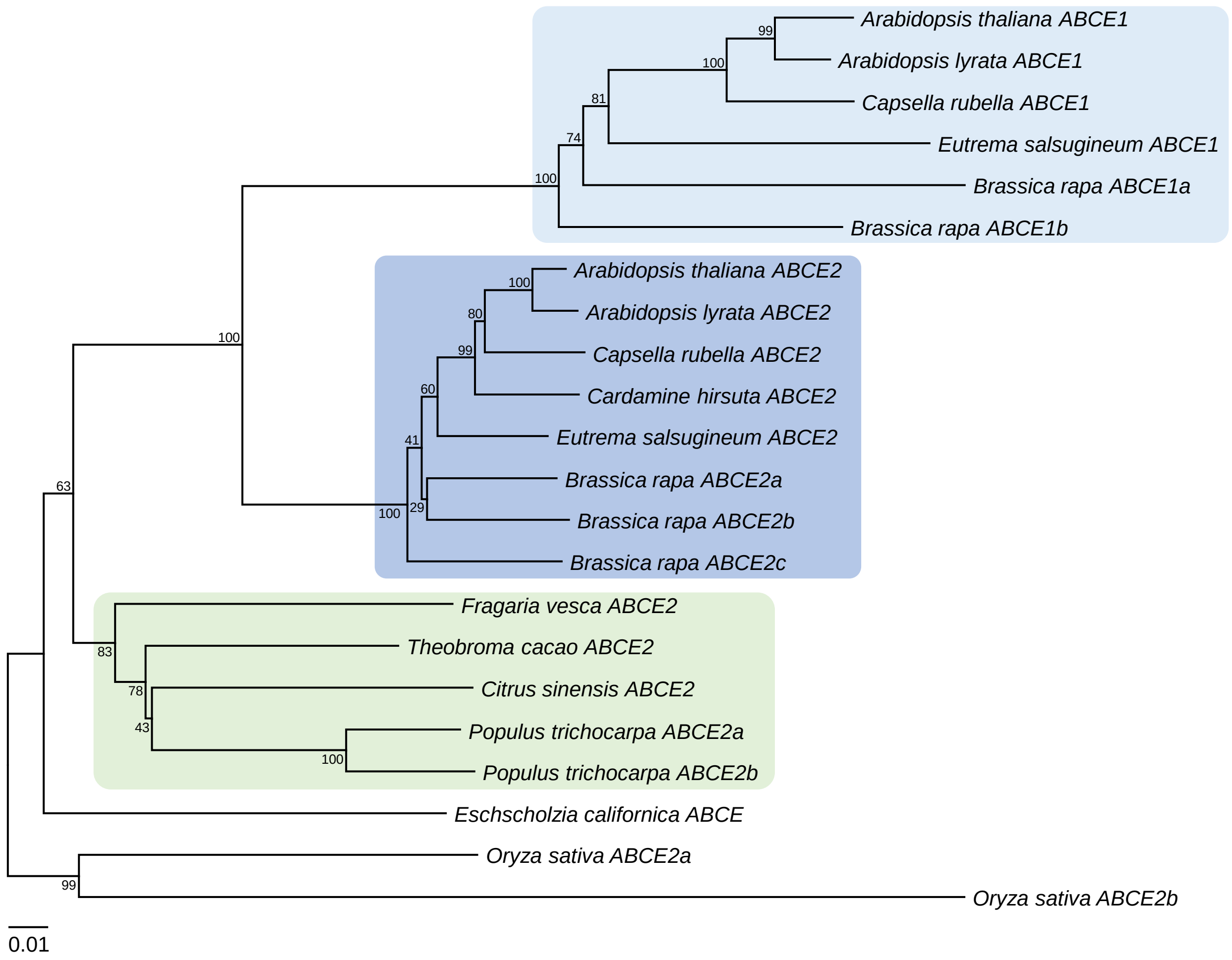


**Fig. S9** Phylogenetic analysis of some Rosidae *ABCE* genes. Rectangles indicate Brassicaceae *ABCE1* (clear blue), *ABCE2* (dark blue), and other rosid *ABCE2* (green) genes. *Eschscholzia californica* and *Oryza sativa ABCE* sequences were used as outgroups. Multiple *ABCE1* or *ABCE2* genes from *Brassica rapa*, *Populus trichocarpa*, and *Oryza sativa* are distinguished with arbitrarily given *a*, *b*, and *c* designations. Refer to Table S4 to see the NCBI Nucleotide codes of the sequences used during the multiple sequence alignment. The phylogenetic tree was obtained using the Neighbor-Joining method. All positions containing gaps and missing data were eliminated (complete deletion option). The percentage of replicate trees in which the associated taxa clustered together in the bootstrap test (1000 replicates) are shown next to the branches. The tree was rooted on the midpoint. The scalebar indicates the evolutionary distance as the number of base substitutions per site, and was computed using the Tamura 3-parameter method.

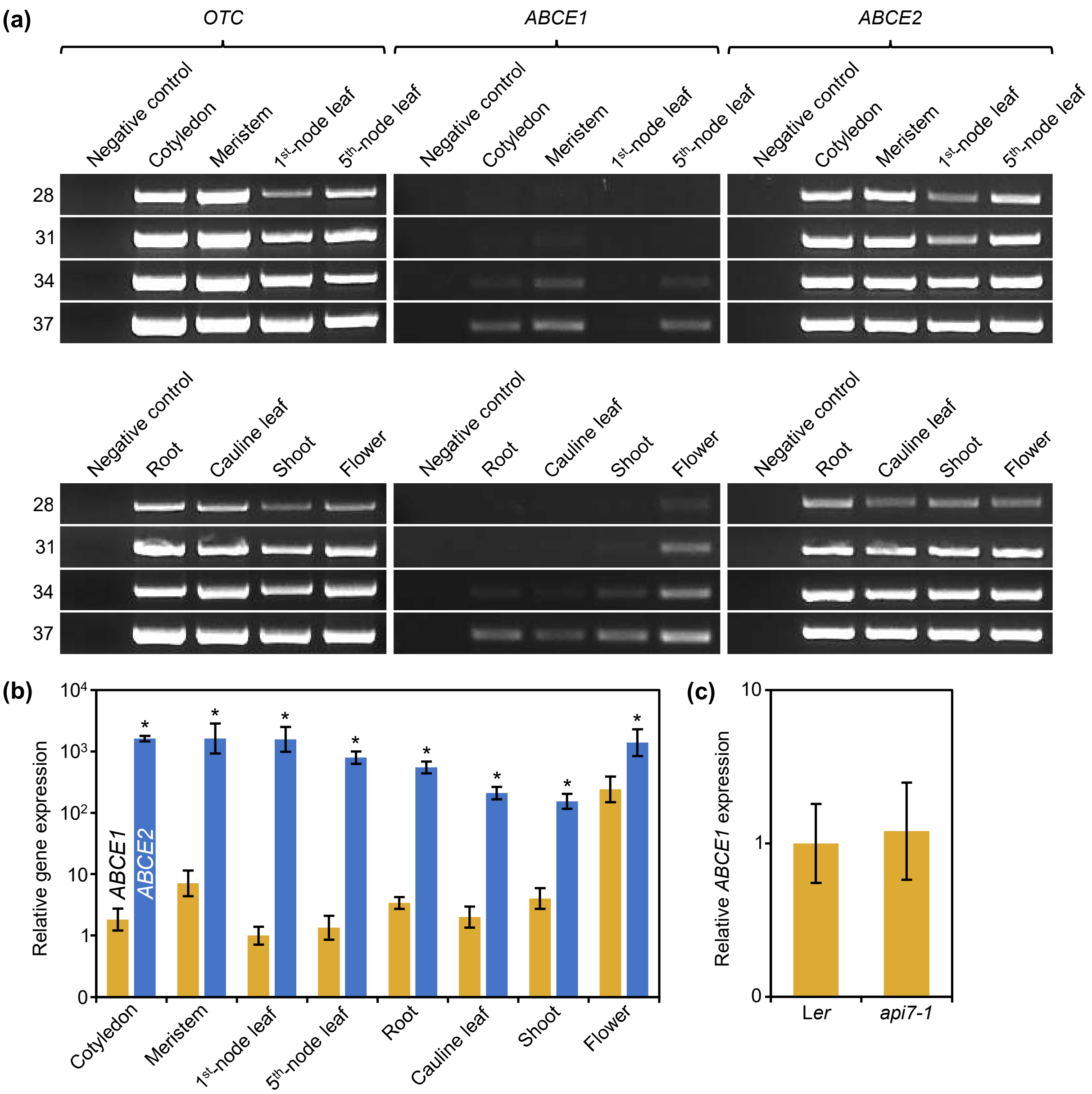


**Fig. S10** *ABCE1* and *ABCE2* expression analyses. (a) Semiquantitative and (b) quantitative PCR analyses of *ABCE1* and *ABCE2* expression in Col-0 plants. (a) The domestic *OTC* gene was used as a control. Negative control samples had no template. The PCR products were visualized after 28, 31, 34, and 37 amplification cycles. (a,b) Samples were collected 7 (cotyledons), 14 (meristems, first- and fifth-node leaves, and roots), or 28 (cauline leaves, shoots, and flowers) das. (c) *ABCE1* expression in wild-type L*er* and mutant *api7-1* first-node leaves. Samples were collected 14 das. (b,c) *ABCE1* expression levels in (b) first-node leaves and (c) L*er* were used as the reference value. Error bars indicate the interval delimited by $2^{-(\Delta\Delta C_T \pm SD)}$, where SD is the standard deviation of the $\Delta\Delta C_T$ values. Note that the relative gene expression levels are in a logarithmic scale. (b) Asterisks indicate values significantly different between *ABCE1* and *ABCE2* in a Mann-Whitney *U* test (**P* < 0.001). (b,c) Three different biological replicates were analyzed in triplicate.

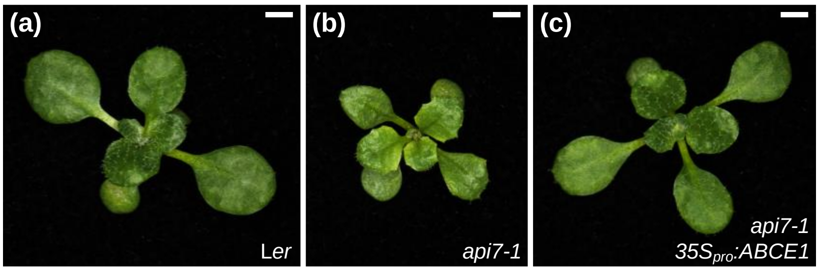


**Fig. S11** The *35S*$_{pro}$*:ABCE1* transgene restores the wild-type phenotype in *api7-1* plants. Rosettes from (a) L*er*, (b) *api7-1*, and (c) *35S*$_{pro}$*:ABCE1 api7-1* plants. Pictures were taken 14 das. Scale bars indicate 2 mm.

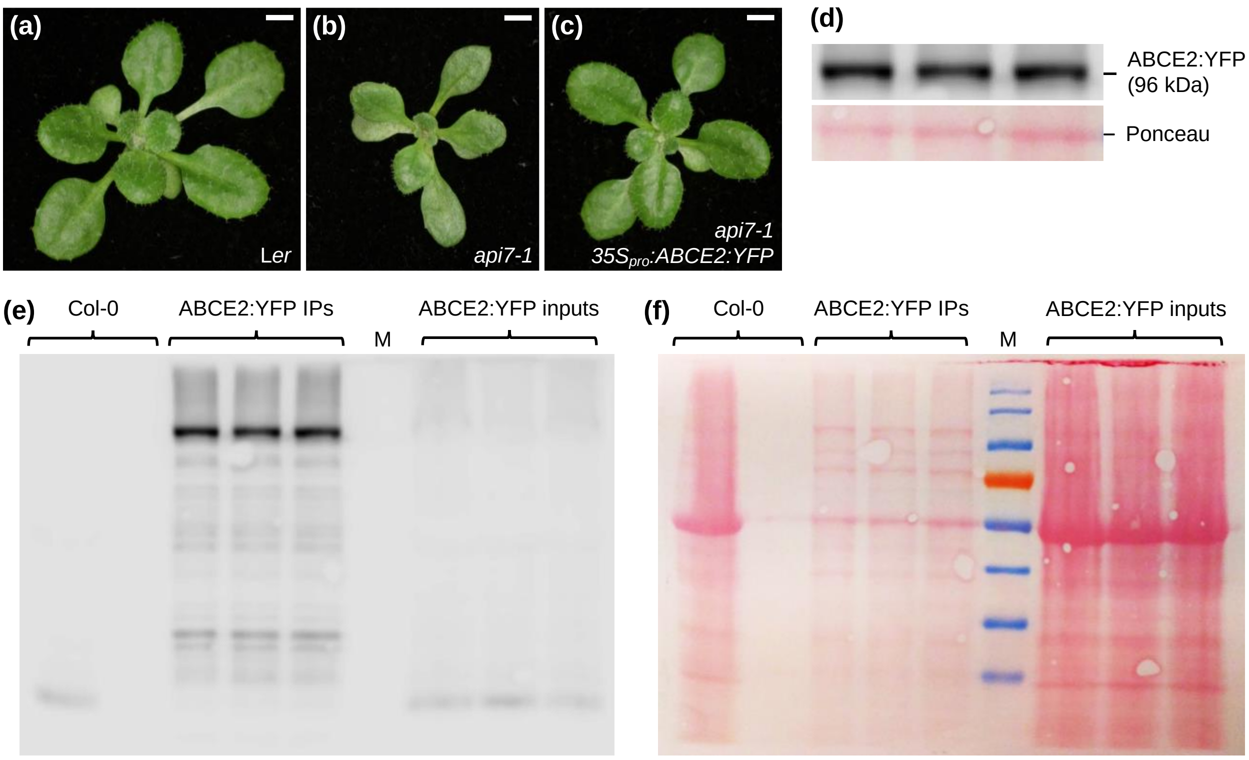


**Fig. S12** The *35S$_{pro}$:ABCE2:YFP* transgene fully restores the wild-type phenotype in *api7-1* plants. (a–c) Rosettes from (a) L*er*, (b) *api7-1*, and (c) *api7-1 35S$_{pro}$:ABCE2:YFP* plants. Pictures were taken 16 das. Scale bars indicate 2 mm. (c) The ABCE2:YFP fusion protein was detected from three independent immunoprecipitates in a western blot probed against GFP. Immunoprecipitates were obtained by immunoprecipitation with anti-GFP magnetic beads of whole-protein extracts from *api7-1 35S$_{pro}$:ABCE2:YFP* plants collected 10 das. A band from the Ponceau staining of the membrane is shown as a loading control. (e,f) Full pictures of (e) the detection of ABCE2:YFP and (f) the membrane stained with Ponceau, from the whole-protein extracts previous to immunoprecipitation (inputs) and the immunoprecipitated samples (IPs). A protein extract from wild-type Col-0 seedlings was used as a control: input and immunoprecipitation were loaded in the left and right lanes, respectively. M: EZ-Run Prestained *Rec* Protein Ladder (Thermo Fisher Scientific, Fisher BioReagents) molecular weight marker.

At4g19210 (ABCE2; 62%)

MADRLTRIAIVSSDRCKPKKCRQECKKSCPVVKTGKLCIEVTVGSKLAFISEELCIGCGICVKKCPFEAIQIINLPRDL
EKDTTHRYGANTFKLHRLPVPRPGQVLGLVGTNGIGKSTALKILAGKLKPNLGRFTSPPDWQEILTHFRGSELQNYFTR
ILEDNLKAIIKPQYVDHIPRAVKGNVGEVLDQKDERDKKAELCADLELNQVIDRDVENLSGGELQRFAIAVVAIQNAEI
YMFDEPSSYLDVKQRLKAAQVVRSLLRPNSYVIVVEHDLSVLDYLSDFICCLYGKPGAYGVVTLPFSVREGINIFLAGF
VPTENLRFRDESLTFKVAETPQESAEEIQSYARYKYPTMTKTQGNFRLRVSEGEFTDSQIIVMLGENGTGKTTFIRMLA
GLLKPDDTEGPDREIPEFNVSYKPQKISPKFQNSVRHLLHQKIRDSYMHPQFMSDVMKPLQIEQLMDQEVVNLSGGELQ
RVALTLCLGKPADIYLIDEPSAYLDSEQRIVASKVIKRFILHAKKTAFVVEHDFIMATYLADRVIVYEGQPSIDCTANC
PQSLLSGMNLFLSHLNITFRRDPTNFRPRINKLESTKDREQKSAGSYYYLDD

At3g13640 (ABCE1; 11%)

MSDRLTRIAIVSEDRCKPKKCRQECKKSCPVVKTGKLCIEVGSTSKSAFISEELCIGCGICVKKCPFEAIQIINLPKDL
AKDTTHRYGANGFKLHRLPIPRPGQVLGLVGTNGIGKSTALKILAGKLKPNLGRFNTPPDWEEILTHFRGSELQSYFIR
VVEENLKTAIKPQHVDYIKEVVRGNLGKMLEKLDERGLMEEICADMELNQVLEREARQVSGGELQRFAIAAVFVKKADI
YMFDEPSSYLDVRQRLKAAQVIRSLLRHDSYVIVVEHDLSVLDYLSDFVCCLYGKPGAYGVVTLPFSVREGINVFLAGF
IPTENLRFRDESLTFRVSETTQENDGEVKSYARYKYPNMTKQLGDFKLEVMEGEFTDSQIIVMLGENGTGKTTFIRMLA
GAFPREEGVQSEIPEFNVSYKPQGNDSKRECTVRQLLHDKIRDACAHPQFMSDVIRPLQIEQLMDQVVKTLSGGEKQRV
AITLCLGKPADIYLIDEPSAHLDSEQRITASKVIKRFILHAKKTAFIVEHDFIMATYLADRVIVYEGQPAVKCIAHSPQ
SLLSGMNHFLSHLNITFRRDPTNFRPRINKLESIKDKEQKTAGSYYYLDD

At4g11420 (eIF3a; 32%)

MANFAKPENALKRADELINVGQKQDALQALHDLITSKRYRAWQKPLEKIMFKYLDLCVDLKRGRFAKDGLIQYRIVCQQ
VNVSSLEEVIKHFLHLATDKAEQARSQADALEEALDVDDLEADRKPEDLQLSIVSGEKGKDRSDRELVTPWFKFLWETY
RTVLEILRNNSKLEALYAMTAHKAFQFCKQYKRTTEFRRLCEIIRNHLANLNKYRDQRDRPDLSAPESLQLYLDTRFDQ
LKVATELGLWQEAFRSVEDIYGLMCMVKKTPKSSLLMVYYSKLTEIFWISSSHLYHAYAWFKLFSLQKNFNKNLSQKDL
QLIASSVVLAALSIPPFDRAQSASHMELENEKERNLRMANLIGFNLEPKFEGKDMLSRSALLSELVSKGVLSCASQEVK
DLFHVLEHEFHPLDLGSKIQPLLEKISKSGGKLSSAPSLPEVQLSQYVPSLEKLATLRLLQQVSKIYQTIRIESLSQLV
PFFQFSEVEKISVDAVKNNFVAMKVDHMKGVVIFGNLGIESDGLRDHLAVFAESLSKVRAMLYPVPSKASKLAGVIPNL
ADTVEKEHKRLLARKSIIEKRKEDQERQQLEMEREEEQKRLKLQKLTEEAEQKRLAAELAERRKQRILREIEEKELEEA
QALLEETEKRMKKGKKKPLLDGEKVTKQSVKERALTEQLKERQEMEKKLQKLAKTMDYLERAKREEAAPLIEAAYQRRL
VEEREFYEREQQREVELSKERHESDLKEKNRLSRMLGNKEIFQAQVISRRQAEFDRIRTEREERISKIIREKKQERDIK
RKQIYYLKIEEERIRKLQEEEEARKQEEAERLKKVEAERKANLDKAFEKQRQREIELEEKSRREREELLRGTNAPPARL
AEPTVTPVGTTAPAAAAAAAGAPAAPYVPKWKRQTTEVSGPSAPTSSETDRRSNRGPPPGDDHWGSNRGAAQNTDRWTS
NRERSGPPAEGGDRWGSGPRGSDDRRSTFGSSRPRPTQR

At3g56150 (eIF3c; 14%)

MTSRFFTQVGSESEDESDYEVEVNEVQNDDVNNRYLQSGSEDDDDTDTKRVVKPAKDKRFEEMTYTVDQMKNAMKINDW
VSLQENFDKVNKQLEKVMRITEAVKPPTLYIKTLVMLEDFLNEALANKEAKKKMSTSNSKALNSMKQKLKKNNKLYEDD
INKYREAPEVEEEKQPEDDDDDDDDDDEVEDDDDSSIDGPTVDPGSDVDEPTDNLTWEKMLSKKDKLLEKLMNKDPKEI
TWDWVNKKFKEIVAARGKKGTARFELVDQLTHLTKIAKTPAQKLEILFSVISAQFDVNPGLSGHMPINVWKKCVLNMLT
ILDILVKYSNIVVDDTVEPDENETSKPTDYDGKIRVWGNLVAFLERVDTEFFKSLQCIDPHTREYVERLRDEPMFLALA
QNIQDYFERMGDFKAAAKVALRRVEAIYYKPQEVYDAMRKLAELVEEEEETEEAKEESGPPTSFIVVPEVVPRKPTFPE
SSRAMMDILVSLIYRNGDERTKARAMLCDINHHALMDNFVTARDLLLMSHLQDNIQHMDISTQILFNRTMAQLGLCAFR
AGMITESHSCLSELYSGQRVRELLAQGVSQSRYHEKTPEQERMERRRQMPYHMHLNLELLEAVHLICAMLLEVPNMAAN
SHDAKRRVISKNFRRLLEISERQAFTAPPENVRDHVMAATRALTKGDFQKAFEVLNSLEVWRLLKNRDSILDMVKDRIK
EEALRTYLFTYSSSYESLSLDQLAKMFDVSEPQVHSIVSKMMINEELHASWDQPTRCIVFHEVQHSRLQSLAFQLTEKL
SILAESNERAMESRTGGGGLDLSSRRRDNNQDYAGAASGGGGYWQDKANYGQGRQGNRSGYGGGRSSGQNGQWSGQNRG
GGYAGRVGSGNRGMQMDGSSRMVSLNRGVRT

At3g57290 (eIF3e; 40%)

MEESKQNYDLTPLIAPNLDRHLVFPIFEFLQERQLYPDEQILKSKIQLLNQTNMVDYAMDIHKSLYHTEDAPQEMVERR
TEVVARLKSLEEAAAPLVSFLLNPNAVQELRADKQYNLQMLKERYQIGPDQIEALYQYAKFQFECGNYSGAADYLYQYR
TLCSNLERSLSALWGKLASEILMQNWDIALEELNRLKEIIDSKSFSSPLNQVQNRIWLMHWGLYIFFNHDNGRTQIIDL
FNQDKYLNAIQTSAPHLLRYLATAFIVNKRRRPQLKEFIKVIQQEHYSYKDPIIEFLACVFVNYDFDGAQKKMKECEEV
IVNDPFLGKRVEDGNFSTVPLRDEFLENARLFVFETYCKIHQRIDMGVLAEKLNLNYEEAERWIVNLIRTSKLDAKIDS
ESGTVIMEPTQPNVHEQLINHTKGLSGRTYKLVNQLLEHTQAQATR

At1g64790 (ILA; 4%)

MSYSMVNASSAVSSPETAKNSDEPPPISSEAVNVLFPSVDPNSKLFRNSLNITISREAPPLTTSRIDFLSLFIFCKLTH
WLSLNPSSHRDEEEEEASPFYPFTIVLTYQPGPGQSPWKEMASPLESLLSISGSVSTSSTLIRLRIFRHDIPEILQNSD
MTSDIAPVIVDMIFQTLAIYDDRASRKAVDDLIVKGLGNVTFMKTFAAMLVQVMEKQLKFCFDTVCYRLLIWSCLLLEK
SQFATVSKNAFVRVASTQASLLRIIMESSFRMRRACKRFMFHLFSQSQAIYSLYMDEVKGSRIPYKDSPELLGLLLEFS
CSSPALFEQSKAIFVDIYVKDVLNSREKQKPNLSNCFKPLLQRLSHEEFQTVILPAAVKMLKRNPEIVLESVGFLLANV
NIDLSKYALELLPVILPQARHTDEDRRLGALSMVMCLSEKSSNPDTIEAMFASVKAIIGGSEGRLQSPHQRIGMLNAVQ
ELASAPEGKYIGSLSRTICSFLIACYKDEGNEDVKLSILSAVASWASRSSVAIQPNLVSFIAAGLKEKEALRRGHLRCV
RIICRNPDTISQISDLLSPLIQLVKTGFTKAVQRLDGIYALLIVSKIAACDIKAEDTMVKEKLWTLISQNEPSLVQITL
ASKLSSDDCVVCVDLLEVLLVEHSSRVLEAFSLKSLSQLLLFLLCHPSWNVRKTAYNSVTKIFLATSQLATTLLDEFSD
FLSITGDQIVSSRTSDADNPADHQAPFVPSVEVLVKALIVISSAAVAGPPSSWIVRAIFCSHHPSIVGTGKRDAVWKRL
QKCLKTCGFDVATFLSTNGESVCKSLLGPMGLTSAKTPEQQAAVYSLSTMMSLAPEDTFTVFKMHLQDLPDRLSHDMLS
ETDIKIFHTPEGMLLSEQGVYVAQTIGAKYTKQEPSSNHSLKKGLASRETANSGRRDTAKLTKKADKGKTAKEEARELM
LKEEASTRENVHRIQKSLSLVLHALGEMGLANPVFCHSQLPFLATFLDPLLRSPIVSAAAFENLVKLARCTVQPLCNWA
LEISTALRLIAIDEVDTSFDFRPSVDKAGKTYEGLFERIVNGLSISCKSGPLPVDTFTFIFPVLYHVLGVVPAYQASVG
PALNELCLGLQADDVANALYGVYSKDVHVRLACLNAVKCIPAVSKCSLPQNVKIATNIWIALHDPEKSVAESADDLWAR
YGHDLGTDYSGIFKALSHINLNVRLAAAEALADALHESPSSIQLSLSTLFSLYIRDATSGEDVFDAGWIGRQGIALALQ
SAADVLTTKDLPAVMTFLISRALADPNTDVRGKMINAGIMIIDKHGKENVSLLFPIFENYLNKEASDEEEYDLVREGVV
IFTGALAKHLARDDPKVHNVVEKLLEVLNTPSESVQRAVSTCLSPLVLSKQEEAPALFLRLLDKLMKSDKYGERRGAAF
GLAGVVMGFGISSLKKYGLIVTLQEALIDRNSAKRREGALLAFECLCEKLGKLFEPYVIKMLPLLLVSFSDQVGAVREA
AECAARAMMSQLSAYGVKLVLPSLLKGLEDKAWRTKQSSVQLLGAMAFCAPQQLSQCLPRVVPKLTEVFKTIQVLTDTH
PKVQSAGQLALQQVGSVIKNPEISSLVPTLLLALTDPNEYTRHALDTLLQTTFVNSVDAPSLALLVPIVHRGLRERSSE
TKKKASQIVGNMCSLVTEPKDMIPYIGLLLPEVKKVLVDPIPEVRSVAARAVGSLIRGMGEDNFPDLVPWLFETLKSDT
SNVERYGAAQGLSEVIAALGTDYFENILPDLIRHCSHQKASVRDGYLTLFKFLPRSLGAQFQKYLQLVLPAILDGLADE
NESVRDAALGAGHVLVEHHATTSLPLLLPAVEDGIFNDNWRIRQSSVELLGDLLFKVAGTSGKALLEGGSDDEGASTEA
QGRAIIDILGMDKRNEVLAALYMVRTDVSLSVRQAALHVWKTIVANTPKTLKEIMPILMSTLISSLASPSSERRQVAGR
SLGELVRKLGERVLPLIIPILSKGLKDPDVDKRQGVCIGLNEVMASAGRSQLLSFMDQLIPTIRTALCDSALEVRESAG
LAFSTLYKSAGLQAMDEIIPTLLEALEDDEMSTTALDGLKQIISVRTAAVLPHILPKLVHLPLSALNAHALGALAEVAG
AGFNTHLGTILPALLSAMGGENKEVQELAQEAAERVVLVIDEEGVETLLSELLKGVSDSQASIRRSSAYLIGYFFKSSK
LYLIDEAPNMISTLIVMLSDSDSTTVAVSWEALARVIGSVPKEVLPSYIKLVRDAVSTARDKERRKRKGGYVVIPGLCL
PKSLKPLLPVFLQGLISGSAELREQAAIGLGELIEVTSEQALKEFVIPITGPLIRIIGDRFPWQVKSAILATLIIILIQR
GGMALKPFLPQLQTTFVKCLQDSTRTIRSSAAVALGKLSALSTRIDPLVGDLMTSFQAADSGVREAILSAMRGVIKHAG
KSIGPAVRVRIFDLLKDLMHHEDDQVRISATSMLGVLSQYLEAAQLSVLLQEVNDLSASQNWGARHGSVLCISSLLKHN
PSTIMTSSLFSSMLNSLKSSLKDEKFPLRESSTKALGRLLLKQLATDPSNTKVVIDVLSSIVSALHDDSSEVRRRALSS
LKAFAKDNPSATMANISVIGPPLAECLKDGNTPVRLAAERCALHVFQLTKGAENVQAAQKYITGLDARRLSKFPEQSDD
SESDDDNVSG

At2g44060 (LEA26; 23%)

MSTSEDKPEIISRVVHQEGDVEIVDRSQKDKDEEKEEGKGGFLDKVKDFIHDIGEKLEGTIGFGKPTADVSAIHIPKIN
LERADIVVDVLVKNPNPVPIPLIDVNYLVESDGRKLVSGLIPDAGTLKAHGEETVKIPLTLIYDDIKSTYNDINPGMII
PYRIKVDLIVDVPVLGRLTLPLEKCGEIPIPKKPDVDIEKIKFQKFSLEETVAILHVRLQNMNDFDLGLNDLDCEVWLC
DVSIGKAEIADSIKLDKNGSGLINVPMTFRPKDFGSALWDMIRGKGTGYTIKGNIDVDTPFGAMKLPIIKEGGETRLKK
EDDDDDDEE

At4g20980 (eIF3d; 13%)

MVTEAFEFVAVPFNSDGWGPPDASDVSSSASPTSVAAANLLPNVPFASFSRSDKLGRVADWTRNLSNPSARPNTGSKSD
PSAVFDFSAFAIDEGFGLASSGGNPDEDAAFRLVDGKPPPRPKFGPKWRFNPHHNRNQLPQRRDEEVEAKKRDAEKERA
RRDRLYNNNRNNIHHQRREAAAFKSSVDIQPEWNMLEQIPFSTFSKLSYTVQEPEDLLLCGGLEYYNRLFDRITPKNER
RLERFKNRNFFKVTTSDDPVIRRLAKEDKATVFATDAILAALMCAPRSVYSWDIVIQRVGNKLFFDKRDGSQLDLLSVH
ETSQEPLPESKDDINSAHSLGVEAAYINQNFSQQVLVRDGKKETFDEANPFANEGEEIASVAYRYRRWKLDDNMHLVAR
CELQSVADLNNQRSFLTLNALNEFDPKYSGVDWRQKLETQRGAVLATELKNNGNKLAKWTAQALLANADMMKIGFVSRV
HPRDHFNHVILSVLGYKPKDFAGQINLNTSNMWGIVKSIVDLCMKLSEGKYVLVKDPSKPQVRIYEVPPDAFENDYVEE
PLPEDEQVQPTEENTEGAEASVAATKETEEKKADDAQA

At5g44320 (eIF3d; 9%)

MVFEAFEVGTVPFNSDGWGPPDASDTSSTSVAAANLLPNVPFASFSRSEKLGRVADWTRALSNPSARPHTGSKSDPSAI
FDFSAFAVDEGFGLTNSGGNADEDAAFRLVDGKPPPRPKFGPKWRFNQYHNRNQLPQRRDEEVEAKKREAEKDRARRDR
LYNNNRNNIHQQRREAAAFKSSVDIQPEWNMLEQIPFSTFSKLSFTVSEPEDLLLCGGLESYDRSFDRITPKADRRLER
FKNRSFKVTTSDDLVIRRLAKEDKATVFATDAILAALMCAPRSVYSWDLVIQRVGNKLFFDKRDGSPLDLLSVHETSQE
PLPEGKDDINSAHSLGLEAAYINQNFAQQVLVKNGKRETFDEPIPNVNEGEENASIAYRYRRWKLDDSMYLVARCELQS
TVDLNNQRSFLTLNALNEFDPKYSGVDWRQKLETQRGAVLANELKNNGNKLAKWTAQALLANADMMKIGFVSRVHPRDH
FNHVILSVLGYKPKDFAGQINLNTNNMWGIVKSIVDLCMKLSEGKYVLVKDPSKPQVRIYEVPADAFDNDYVEEPLPED
EQVQPPEENTDAGAETNGVSSTNVAVEDKKSEVEA

At5g17020 (XPO1A; 6%)

MAAEKLRDLSQPIDVGVLDATVAAFFVTGSKEERAAADQILRDLQANPDMWLQVVHILQNTNSLDTKFFALQVLEGVIK
YRWNALPVEQRDGMKNYISEVIVQLSSNEASFRSERLYVNKLNVILVQIVKHDWPAKWTSFIPDLVAAAKTSETICENC
MAILKLLSEEVFDFSRGEMTQQKIKELKQSLNSEFKLIHELCLYVLSASQRQDLIRATLSALHAYLSWIPLGYIFESTL
LETLLKFFPVPAYRNLTIQCLTEVAALNFGDFYNVQYVKMYTIFIGQLRIILPPSTKIPEAYSSGSGEEQAFIQNLALF
FTSFFKFHIRVLESTPEVVSLLLAGLEYLINISYVDDTEVFKVCLDYWNSLVLELFDAHHNSDNPAVSASLMGLQPFLP
GMVDGLGSQVMQRRQLYSHPMSKLRGLMINRMAKPEEVLIVEDENGNIVRETMKDNDVLVQYKIMRETLIYLSHLDHDD
TEKQMLRKLNKQLSGEEWAWNNLNTLCWAIGSISGSMAEDQENRFLVMVIRDLLNLCEITKGKDNKAVIASNIMYVVGQ
YPRFLRAHWKFLKTVVNKLFEFMHETHPGVQDMACDTFLKIVQKCKRKFVIVQVGENEPFVSELLTGLATTVQDLEPHQ
IHSFYESVGNMIQAESDPQKRDEYLQRLMALPNQKWAEIIGQARHSVEFLKDQVVIRTVLNILQTNTSAATSLGTYFLS
QISLIFLDMLNVYRMYSELVSTNITEGGPYASKTSFVKLLRSVKRETLKLIETFLDKAEDQPHIGKQFVPPMMESVLGD
YARNVPDARESEVLSLFATIINKYKATMLDDVPHIFEAVFQCTLEMITKNFEDYPEHRLKFFSLLRAIATFCFPALIKL
SSPQLKLVMDSIIWAFRHTERNIAETGLNLLLEMLKNFQQSEFCNQFYRSYFMQIEQEIFAVLTDTFHKPGFKLHVLVL
QQLFCLPESGALTEPLWDATTVPYPYPDNVAFVREYTIKLLSSSFPNMTAAEVTQFVNGLYESRNDPSGFKNNIRDFLV
QSKEFSAQDNKDLYAEEAAAQRERERQRMLSIPGLIAPNEIQDEMVDS

At3g03110 (XPO1B; 2%)

MAAEKLRDLSQPIDVVLLDATVEAFYSTGSKEERASADNILRDLKANPDTWLQVVHILQNTSSTHTKFFALQVLEGVIK
YRWNALPVEQRDGMKNYISDVIVQLSRDEASFRTERLYVNKLNIILVQIVKQEWPAKWKSFIPDLVIAAKTSETICENC
MAILKLLSEEVFDFSKGEMTQQKIKELKQSLNSEFQLIHELCLYVLSASQRQELIRATLSALHAYLSWIPLGYIFESPL
LEILLKFFPVPAYRNLTLQCLSEVASLNFGDFYDMQYVKMYSIFMNQLQAILPLNLNIPEAYSTGSSEEQAFIQNLALF
FTSFFKLHIKILESAPENISLLLAGLGYLISISYVDDTEVFKVCLDYWNSLVLELFGTRHHACHPALTPSLFGLQMAFL
PSTVDGVKSEVTERQKLYSDPMSKLRGLMISRTAKPEEVLIVEDENGNIVRETMKDNDVLVQYKIMRETLIYLSHLDHE
DTEKQMLSKLSKQLSGEEWAWNNLNTLCWAIGSISGSMVVEQENRFLVMVIRDLLSLCEVVKGKDNKAVIASNIMYVVG
QYSRFLRAHWKFLKTVVHKLFEFMHETHPGVQDMACDTFLKIVQKCKRKFVIVQVGESEPFVSELLSGLATIVGDLQPH
QIHTFYESVGSMIQAESDPQKRGEYLQRLMALPNQKWAEIIGQARQSADILKEPDVIRTVLNILQTNTRVATSLGTFFL
SQISLIFLDMLNVYRMYSELVSSSIANGGPYASRTSLVKLLRSVKREILKLIETFLDKAENQPHIGKQFVPPMMDQVLG
DYARNVPDARESEVLSLFATIINKYKVVMRDEVPLIFEAVFQCTLEMITKNFEDYPEHRLKFFSLLRAIATFCFRALIQ
LSSEQLKLVMDSVIWAFRHTERNIAETGLNLLLEMLKNFQKSDFCNKFYQTYFLQIEQEVFAVLTDTFHKPGFKLHVLV
LQHLFSLVESGSLAEPLWDAATVPHPYSNNVAFVLEYTTKLLSSSFPNMTTTEVTQFVNGLYESRNDVGRFKDNIRDFL
IQSKEFSAQDNKDLYAEEAAAQMERERQRMLSIPGLIAPSEIQDDMADS

At1g61580 (RPL3B; 12%)

MSHRKFEHPRHGSLGFLPRKRASRHRGKVKAFPKDDPTKPCRLTSFLGYKAGMTHIVRDVEKPGSKLHKKETCEAVTII
ETPPMVVVGVVGYVKTPRGLRSLCTVWAQHLSEELRRRFYKNWAKSKKKAFTRYSKKHETEEGKKDIQSQLEKMKKYCS
VIRVLAHTQIRKMKGLKQKKAHLNEIQINGGDIAKKVDYACSLFEKQVPVDAIFQKDEMIDIIGVTKGKGYEGVVTRWG
VTRLPRKTHRGLRKVACIGAWHPARVSYTVARAGQNGYHHRTEMNKKVYRVGKVGQETHSAMTEYDRTEKDITPMGGFP
HYGIVKEDYLMIKGCCVGPKKRVVTLRQTLLKQTSRLAMEEIKLKFIDAASNGGHGRFQTSQEKAKFYGRTIKA

At4g38740 (ROC1; 38%)

MAFPKVYFDMTIDGQPAGRIVMELYTDKTPRTAENFRALCTGEKGVGGTGKPLHFKGSKFHRVIPNFMCQGGDFTAGNG
TGGESIYGSKFEDENFERKHTGPGILSMANAGANTNGSQFFICTVKTDWLDGKHVVFGQVVEGLDVVKAIEKVGSSSGK
PTKPVVVADCGQLS

At3g13460 (ECT2; 11%)

MATVAPPADQATDLLQKLSLDSPAKASEIPEPNKKTAVYQYGGVDVHGQVPSYDRSLTPMLPSDAADPSVCYVPNPYNP
YQYYNVYGSGQEWTDYPAYTNPEGVDMNSGIYGENGTVVYPQGYGYAAYPYSPATSPAPQLGGEGQLYGAQQYQYPNYF
PNSGPYASSVATPTQPDLSANKPAGVKTLPADSNNVASAAGITKGSNGSAPVKPTNQATLNTSSNLYGMGAPGGGLAAG
YQDPRYAYEGYYAPVPWHDGSKYSDVQRPVSGSGVASSYSKSSTVPSSRNQNYRSNSHYTSVHQPSSVTGYGTAQGYYN
RMYQNKLYGQYGSTGRSALGYGSSGYDSRTNGRGWAATDNKYRSWGRGNSYYYGNENNVDGLNELNRGPRAKGTKNQKG
NLDDSLEVKEQTGESNVTEVGEADNTCVVPDREQYNKEDFPVDYANAMFFIIKSYSEDDVHKSIKYNVWASTPNGNKKL
AAAYQEAQQKAGGCPIFLFFSVNASGQFVGLAEMTGPVDFNTNVEYWQQDKWTGSFPLKWHIVKDVPNSLLKHITLENN
ENKPVTNSRDTQEVKLEQGLKIVKIFKEHSSKTCILDDFSFYEVRQKTILEKKAKQTQKQVSEEKVTDEKKESATAESA
SKESPAAVQTSSDVKVAENGSVAKPVTGDVVANGC

At4g33250 (eIF3k; 23%)

MGVEIQSPQEQSSYTVEQLVALNPFNPEILPDLENYVNVTSQTYSLEVNLCLLRLYQFEPERMNTHIVARILVKALMAM
PTPDFSLCLFLIPERVQMEEQFKSLIVLSHYLETGRFQQFWDEAAKNRHILEAVPGFEQAIQAYASHLLSLSYQKVPRS
VLAEAVNMDGASLDKFIEQQVTNSGWIVEKEGGSIVLPQNEFNHPELKKNTGENVPLEHIARIFPILG

At1g76810 (eIF5B; 4%)

MGRKKPSARGGDAEQQPPASSLVGATKSKKKGAQIDDDEYSIGTELSEESKVEEEKVVVITGKKKGKKGNKKGTQQDDD
DDFSDKVSAAGVKDDVPEIAFVGKKKSKGKKGGGSVSFALLDDEDEKEDNESDGDKDDEPVISFTGKKHASKKGKKGGN
SFAASAFDALGSDDDDTEEVHEDEEEESPITFSGKKKKSSKSSKKNTNSFTADLLDEEEGTDASNSRDDENTIEDEESP
EVTFSGKKKSSKKKGGSVLASVGDDSVADETKTSDTKNVEVVETGKSKKKKKNNKSGRTVQEEEDLDKLLAALGETPAA
ERPASSTPVEEKAAQPEPVAPVENAGEKEGEEETAAAKKKKKKKEKEKEKKAAAAAAATSSVEVKEEKQEESVTEPLQP
KKKDAKGKAAEKKIPKHVREMQEALARRQEAEERKKKEEEEKLRKEEEERRRQEELEAQAEEAKRKRKEKEKEKLLRKK
LEGKLLTAKQKTEAQKREAFKNQLLAAGGGLPVADNDGDATSSKRPIYANKKKSSRQKGIDTSVQGEDEVEPKENQADE
QDTLGEVGLTDTGKVDLIELVNTDENSGPADVAQENGVEEDDEEDEWDAKSWGTVDLNLKGDFDDEEEEAQPVVKKELK
DAISKAHDSEPEAEKPTAKPAGTGKPLIAAVKATPEVEDATRTKRATRAKDASKKGKGLAPSESIEGEENLRSPICCIM
GHVDTGKTKLLDCIRGTNVQEGEAGGITQQIGATYFPAENIRERTKELKADAKLKVPGLLVIDTPGHESFTNLRSRGSS
LCDLAILVVDIMHGLEPQTIESLNLLRMRNTEFIVALNKVDRLYGWKTCKNAPIVKAMKQQNKDVINEFNLRLKNIINE
FQEQGLNTELYYKNKDMGDTFSIVPTSAISGEGVPDLLLWLVQWAQKTMVEKLTYVDEVQCTVLEVKVIEGHGTTIDVV
LVNGELHEGDQIVVCGLQGPIVTTIRALLTPHPMKELRVKGTYLHYKEIKAAQGIKITAQGLEHAIAGTALHVVGPDDD
IEAIKESAMEDMESVLSRIDKSGEGVYVQASTLGSLEALLEYLKSPAVKIPVSGIGIGPVHKKDVMKAGVMLERKKEYA
TILAFDVKVTTEARELADEMGVKIFCADIIYHLFDLFKAYIENIKEEKKKESADEAVFPCVLQILPNCVFNKKDPIVLG
VDVIEGILKIGTPICVPGREFIDIGRIASIENNHKPVDYAKKGNKVAIKIVGSNAEEQKMFGRHFDMEDELVSHISRRS
IDILKSNYRDELSLEEWKLVVKLKNIFKIQ

At3g53610 (RAB8; 18%)

MAAPPARARADYDYLIKLLLIGDSGVGKSCLLLRFSDGSFTTSFITTIGIDFKIRTIELDGKRIKLQIWDTAGQERFRT
ITTAYYRGAMGILLVYDVTDESSFNNIRNWIRNIEQHASDSVNKILVGNKADMDESKRAVPKSKGQALADEYGMKFFET
SAKTNLNVEEVFFSIAKDIKQRLADTDARAEPQTIKINQSDQGAGTSQATQKSACCGT

At5g37475 (eIF3j; 16%)

**MDDWEAEDFQPLPSKVELKSNWDDEDVDENDIKDSWEEEDVSAPPPIVKPASEKAPKKPAVKAVEKKVKTVEAPKGTSR**
**EEPLDPIAEKLRMQRLVEEADYQSTAELFGVKTEEKSVDMLIPKSESDFLDYAELISQRLVPFEKSFHYIGLLKAVMRL**
**SVANMKAADVKDVASSITAIANEKLKAEKEAAAGKKKSGKKKQLHVDKPDDDLVSGPYDAMDDDDFM**

At3g43600 (AAO2; 3%)

**MSLVFAINGQRFELELSSVDPSTTLLEFLRYQTSFKSVKLSCGEGGCGACVVLLSKFDPVLQKVEDFTVSSCLTLLCSV**
**NHCNITTSEGLGNSRDGFHPIHKRLSGFHASQCGFCTPGMSVSLFSALLDADKSQYSDLTVVEAEKAVSGNLCRCTGYR**
**PIVDACKSFASDVDIEDLGLNSFCRKGDKDSSSLTRFDSEKRICTFPEFLKDEIKSVDSGMYRWCSPASVEELSSLLEA**
**CKANSNTVSMKLVAGNTSMGYYKDEREQNYDKYIDITRIPHLKEIRENQNGVEIGSVVTISKVIAALKEIRVSPGVEKI**
**FGKLATHMEMIAARFIRNFGSIGGNLVMAQRKQFPSDMATILLAAGAFVNIMSSSRGLEKLTLEEFLERSPLEAHDLVL**
**SIEIPFWHSETNSELFFETYRAAPRPHGSALAYLNAAFLAEVKDTMVVNCRLAFGAYGTKHAIRCKEIEEFLSGKVITD**
**KVLYEAITLLGNVVVPEDGTSNPAYRSSLAPGFLFKFLHTLMTHPTTDKPSNGYHLDPPKPLPMLSSSQNVPINNEYNP**
**VGQPVTKVGASLQASGEAVYVDDIPSPTNCLYGAFIYSKKPFARIKGIHFKDDLVPTGVVAVISRKDVPKGGKNIGMKI**
**GLGSDQLFAEDFTTSVGECIAFVVADTQRHADAAVNLAVVEYETEDLEPPILSVEDAVKKSSLFDIIPFLYPQQVGDTS**
**KGMAEADHQILSSEIRLGSQYVFYMETQTALAVGDEDNCIVVYSSTQTPQYVQSSVAACLGIPENNIRVITRRVGGGFG**
**GKSVKSMPVATACALAAKKLQRPVRTYVNRKTDMIMTGGRHPMKITYSVGFKSTGKITALELEILIDAGASYGFSMFIP**
**SNLIGSLKKYNWGALSFDIKLCKTNLLSRAIMRSPGDVQGTYIAEAIIENIASSLSLEVDTIRKINLHTHESLALFYKD**
**GAGEPHEYTLSSMWDKVGVSSKFEERVSVVREFNESNMWRKRGISRVPIIYEVLLFATPGRVSVLSDGTIVVEIGGIEL**
**GQGLWTKVKQMTSYALGMLQCDGTEELLEKIRVIQSDSLSMVQGNFTGGSTTSEGSCAAVRLCCETLVERLKPLMERSD**
**GPITWNELISQAYAQSVNLSASDLYTPKDTPMQYLNYGTAVSEVEVDLVTGQTTVLQTDILYDCGKSLNPAVDLGQIEG**
**SFVQGLGFFMLEEYIEDPEGLLLTDSTWTYKIPTVDTIPKQFNVEILNGGCHEKRVLSSKASGEPPLLLAASVHCATRQ**
**AVKEARKQLCMWKGENGSSGSAFQLPVPATMPVVKELCGLDIIESYLEWKLHDNSNL**

At1g65860 (FMO GS-OX1; 4%)

**MAPTQNTICSKHVAVIGAGAAGLVTARELRREGHTVVVFDREKQVGGLWNYSSKADSDPLSLDTTRTIVHTSIYESLRT**
**NLPRECMGFTDFPFVPRIHDISRDSRRYPSHREVLAYLQDFAREFKIEEMVRFETEVVCVEPVNGKWSVRSKNSVGFAA**
**HEIFDAVVVCSGHFTEPNVAHIPGIKSWPGKQIHSHNYRVPGPFNNEVVVVIGNYASGADISRDIAKVAKEVHIASRAS**
**ESDTYQKLPVPQNNLWVHSEIDFAHQDGSILFKNGKVVYADTIVHCTGYKYYFPFLETNGYININENRVEPLYKHVFLP**
**ALAPSLSFIGLPGMAIQFVMFEIQSKWVAAVLSGRVILPSQDKMMEDIIEWYATLDVLGIPKRHTHKLGKISCEYLNWI**
**AEECHCSPVENWRIQEVERGFQRMVSHPEIYRDEWDDDDLMEEAYKDFARKKLISSHPSYFLES**

At2g20830 (Folic acid binding / transferase; 8%)

**MSSGLNEDFLDCIVRLEETHVQQGFDEGYEEGLVSGREDARHLGLKLGFETGELIGFYRGCSALWNSALRIDPTRFSPQ**
**LHKHLNDFHVLLDKIPLLDPEDEAKDGIKDDLRVKFSIICASLGFSKKQFEWSEEMLREMLGCCKVYISEARNKTALEA**
**IERALKPFPPAAIVNKFEDAAYGRVGYTVVSSLANGSSSSLKNAVFAMVKTALDTINLELHCGSHPRLGVVDHICFHPL**
**SQTSIEQVSSVANSLAMDIGSILRVPTYLYGAAEKEQCTLDSIRRKLGYFKANREGHEWAGGFDLEMVPLKPDAGPQEV**
**SKAKGVVAVGACGWVSNYNVPVMSNDLKAVRRIARKTSERGGGLASVQTMALVHGEGVIEVACNLLNPSQVGGDEVQGL**
**IERLGREEGLLVGKGYYTDYTPDQIVERYMDLLNNS**

At5g58410 (HEAT/U-box domain-containing protein; 1%)

MGKPKGDAARSKARPSSSSLAASLLPSGSAAAVGFGGYVGSSRFQTSLSNEDSASFLDLDSEVAQHLQRLSRKDPTTKI
KALASLSELVKQKQGKELLPIIPQWTFEYKKLILDYSRDVRRATHDVMTNVVTGAGRDIAPHLKSIMGPWWFSQFDLAS
EVSQAAKSSFQVGSSFGNSVFLVEAAFPAQEKRLHALNLCSAEIFAYLEENLKLTPQNLSDKSLASDELEEMYQQMISS
SLVGLATLLDILLREPDNTGSANINSESKLASKARAVATSSAEKMFSSHKCFLNFLKSESPSIRSATYSLLSSFIKNVP
EVFGEGDVRSLAPALLGVFRENNPTCHSSMWEAVLLFSKKFPQSWVYLNVHKSVLNHLWQFLRNGCYGSPQVSYPALIL
FLEVMPAQSVESDKFFVNFFKNLLAGRSMCESSSTDQLSLLRATTECFLWGLRNASRYCDVPNSIHDLQVDLIDKVLVK
ILWADFTELSKGSIPPNQRKSAENLGMGNSVSYLQELGRCILEILSGINLLEQNLLSFFCKAVQESFLNMLQQGDLEIV
AGSMRKMIDFLLLLERYSVLEGESWPLHQFMGPLLSKAFPWIRSSELLDGVKLLSVSVSVFGPRKVVPVLIDDIETSTL
LSVEKEKNMSPEKLIKVFQEIFIPWCMDGYDSSTAARQDLLFSLLDDECFTQQWSDVISYVFNQQHQGFNNLAAMKMLL
EKARDEITKRSSGQELNQRIGSRPEHWHHTLIESTAISLVHSSSATTSAVQFLCSVLGGSTQDSSISFVSRSSLVLIY
RGILEKLLSFIKQSPLCSVNDTCSSLIVEAIAFDSSSSVDVIVVAKFAAEVIDGSFFSLKSLSQDATLLTTVLSSIFII
DLENRMTSLVDNTLSESKEKRKDRNFVCDYVHAVCSKMDNQFWKSINYDVRKSSASTLAQFLRSVVLLEDDLQPFELTL
LCASRMTEVLEYLSLDQSDEENICGLLLLESDAWPIWVSPSSSASIDTHGMPVQLCELRKSKSQRYVSFIDSLIMKLGI
HRFIVGHKDHGFASQAWLSVEILCTWEWPGGKVQTSFLPNLVSFCKDEPSSGGLLNSIFDILLNGALVHVKDEEEGLGN
MWVDFNNNIVDVVEPFLRALVSFLHILFKEDLWGEEEAMAAFKMITDKLFIGEETSKNCLRIIPYIMSIIISPLRTKVK
SGGSGKDTLLPLEVLLRNWLERSLSFPPLVLWQSGEDIQDWFQLVISCYPVSDKAEEAKELQRHLSTEERTLLLDLFRK
QKQDPGASTVVTQLPAVQILLARLIMIAVSYCGNDFNEDDWDFVFSNLKRLIQSAVVVMEETSENVNDFISGVSSMEKE
KENDTLEGLGHIVFISDPSINSAQNALSAFSLLNALVNHKSVEGEDNLKSLADETWDPVKDRILEGVLRLFFCTGLTEA
IAASYSPEAASIVASFRVDHLQFWELVAHLVVDSSPRARDRAVRAVEFWGLSRGSISSLYAIMFSSNPIPSLQLAAYTV
LSTEPISRLAIVADLNAPLNDESLNDQDSSNAGLPSEDKLLLRDEVSCMVEKLDHELLDTDLTAPERVQTFLAWSLLLS
NVNSLPSLTQGRERLVQYIEKTANPLILDSLFQHIPLELYMGQSLKKKDGDIPSELSVVASAATRAIITGSSLSTVESL
WPIETGKMASLAGAIYGLMLRVLPAYVREWFSEMRDRSASSLIEAFTRTWCSPSLIKNELSQIKKADFNDESFSVSISK
AANEVVATYTKDETGMDLVIRLPVSYPLKPVDVNCAKSIGISEAKQRKWLMSMQMFVRHQNGALAEAIRIWKRNSDKEF
EGVEDCPICYSVIHIGNHSLPRRACVTCKYKFHKACLDKWFYTSNKKLCPLCQSPC

At2g42910 (PRS4; 8%)

MSENAANNIMETKICTDAIVSELQKKKVHLFYCLECEELARNIAAESDHITLQSINWRSFADGFPNLFINNAHDIRGQH
VAFLASFSSPAVIFEQISVIYLLPRLFVASFTLVLPFFPTGSFERMEEEGDVATAFTMARIVSNIPISRGGPTSVVIYD
IHALQERFYFADQVLPLFETGIPLLTKRLQQLPETEKVIVAFPDDGAWKRFHKLLDHYPTVVCTKVREGDKRIVRLKEG
NPAGCHVVIVDDLVQSGGTLIECQKVLAAHGAVKVSAYVTHGVFPKSSWERFTHKKNGLEEAFAYFWITDSCPQTVKAI
GNKAPFEVLSLAGSIADALQI

At3g08850 (RAPTOR1; 1%)

MALGDLMVSRFSQSSVSLVSNHRYDEDCVSSHDDGDSRRKDSEAKSSSSYGNGTTEGAATATSMAYLPQTIVLCELRHD
ASEASAPLGTSEIVLVPKWRLKERMKTGCVALVLCLNITVDPPDVIKISPCARIEAWIDPFSMAPPKALETIGKNLSTQ
YERWQPRARYKVQLDPTVDEVRKLCLTCRKYAKTERVLFHYNGHGVPKPTANGEIWVFNKSYTQYIPLPISELDSWLKT
PSIYVFDCSAARMILNAFAELHDWGSSGSSGSSRDCILLAACDVHETLPQSVEFPADVFTSCLTTPIKMALKWFCRRSL
LKEIIDESLIDRIPGRQNDRKTLLGELNWIFTAVTDTIAWNVLPHELFQRLFRQDLLVASLFRNFLLAERIMRSANCNP
ISHPMLPPTHQHHMWDAWDMAAEICLSQLPQLVLDPSTEFQPSPFFTEQLTAFEVWLDHGSEHKKPPEQLPIVLQVLLS
QCHRFRALVLLGRFLDMGSWAVDLALSVGIFPYVLKLLQTTTNELRQILVFIWTKILALDKSCQIDLVKDGGHTYFIRF
LDSSGAFPEQRAMAAFVLAVIVDGHRRGQEACLEANLIGVCLGHLEASRPSDPQPEPLFLQWLCLCLGKLWEDFMEAQI
MGREANAFEKLAPLLSEPQPEVRAAAVFALGTLLDIGFDSNKSVVEDEFDDDEKIRAEDAIIKSLLDVVSDGSPLVRAE
VAVALARFAFGHKQHLKLAAASYWKPQSSSLLTSLPSIAKFHDPGSATIVSLHMSPLTRASTDSQPVARESRISSSPLG
SSGLMQGSPLSDDSSLHSDSGMMHDSVSNGAVHQPRLLDNAVYSQCVRAMFALAKDPSPRIASLGRRVLSIIGIEQVVA
KPSKPTGRPGEAATTSHTPLAGLARSSSWFDMHAGNLPLSFRTPPVSPPRTNYLSGLRRVCSLEFRPHLLGSPDSGLAD
PLLGASGSERSLLPLSTIYGWSCGHFSKPLLGGADASQEIAAKREEKEKFALEHIAKCQHSSISKLNNNPIANWDTRFE
TGTKTALLHPFSPIVVAADENERIRVWNYEEATLLNGFDNHDFPDKGISKLCLINELDDSLLLVASCDGSVRIWKNYAT
KGKQKLVTGFSSIQGHKPGARDLNAVVDWQQQSGYLYASGETSTVTLWDLEKEQLVRSVPSESECGVTALSASQVHGGQ
LAAGFADGSLRLYDVRSPEPLVCATRPHQKVERVVGLSFQPGLDPAKVVSASQAGDIQFLDLRTTRDTYLTIDAHRGSL
TALAVHRHAPIIASGSAKQLIKVFSLQGEQLGIIRYYPSFMAQKIGSVSCLTFHPYQVLLAAGAADSFVSIYTHDNSQA
R

At5g01770 (RAPTOR2; 1%)

MALGDLMVSRLSQSSVTVVTNHLYDDDDNCASSAHDDSRVSIIASPRVASSSYENLSAATSMAYLPQTLVLCDLRHDDA
SDIVQPPRWRLKERMKTGCVALVMCLHITVDPPDVIKISPCARLECWIDPFSMFPPRRALEAIGQNLSIQYERWLARAR
YKVELDPTKDDVRKLCLSCRKYAKTERVLFHYNGHGVPKPTPNGEIWVYNKNFTQYIPLPVSELDSWLKTPTIYVFDCS
AARVILNAFAEGESSGPPKDCILLAACDVHETLPQSVEFPADVFTSCLTTPINIALKWFCRRSLLKEFIDESLIDRIPG
RQNDRKTLLGELNWIFTAVTDTIAWNVLPRELFQRLFRQDLLVASLFRNFLLAERIMRSGNCTPISHPMLPPTHQHHMW
DAWDMAAEICLSQLPQFFLDPNTEFQPSSFFTEQLTAFEVWLDHGSEHKKPPEQLPIVLQVLLSQCHRYRALVLLGRFL
DMGPWAVDLALSVGIYPCVVKLLQTTTIELRQILVFIWTKILALDKSCQVDLVKDRGHIYFIRFLDSSDAFPEQRAMAA
FILAVIVDGYKRGQESCLEANLIAVCLGHLEATQLCDPPPEPLFLQWLCLCLGKLWEDYLEAQIMGREANASENLIAGH
TNLLQVRAAAVFALGTLLDVGFDSGKGVCDEEFDDDENIVEDIIIKSLLDVVSDGSPLVRTEVAVALARFAFGHKQHLK
SVADSYWKPQNSLRTSLPSMAKFHDSGTSIVASSDMGSLTRASPDSQPVAREGRISSSLQEPFSGLMQGSPLADDSSLH
SDVGIIHDGVSNGVVHQPRPLDNAIYSQSVLAMFTLAKDPSPRIASLGRRVLSVIGIEQIVAKPSKSNGRPGEAASASH
TPLAGLVRSSSWFDMHTGHLPLTFRTPPVSPPQTSYLTGLRRVCSLELRPHLLGSPDSGLADPILGVSGSERSLLPQST
IYNWSCGHFSKPLLGGADANEEIAAQREEKKKFSLEHIAKCQHSSISGLSNIPIANWDTKFETGTKTALLHPFSPIVVA
ADENERIRVWNYEEATLLNGFDNNDFPDKGISNLCLVNELDDSLLLVASCNVPTLSRASFAIRIWKDYATKGRQKLVTG
FSSIQGQKPGASGLNAVVDWQQQSGYLYVSGESLSIMVWDLDKEQLVKSMPFESGCSVTALSASQVHGSQLAAGFADGS
VRLYDVRTPDFLVCATRPHQRVEKVVGLSFQPGLDPAKIVSASQAGDIQFLDLRRPKETYLTIDAHRGSLTALGVHRHA
PIIASGSAKQLIKVFSLKGEQLGIIKYHTSFMGQQIGPVSCLAFHPYQMLLAAGAAGSFVSLYTHHNTQLPR

**Fig. S13** Amino acid sequences of proteins identified by LC-ESI-MS/MS in co-immunoprecipitated ABCE2:YFP protein. The Arabidopsis Genome Initiative (AGI) gene identifier (AtNgNNNNN) is shown, together with the TAIR10 annotation or protein name and peptide coverage (in percentage) for each protein. Full-length protein sequences were obtained from TAIR. The unique peptides identified by LC-ESI-MS/MS are shaded in black; some unique peptide sequences overlap. Peptide coverage was calculated by dividing the total number of residues of each protein by that of those covered by the peptides.

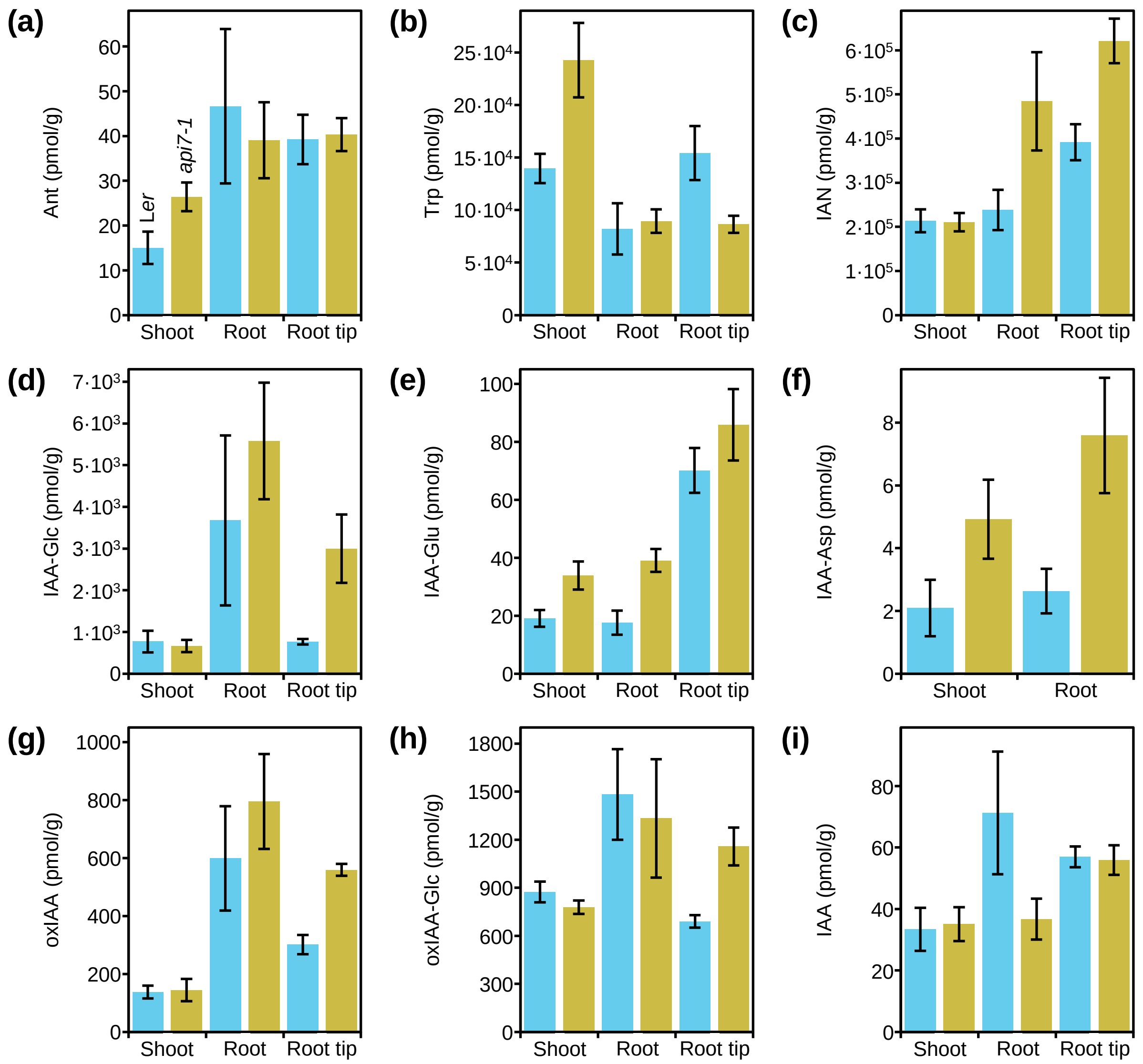


**Fig. S14** Tissue profiling of IAA metabolites in *api7-1* seedlings. The levels of (a–c) IAA precursors, (d) the IAA storage molecule IAA-Glc, (e–h) IAA catabolites, and (i) IAA were quantified in shoots, roots, and root tips from L*er* and *api7-1* seedlings 9 das. Concentrations are shown as the mean values from four biological replicates in pmol·g$^{-1}$ of fresh weight. Error bands represent the standard deviation.

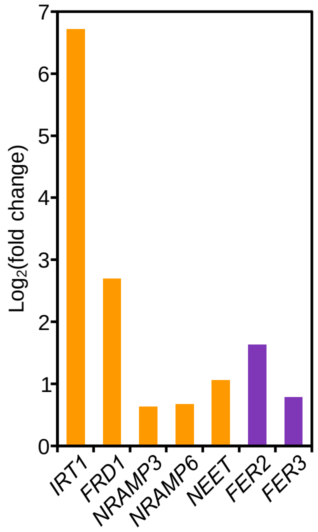


**Fig. S15** Expression levels of some genes deregulated in *api7-1* plants. Expression levels of genes related to iron homeostasis and FeS cluster biogenesis (orange), and response to oxidative stress (purple). Values are shown as the binary logarithm of the foldchange between *api7-1* and L*er* mean reads. Mean reads were calculated from three biological replicates.

**Table S1.** Primer sets used in this work

| Purpose | Name | Forward primer (F; 5′ → 3′) | Reverse primer (R; 5′ → 3′) |
|---|---|---|---|
| Linkage analysis | nga1111_F/R | GGGTTCGGTTACAATCGTGT | AGTTCCAGATTGAGCTTTGAGC |
| | AtF28J12.3_F/R | GCTCCGCCGTTGGATTCTG | GTTCGGGTTTAATTCTCGGGT |
| | AtM7J12.1_F/R | AGCAACTTGTGTTCTCATTT | TTATAGGGTACGACAACCAT |
| | nga1139_F/R | CTAGGCTCGGGTGAGTCAC | TTTTTCCTTGTGTTGCATTCC |
| | nga1107_F/R | GCGAAAAAACAAAAAAATCCA | CGACGAATCGACAGAATTAGG |
| | g3883_F/R | CATCCATCAAACAAACTCC | TGTTTCAGAGTAGCCAATTC |
| | T13K14_F/R | CTGAAACATATAAGAGAATCATCC | ACTCGTAGTTTGGTGTTGAGAC |
| | AtF16G20.1_F/R | TCAGTGTTACTATGTACCAAGTA | TAGGACGTAATATCCTTAGTTAC |
| | AG_F/R | CAACAGGTTTCTTCTTCTTCTC | CAAACACCATTTAATCTTGACA |
| | T18B16_F/R | TAACTTCTTGCAGCCTCTGAAG | TTCTATTGGGATGCTGCCCTC |
| Sequencing of *ABCE1* and *ABCE2* | ABCE1_F1/R1 | GGTTAGCTAGTCCCTTTCAAAG | GAAGTATGCTAATGTGGCCC |
| | ABCE1_F2/R2 | ACTACCTCTTGCGCAGACTC | GGCATCAGACTCACTTCATGA |
| | ABCE1_F3/R3 | GGAAGTGAAGTCCTATGCAAGA | CTTGAAATCTCCAAGTTGCTTAGT |
| | ABCE1_R4 | | CTGCCAAGATTTGGTTTGAG |
| | ABCE2_F1/R1 | TCGGTTCACCATTTTTATCTGAAG | CACCACAAGATGCTAACAATGAT |
| | ABCE2_F2/R2 | AGCCTGCGGATATATACCTGAT | GGTCGTATCTTTCTCCAAGTCT |
| | ABCE2_F3/R3 | GTTACACCGTATGGGCAAGAG | GGAGACTTACAGATAAGAAGAGA |
| | ABCE2_F4/R4 | CTTAGAACAATCGGCACACG | AGAGAAATCGAGATTAGTACCTGAG |
| | ABCE2_F5/R5 | GTTCTGATACCCTGTGCATG | ATCTCTTCAGCACTTTCTTGTG |
| Genotyping | api7-1_F/R | TGCCTCTAGAAATGGCACCT | GTATGGCAACAACAGCGATT |
| | GABI_509C06_LP/RP | TTCTTGGTCTGAAATTGGTGG | TGGCTGGATTTGTTCCTACAG |
| | o8409 (GABI-Kat lines)[1] | ATATTGACCATCATACTCATTGC | |
| | M13_F/R | TGTAAAACGACGGCCAGT | GGAAACAGCTATGACCATGATT |
| | GFP_R | | CACGTATCCCTCAGGCATGG |
| | YFP_R | | GACTTGAAGAAGTCGTGCTGC |

**Table S1 (continued).** Primer sets used in this work

| Purpose | Name | Forward primer (F; 5′ → 3′) | Reverse primer (R; 5′ → 3′) |
|---|---|---|---|
| qRT-PCR | qABCE1_F/R | CCTAAATCTTCGGAAAGTGAAC | GGCCATGAACCAACTTACGC |
| | qABCE2_F/R | GACAACTACCAAGAGAATATAGG | CAACTCAGGAAGTACAAAGCC |
| | qACTIN2_F/R[2] | GCACCCTGTTCTTCTTACCG | AACCCTCGTAGATTGGCACA |
| | OTC3D/OTCR[3] | TCCTTGCCAAATCATGGCCG | GCATGCATGCGATTCTCCGC |
| Gateway cloning | ABCE2pro:ABCE2_F/R | GGGGACAAGTTTGTACAAAAAAGCAGGCT | GGGGACCACTTTGTACAAGAAAGCTGGGT |
| | | TTTCTATCTTGTTATTCTTCGTTTTT | AGTTTTCCTATATCAGTGAGTTAG |
| | 35Spro:ABCE2:GFP-YFP_F/R | GGGGACAAGTTTGTACAAAAAAGCAGGCT | GGGGACCACTTTGTACAAGAAAGCTGGGT |
| | | GGATGGCAGATCGATTGACACGTA | CATCATCCAAGTAGTAGTATGAGC |
| | ABCE1pro_F/R | GGGGACAAGTTTGTACAAAAAAGCAGGCT | CAATACGTGTCAATCGATCTGCCATCTCTC |
| | | TACTTTTCTCTCGGCCGTACT | TTGAGAATATTACATACAAG |
| | ABCE2tu_F/R | CTTGTATGTAATATTCTCAAGAGAGATGGC | GGGGACCACTTTGTACAAGAAAGCTGGGT |
| | | AGATCGATTGACACGTATTG | CTAATCATCCAAGTAGTAGTA |
| | ABCE2pro_F/R | GGGGACAAGTTTGTACAAAAAAGCAGGCT | AATCCGCGTCAATCGATCTGACATCTCTCA |
| | | TTTCTATCTTGTTATTCTTCGTTTTT | ACAGACCTACAAAATACATG |
| | ABCE1tu_F/R | CATGTATTTTGTAGGTCTGTTGAGAGATGT | GGGGACCACTTTGTACAAGAAAGCTGGGT |
| | | CAGATCGATTGACGCGGATT | TCAATCGTCTAAGTAGTAGTA |

Sequences were taken from [1](S1), [2](S2), and [3](S3).

**Table S2.** Excitation and detection parameters of fluorophores

| Fluorophore | Laser type | Excitation (nm) | Detector type | Detection (nm) |
|---|---|---|---|---|
| GFP | Argon ion | 488 | Barrier filter | 515/30 |
| YFP | | | | |
| VENUS | | | | |
| DAPI | Diode | 408 | Barrier filter | 450/35 |
| Propidium iodide | Helium-neon | 543 | Barrier filter | 605/75 |

Nuclei or cell walls were stained by immersing complete seedlings in a 0.2 $\mu g \cdot ml^{-1}$ 4′,6-diamidino-2-phenylindole (DAPI) solution (Sony Biotechnology) for 12 min or a 10 $\mu g \cdot ml^{-1}$ propidium iodide solution (Sigma-Aldrich) for 8 min, respectively.

**Table S3.** Quality control summary of the RNA-seq assay

| Sample | Number of clean reads | Q30 quality score (%)* | Mapped reads (%) |
|---|---|---|---|
| L*er* replicate 1 | 31536086 | 95.11 | 95.40 |
| L*er* replicate 2 | 31336166 | 95.24 | 95.50 |
| L*er* replicate 3 | 30756110 | 94.99 | 95.43 |
| *api7-1* replicate 1 | 29977410 | 94.95 | 96.14 |
| *api7-1* replicate 2 | 27789921 | 95.39 | 96.35 |
| *api7-1* replicate 3 | 29383285 | 94.36 | 95.95 |

*Q30 quality score indicates the percentage of bases whose correct base recognition rates are greater than 99.9% in total bases.

**Table S4.** NCBI accession numbers of the sequences used for phylogenetic analysis

| Species | *ABCE1* gene | *ABCE2* gene |
|---|---|---|
| *Arabidopsis thaliana* | NM_112210.3 | ABCE2 NM_118041.5 |
| *Arabidopsis lyrata* subsp. *lyrata* | XM_021033623.1 | XM_021033596.1 |
| *Capsella rubella* | XM_006299733.2 | XM_006285967.2 |
| *Cardamine hirsuta* | - | JX097073.1 |
| *Eutrema salsugineum* | XM_006418662.2 | XM_006413929.2 |
| *Brassica rapa* (*a*) | XM_033286408.1 | XM_033276765.1 |
| *Brassica rapa* (*b*) | XM_009119375.3 | XM_009138752.3 |
| *Brassica rapa* (*c*) | - | XM_018655198.2 |
| *Fragaria vesca* subsp. *vesca* | - | XM_004291176.2 |
| *Theobroma cacao* | - | XM_018117748.1 |
| *Citrus sinensis* | - | XM_015531558.2 |
| *Populus trichocarpa* (*a*) | - | XM_024599960.1 |
| *Populus trichocarpa* (*b*) | - | XM_024597900.1 |
| *Eschscholzia californica* | - | Eca_sc194497.1_g0120.1* |
| *Oryza sativa* (*a*) | - | XM_015762126.2 |
| *Oryza sativa* (*b*) | - | XM_026023394.1 |

Multiple *ABCE1* or *ABCE2* genes from *Brassica rapa*, *Populus trichocarpa*, and *Oryza sativa* are distinguished with arbitrarily given *a*, *b*, and *c* designations. *Obtained from Eschscholzia Genome DataBase (http://eschscholzia.kazusa.or.jp/cgi-bin/list.cgi).

**Table S5.** Morphometry of the leaf venation pattern of the *api7-1* mutant

| Organ | Genotype | Area (mm$^2$) | Circularity | Vein density | Vein branching points | Free-ending veins |
|---|---|---|---|---|---|---|
| Cotyledons | L*er* | 2.9 ± 0.5 | 0.84 ± 0.03 | 2.7 ± 0.2 | 6.5 ± 0.5 | 2.1 ± 1.7 |
| | *api7-1* | ***1.9 ± 0.5*** | 0.85 ± 0.01 | ***3.1 ± 0.3*** | *8.2 ± 2.4* | **4.4 ± 2.2** |
| First-node leaves | L*er* | 33.9 ± 8.3 | 0.86 ± 0.02 | 3.1 ± 0.2 | 178.6 ± 35.6 | 77.8 ± 17.1 |
| | *api7-1* | ***11.2 ± 4.8*** | ***0.76 ± 0.05*** | 3.0 ± 0.3 | ***72.4 ± 24.6*** | ***35.9 ± 10.8*** |
| Third-node leaves | L*er* | 51.3 ± 11.2 | 0.85 ± 0.01 | 3.8 ± 0.3 | 361.7 ± 56.9 | 126.9 ± 24.4 |
| | *api7-1* | ***18.8 ± 4.0*** | *0.84 ± 0.02* | *3.5 ± 0.4* | ***145.2 ± 25.1*** | ***52.7 ± 10.6*** |
| Cauline leaves | L*er* | 200.6 ± 32.8 | 0.56 ± 0.23 | 3.9 ± 0.3 | 1307.6 ± 213.8 | 439.9 ± 66.1 |
| | *api7-1* | ***136.6 ± 50.5*** | *0.69 ± 0.04* | 4.2 ± 0.5 | **1023.0 ± 201.1** | *373.3 ± 73.4* |
| Sepals | L*er* | 1.4 ± 0.2 | 0.66 ± 0.05 | 6.6 ± 0.7 | 13.4 ± 4.2 | 9.5 ± 2.3 |
| | *api7-1* | *1.3 ± 0.1* | 0.65 ± 0.04 | 7.0 ± 1.0 | 14.7 ± 4.1 | 10.8 ± 4.8 |
| Petals | L*er* | 2.4 ± 0.4 | 0.70 ± 0.03 | 4.2 ± 0.6 | 6.4 ± 0.8 | 6.5 ± 1.8 |
| | *api7-1* | 2.2 ± 0.2 | 0.70 ± 0.03 | 4.0 ± 0.4 | 5.9 ± 1.1 | 7.7 ± 2.1 |

All values are means ± standard deviation from 12 measurements. Organs were collected 6 (cotyledons), 21 (first- and third-node leaves), and 35 (cauline leaves, petals, and sepals) das. Values in italics, bold, or bold and italics are significantly different from those of L*er* in a Student's *t* test, with $P < 0.05$, $P < 0.01$, or $P < 0.001$, respectively.

**Table S6.** Mutations identified in the *api7-1* candidate interval

| Mutation | Region affected | Predicted effect |
|---|---|---|
| G→A | At4g19185, 1st intron | - |
| G→A | At4g19185, 1st exon | Cys55→Cys (Synonymous) |
| C→T | At4g19210, 6th exon | Pro138→Ser |
| G→A | At4g19390, 2nd exon | Ile128→Ile (Synonymous) |

**Table S7.** ABCE2 interactors identified in a co-immunoprecipitation assay

| AGI gene code | Protein | | Peptides | | Peptide coverage |
|---|---|---|---|---|---|
| | Abbreviation | Full name | Total[1] | Unique[2] | (%) |
| At4g19210 | ABCE2 | ATP-BINDING CASSETTE E2 | 153 (20) | 26 (5) | 62 |
| At3g13640 | ABCE1 | ATP-BINDING CASSETTE E1 | | | |
| - | YFP | Yellow fluorescent protein | 59 | 10 | - |
| At4g11420 | eIF3a[3] | EUKARYOTIC TRANSLATION INITIATION FACTOR 3 SUBUNIT A | 48 | 24 | 32 |
| At3g56150 | eIF3c[3] | EUKARYOTIC TRANSLATION INITIATION FACTOR 3 SUBUNIT C | 29 | 11 | 14 |
| At3g57290 | eIF3e[3] | EUKARYOTIC TRANSLATION INITIATION FACTOR 3 SUBUNIT E | 25 | 15 | 40 |
| At1g64790 | ILA[3] | ILITYHIA | 17 | 11 | 4 |
| At2g44060 | LEA26[3] | LATE EMBRYOGENESIS ABUNDANT 26 | 15 | 6 | 23 |
| At4g20980 | eIF3d[3] | EUKARYOTIC TRANSLATION INITIATION FACTOR 3 SUBUNIT D | 13 (8) | 6 (4) | 13 |
| At5g44320 | eIF3d[3] | EUKARYOTIC TRANSLATION INITIATION FACTOR 3 SUBUNIT D | | | |
| At5g17020 | XPO1A[3] | EXPORTIN 1A | 12 (4) | 5 (2) | 6 |
| At3g03110 | XPO1B[3] | EXPORTIN 1B | | | |
| At1g61580 | RPL3B | RIBOSOMAL PROTEIN L3 B | 12 | 4 | 12 |
| At4g38740 | ROC1 | ROTAMASE CYP 1 | 11 | 5 | 38 |
| At3g13460 | ECT2[3] | EVOLUTIONARILY CONSERVED C-TERMINAL REGION 2 | 8 | 6 | 11 |
| At4g33250 | eIF3k[3] | EUKARYOTIC TRANSLATION INITIATION FACTOR 3 SUBUNIT K | 8 | 4 | 23 |
| At1g76810 | eIF5B[3] | EUKARYOTIC TRANSLATION INITIATION FACTOR 5B | 6 | 4 | 4 |
| At3g53610 | RAB8 | RAB GTPASE HOMOLOG 8 | 6 | 3 | 18 |
| At5g37475 | eIF3j | EUKARYOTIC TRANSLATION INITIATION FACTOR 3 SUBUNIT J | 5 | 3 | 16 |
| At3g43600 | AAO2 | ALDEHYDE OXIDASE 2 | 4 | 3 | 3 |
| At1g65860 | FMO GS-OX1 | FLAVIN-MONOOXYGENASE GLUCOSINOLATE S-OXYGENASE 1 | 4 | 3 | 4 |
| At2g20830 | -[4] | FOLIC ACID BINDING / TRANSFERASE | 4 | 2 | 8 |
| At5g58410 | LTN1 | E3 UBIQUITIN-PROTEIN LIGASE LISTERIN | 4 | 2 | 1 |

**Table S7 (continued).** ABCE2 interactors identified in a co-immunoprecipitation assay

| AGI gene code | Protein | | Peptides | | Peptide coverage |
|---|---|---|---|---|---|
| | Abbreviation | Full name | Total[1] | Unique[2] | (%) |
| At2g42910 | PRS4 | PHOSPHORIBOSYL DIPHOSPHATE SYNTHASE 4 | 4 | 2 | 8 |
| At3g08850 | RAPTOR1 | REGULATORY-ASSOCIATED PROTEIN OF TOR 1 | 4 (3) | 2 (2) | 1 |
| At5g01770 | RAPTOR2 | REGULATORY-ASSOCIATED PROTEIN OF TOR 2 | | | |

Three biological replicates were assayed and we identified 20 candidate interactors. Translation initiation factors were named according to (S4). [1]Sum of the number of peptides identified in the three biological replicates. [2]Number of associated peptides with significantly different sequences from each other. Values within parentheses refer to the second protein of the paralogous group, whose peptides were also associated with the first protein in all cases. [3]Enriched proteins (identified with at least twice the number of peptides associated with the same protein in the control co-immunoprecipitations). The rest of the proteins were unique to ABCE2:YFP samples. [4]This protein was included after being first discarded due to its predicted mitochondrial localization.

**Table S8.** Conservation level and described functions of putative ABCE2 interactors

| AGI gene code | Protein | Conservation between orthologs (%)[1] | | | Described molecular function(s) and available evidence of its interaction with ABCE proteins[2] |
|---|---|---|---|---|---|
| | | Arabidopsis and *S. cerevisiae* | Arabidopsis and *H. sapiens* | *S. cerevisiae* and *H. sapiens* | |
| At4g19210 | ABCE2 | 68.9 (82.6) | 75.4 (86.0) | 68.3 (82.9) | In Arabidopsis: suppression of RNA silencing (S5,S6).<br>In yeast (Rli1): ribosome dissociation (S7).<br>In humans (ABCE1): inhibition of RNAse L, suppression of RNA silencing, and ribosome dissociation (S8,S9,S10). |
| At4g11420 | eIF3a | 26.3 (43.4) | 25.7 (41.6) | 18.3 (32.0) | In Arabidopsis, yeast (Rpg1), and humans (EIF3A): translation initiation (S11). |
| At3g56150 | eIF3c | 24.3 (41.7) | 34.5 (49.9) | 23.4 (39.2) | In Arabidopsis, yeast (Nip1), and humans (EIF3C): translation initiation (S11). |
| At4g20980 | eIF3d | NC | 39.0 (54.4) | NC | In Arabidopsis and humans (EIF3D): translation initiation (S11). |
| At3g57290 | eIF3e | NC | 50.1 (69.0) | NC | In Arabidopsis and humans (EIF3E): translation initiation (S11). |
| At5g37475 | eIF3j | 22.5 (37.0) | 29.7 (45.1) | 26.2 (40.9) | In yeast (also named High-Copy suppressor of Rpg1 [Hcr1]): translation initiation and, as a non-stoichiometric subunit of the eIF3 complex, participates in pre-40S maturation, and as an accessory factor for Rli1-mediated ribosome dissociation (S12,S13).<br>In humans (EIF3J): start codon selection and eIF3 complex formation during translation initiation (S14,S15). It is also present in the 40S post-splitting complex with ABCE1 (S16). |
| At4g33250 | eIF3k | NC | 31.4 (50.7) | NC | In Arabidopsis and humans (EIF3K): translation initiation (S11) . |
| At1g76810 | eIF5B | 36.3 (50.1) | 39.7 (55.6) | 36.5 (54.1) | In yeast (Fun12): 40S and 60S joining during pre-40S maturation and translation initiation (S17,S18,S19,S20). |
| At1g61580 | RPL3B | 65.3 (79.6) | 64.9 (80.4) | 65.3 (80.0) | In Arabidopsis, yeast (Rpl3), and humans (RPL3): ribosomal protein. Translation (S21,S22). |

**Table S8 (continued).** Conservation level and described functions of putative ABCE2 interactors

| AGI code | Protein | Conservation between orthologs (%)[1] | | | Described molecular function(s) and available evidence of its interaction with ABCE proteins[2] |
|---|---|---|---|---|---|
| | | Arabidopsis and *S. cerevisiae* | Arabidopsis and *H. sapiens* | *S. cerevisiae* and *H. sapiens* | |
| At1g64790 | ILA | 27.6 (45.1) | 32.4 (50.2) | 27.7 (47.1) | In Arabidopsis: translation regulation through two pathways, one involving GCN2 and eIF2α, and the other involving GCN20 (S23,S24).<br>In yeast (Gcn1): translation downregulation by binding to translating ribosomes with Gcn2 and Gcn20 (S25,S26). |
| At3g13460 | ECT2 | 12.5 (19.6) | 25.5 (35.9) | 15.5 (24.6) | In Arabidopsis: m$^6$A reader that regulates 3′UTR processing in the nucleus and mRNA stability in the cytoplasm (S27,S28,S29).<br>In yeast (Pho92) and humans (YTHDF2): m$^6$A reader that decreases mRNA stability (S30,S31). |
| At5g58410 | LTN1 | 19.2 (36.1) | 22.5 (37.9) | 20.8 (36.8) | In yeast, *Drosophila melanogaster*, and humans: ubiquitination of nascent non-stop proteins for their degradation during ribosome quality control (S32,S33,S34). |
| At4g38740 | ROC1 | 64.0 (74.4) | 67.4 (80.2) | 64.2 (74.5) | In Arabidopsis, yeast (Cpr1), and humans (PPIA): it is a cyclophilin that belongs to the peptidyl-prolyl cis-trans isomerase family and participates in protein folding (S35,S36,S37). |
| At3g08850 | RAPTOR1 | 27.3 (42.1) | 40.4 (54.7) | 31.8 (45.3) | In Arabidopsis, yeast (Kog1), and humans (RAPTOR): it is part of the TORC1 complex, composed of TOR, RAPTOR, and LST8-1 proteins. It controls cellular growth in response to different signals through regulation of translation, as it promotes translation reinitiation and ribosome biogenesis (S38,S39).<br>In Arabidopsis: ABCE2 has been shown to interact with LST8-1 (S40). |
| At2g20830 | - | 26.7 (47.2) | 30.2 (43.4) | 27.8 (50.9) | In Arabidopsis: not studied. A BLASTp search suggested homology to *S. cerevisiae* and human Lto1.<br>In yeast and humans: FeS cluster assembly on Rli1 and ABCE1, respectively (S41,S42). |

**Table S8 (continued).** Conservation level and described functions of putative ABCE2 interactors

| AGI code | Protein | Conservation between orthologs (%)[1] | | | Described molecular function(s) and available evidence of its interaction with ABCE proteins[2] |
|---|---|---|---|---|---|
| | | Arabidopsis and *S. cerevisiae* | Arabidopsis and *H. sapiens* | *S. cerevisiae* and *H. sapiens* | |
| At5g17020 | XPO1A | 40.7 (61.7) | 48.5 (67.6) | 46.2 (65.8) | In Arabidopsis, yeast (Crm1), and humans (XPO1): nuclear export receptor (S43,S44).<br>In yeast, humans, and *Xenopus laevis*: ABCE might be an XPO1 cargo (S45).<br>In yeast: *xpo1-1* mutants accumulate ABCE1 in nucleus (S46,S47). |
| At2g42910 | PRS4 | 20.7 (38.2) | 19.2 (40.0) | 60.3 (76.6) | In yeast (Prs4) and humans (PRPS1): synthesis of phosphoribosylpyrophosphate (PRPP), which is required for nucleotide biosynthesis (S48). |
| At1g65860 | FMO GS-OX1 | NC | NC | NC | In Arabidopsis: synthesis of aliphatic glucosinolates (S49). |
| At3g53610 | RAB8 | 51.1 (70.0) | 58.7 (73.1) | 48.9 (67.0) | In Arabidopsis: it might be involved in post-Golgi transport to the plasma membrane (S50,S51).<br>In yeast (Sec4): involved in membrane trafficking during cytokinesis and autophagy (S52,S53). |
| At3g43600 | AAO2 | NC | 29.6 (48.1) | NC | In Arabidopsis: it might be involved in ABA biosynthesis (S54,S55).<br>In humans (AOX1): it is an oxidase with broad substrate specificity (S56). |
| At2g44060 | LEA26 | NC | NC | NC | In Arabidopsis: unknown. |

[1]Identity and similarity (between parentheses) percentages were obtained by global pairwise sequence alignments between pairs of protein sequences using the Needle EMBOSS tool. Protein sequences were obtained from TAIR for Arabidopsis, *Saccharomyces* Genome Database (SGD; https://www.yeastgenome.org/) for *S. cerevisiae*, and UniProt for *Homo sapiens* proteins. NC: not conserved. [2]The abbreviated names for *S. cerevisiae* and *Homo sapiens* orthologs are indicated in parentheses. The full names of Arabidopsis proteins are provided in Table S7.

**SUPPLEMENTARY REFERENCES**


S1. Kleinboelting, N., Huep, G., Kloetgen, A., Viehoever, P. and Weisshaar, B. (2012) GABI-Kat SimpleSearch: new features of the *Arabidopsis thaliana* T-DNA mutant database. *Nucleic Acids Res.*, **40**, D1211-1215.

S2. Moschopoulos, A., Derbyshire, P. and Byrne, M.E. (2012) The *Arabidopsis* organelle-localized glycyl-tRNA synthetase encoded by *EMBRYO DEFECTIVE DEVELOPMENT1* is required for organ patterning. *J. Exp. Bot.*, **63**, 5233-5243.

S3. Quesada, V., Ponce, M.R. and Micol, J.L. (1999) *OTC* and *AUL1*, two convergent and overlapping genes in the nuclear genome of *Arabidopsis thaliana*. *FEBS Lett.*, **461**, 101-106.

S4. Browning, K.S. and Bailey-Serres, J. (2015) Mechanism of cytoplasmic mRNA translation. *The Arabidopsis book*, **13**, e0176.

S5. Mõttus, J., Maiste, S., Eek, P., Truve, E. and Sarmiento, C. (2020) Mutational analysis of *Arabidopsis thaliana* ABCE2 identifies important motifs for its RNA silencing suppressor function. *Plant Biol.*, **23**, 21-31.

S6. Sarmiento, C., Nigul, L., Kazantseva, J., Buschmann, M. and Truve, E. (2006) AtRLI2 is an endogenous suppressor of RNA silencing. *Plant Mol. Biol.*, **61**, 153-163.

S7. Shoemaker, C.J. and Green, R. (2011) Kinetic analysis reveals the ordered coupling of translation termination and ribosome recycling in yeast. *Proc. Natl. Acad. Sci. USA*, **108**, E1392-E1398.

S8. Kärblane, K., Gerassimenko, J., Nigul, L., Piirsoo, A., Smialowska, A., Vinkel, K., Kylsten, P., Ekwall, K., Swoboda, P., Truve, E. *et al.* (2015) ABCE1 is a highly conserved RNA silencing suppressor. *PLOS ONE*, **10**, e0116702.

S9. Pisarev, A.V., Skabkin, M.A., Pisareva, V.P., Skabkina, O.V., Rakotondrafara, A.M., Hentze, M.W., Hellen, C.U. and Pestova, T.V. (2010) The role of ABCE1 in eukaryotic posttermination ribosomal recycling. *Mol. Cell*, **37**, 196-210.

S10. Bisbal, C., Martinand, C., Silhol, M., Lebleu, B. and Salehzada, T. (1995) Cloning and characterization of a RNase L inhibitor. A new component of the interferon-regulated 2-5A pathway. *J. Biol. Chem.*, **270**, 13308-13317.

S11. Burks, E.A., Bezerra, P.P., Le, H., Gallie, D.R. and Browning, K.S. (2001) Plant initiation factor 3 subunit composition resembles mammalian initiation factor 3 and has a novel subunit. *J. Biol. Chem.*, **276**, 2122-2131.

S12. Valášek, L., Hašek, J., Nielsen, K.H. and Hinnebusch, A.G. (2001) Dual function of eIF3j/Hcr1p in processing 20 S pre-rRNA and translation initiation. *J. Biol. Chem.*, **276**, 43351-43360.

S13. Young, D.J. and Guydosh, N.R. (2019) Hcr1/eIF3j is a 60S ribosomal subunit recycling accessory factor *in vivo*. *Cell Rep.*, **28**, 39-50.

S14. Borgo, C., Franchin, C., Salizzato, V., Cesaro, L., Arrigoni, G., Matricardi, L., Pinna, L.A. and Donella-Deana, A. (2015) Protein kinase CK2 potentiates translation efficiency by phosphorylating eIF3j at Ser127. *Biochim. Biophys. Acta*, **1853**, 1693-1701.

S15. ElAntak, L., Wagner, S., Herrmannová, A., Karásková, M., Rutkai, E., Lukavsky, P.J. and Valášek, L. (2010) The indispensable N-terminal half of eIF3j/HCR1 cooperates with its structurally conserved binding partner eIF3b/PRT1-RRM and with eIF1A in stringent AUG selection. *J. Mol. Biol.*, **396**, 1097-1116.

S16. Kratzat, H., Mackens-Kiani, T., Ameismeier, M., Potocnjak, M., Cheng, J., Dacheux, E., Namane, A., Berninghausen, O., Herzog, F., Fromont-Racine, M. *et al.* (2021) A structural inventory of native ribosomal ABCE1-43S pre-initiation complexes. *EMBO J.*, **40**, e105179.

S17. Fringer, J.M., Acker, M.G., Fekete, C.A., Lorsch, J.R. and Dever, T.E. (2007) Coupled release of eukaryotic translation initiation factors 5B and 1A from 80S ribosomes following subunit joining. *Mol. Cell. Biol.*, **27**, 2384-2397.

S18. Lebaron, S., Schneider, C., van Nues, R.W., Swiatkowska, A., Walsh, D., Böttcher, B., Granneman, S., Watkins, N.J. and Tollervey, D. (2012) Proofreading of pre-40S ribosome maturation by a translation initiation factor and 60S subunits. *Nat. Struct. Mol. Biol.*, **19**, 744-753.

S19. Strunk, B.S., Novak, M.N., Young, C.L. and Karbstein, K. (2012) A translation-like cycle is a quality control checkpoint for maturing 40S ribosome subunits. *Cell*, **150**, 111-121.

S20. Wang, J., Johnson, A.G., Lapointe, C.P., Choi, J., Prabhakar, A., Chen, D.H., Petrov, A.N. and Puglisi, J.D. (2019) eIF5B gates the transition from translation initiation to elongation. *Nature*, **573**, 605-608.

S21. García-Gómez, J.J., Fernández-Pevida, A., Lebaron, S., Rosado, I.V., Tollervey, D., Kressler, D. and de la Cruz, J. (2014) Final pre-40S maturation depends on the functional integrity of the 60S subunit ribosomal protein L3. *PLOS Genet.*, **10**, e1004205.

S22. Meskauskas, A. and Dinman, J.D. (2007) Ribosomal protein L3: gatekeeper to the A-site. *Mol. Cell*, **25**, 877-888.

S23. Faus, I., Niñoles, R., Kesari, V., Llabata, P., Tam, E., Nebauer, S.G., Santiago, J., Hauser, M.T. and Gadea, J. (2018) Arabidopsis ILITHYIA protein is necessary for proper chloroplast biogenesis and root development independent of eIF2$\alpha$ phosphorylation. *J. Plant Physiol.*, **224-225**, 173-182.

S24. Izquierdo, Y., Kulasekaran, S., Benito, P., López, B., Marcos, R., Cascón, T., Hamberg, M. and Castresana, C. (2018) Arabidopsis *nonresponding to oxylipins* locus *NOXY7* encodes a yeast GCN1 homolog that mediates noncanonical translation regulation and stress adaptation. *Plant Cell Environ.*, **41**, 1438-1452.

S25. Lee, S.J., Swanson, M.J. and Sattlegger, E. (2015) Gcn1 contacts the small ribosomal protein Rps10, which is required for full activation of the protein kinase Gcn2. *Biochem. J.*, **466**, 547-559.

S26. Sattlegger, E. and Hinnebusch, A.G. (2005) Polyribosome binding by GCN1 is required for full activation of eukaryotic translation initiation factor 2$\alpha$ kinase GCN2 during amino acid starvation. *J. Biol. Chem.*, **280**, 16514-16521.

S27. Arribas-Hernández, L., Bressendorff, S., Hansen, M.H., Poulsen, C., Erdmann, S. and Brodersen, P. (2018) An m$^6$A-YTH module controls developmental timing and morphogenesis in Arabidopsis. *Plant Cell*, **30**, 952-967.

S28. Scutenaire, J., Deragon, J.M., Jean, V., Benhamed, M., Raynaud, C., Favory, J.J., Merret, R. and Bousquet-Antonelli, C. (2018) The YTH domain protein ECT2 is an m$^6$A reader required for normal trichome branching in Arabidopsis. *Plant Cell*, **30**, 986-1005.

S29. Wei, L.H., Song, P., Wang, Y., Lu, Z., Tang, Q., Yu, Q., Xiao, Y., Zhang, X., Duan, H.C. and Jia, G. (2018) The m$^6$A reader ECT2 controls trichome morphology by affecting mRNA stability in Arabidopsis. *Plant Cell*, **30**, 968-985.

S30. Kang, H.J., Jeong, S.J., Kim, K.N., Baek, I.J., Chang, M., Kang, C.M., Park, Y.S. and Yun, C.W. (2014) A novel protein, Pho92, has a conserved YTH domain and regulates phosphate metabolism by decreasing the mRNA stability of *PHO4* in *Saccharomyces cerevisiae*. *Biochem. J.*, **457**, 391-400.

S31. Wang, X., Lu, Z., Gomez, A., Hon, G.C., Yue, Y., Han, D., Fu, Y., Parisien, M., Dai, Q., Jia, G. *et al.* (2014) N$^6$-methyladenosine-dependent regulation of messenger RNA stability. *Nature*, **505**, 117-120.

S32. Bengtson, M.H. and Joazeiro, C.A. (2010) Role of a ribosome–associated E3 ubiquitin ligase in protein quality control. *Nature*, **467**, 470-473.

S33. Kashima, I., Takahashi, M., Hashimoto, Y., Sakota, E., Nakamura, Y. and Inada, T. (2014) A functional involvement of ABCE1, eukaryotic ribosome recycling factor, in nonstop mRNA decay in *Drosophila melanogaster* cells. *Biochimie*, **106**, 10-16.

S34. Shao, S., von der Malsburg, K. and Hegde, R.S. (2013) Listerin-dependent nascent protein ubiquitination relies on ribosome subunit dissociation. *Mol. Cell*, **50**, 637-648.

S35. Coaker, G., Zhu, G., Ding, Z., Van Doren, S.R. and Staskawicz, B. (2006) Eukaryotic cyclophilin as a molecular switch for effector activation. *Mol. Microbiol.*, **61**, 1485-1496.

S36. Davis, T.L., Walker, J.R., Campagna-Slater, V., Finerty, P.J., Paramanathan, R., Bernstein, G., MacKenzie, F., Tempel, W., Ouyang, H., Lee, W.H. *et al.* (2010) Structural and biochemical characterization of the human cyclophilin family of peptidyl-prolyl isomerases. *PLOS Biol.*, **8**, e1000439.

S37. Haendler, B., Keller, R., Hiestand, P.C., Kocher, H.P., Wegmann, G. and Movva, N.R. (1989) Yeast cyclophilin: isolation and characterization of the protein, cDNA and gene. *Gene*, **83**, 39-46.

S38. Kim, D.H. and Sabatini, D.M. (2004) Raptor and mTOR: subunits of a nutrient-sensitive complex. *Curr. Top. Microbiol. Immunol.*, **279**, 259-270.

S39. Schepetilnikov, M., Dimitrova, M., Mancera-Martínez, E., Geldreich, A., Keller, M. and Ryabova, L.A. (2013) TOR and S6K1 promote translation reinitiation of uORF-containing mRNAs via phosphorylation of eIF3h. *EMBO J.*, **32**, 1087-1102.

S40. Van Leene, J., Han, C., Gadeyne, A., Eeckhout, D., Matthijs, C., Cannoot, B., De Winne, N., Persiau, G., Van De Slijke, E., Van de Cotte, B. *et al.* (2019) Capturing the phosphorylation and protein interaction landscape of the plant TOR kinase. *Nat. Plants*, **5**, 316-327.

S41. Paul, V.D., Mühlenhoff, U., Stümpfig, M., Seebacher, J., Kugler, K.G., Renicke, C., Taxis, C., Gavin, A.C., Pierik, A.J. and Lill, R. (2015) The deca-$GX_3$ proteins Yae1-Lto1 function as adaptors recruiting the ABC protein Rli1 for iron-sulfur cluster insertion. *eLife*, **4**, e08231.

S42. Zhai, C., Li, Y., Mascarenhas, C., Lin, Q., Li, K., Vyrides, I., Grant, C.M. and Panaretou, B. (2014) The function of ORAOV1/LTO1, a gene that is overexpressed frequently in cancer: essential roles in the function and biogenesis of the ribosome. *Oncogene*, **33**, 484-494.

S43. Xu, X., Wan, W., Jiang, G., Xi, Y., Huang, H., Cai, J., Chang, Y., Duan, C.G., Mangrauthia, S.K., Peng, X. *et al.* (2019) Nucleocytoplasmic trafficking of the *Arabidopsis* WD40 repeat protein XIW1 regulates ABI5 stability and abscisic acid responses. *Mol. Plant*, **12**, 1598-1611.

S44. Zhu, G., Chang, Y., Xu, X., Tang, K., Chen, C., Lei, M., Zhu, J.K. and Duan, C.G. (2019) EXPORTIN 1A prevents transgene silencing in *Arabidopsis* by modulating nucleo-cytoplasmic partitioning of HDA6. *J. Integra. Plant Biol.*, **61**, 1243-1254.

S45. Kirli, K., Karaca, S., Dehne, H.J., Samwer, M., Pan, K.T., Lenz, C., Urlaub, H. and Görlich, D. (2015) A deep proteomics perspective on CRM1-mediated nuclear export and nucleocytoplasmic partitioning. *eLife*, **4**, e11466.

S46. Kispal, G., Sipos, K., Lange, H., Fekete, Z., Bedekovics, T., Janáky, T., Bassler, J., Aguilar Netz, D.J., Balk, J., Rotte, C. *et al.* (2005) Biogenesis of cytosolic ribosomes requires the essential iron-sulphur protein Rli1p and mitochondria. *EMBO J.*, **24**, 589-598.

S47. Yarunin, A., Panse, V.G., Petfalski, E., Dez, C., Tollervey, D. and Hurt, E.C. (2005) Functional link between ribosome formation and biogenesis of iron-sulfur proteins. *EMBO J.*, **24**, 580-588.

S48. Hernando, Y., Carter, A.T., Parr, A., Hove-Jensen, B. and Schweizer, M. (1999) Genetic analysis and enzyme activity suggest the existence of more than one minimal functional unit capable of synthesizing phosphoribosyl pyrophosphate in *Saccharomyces cerevisiae*. *J. Biol. Chem.*, **274**, 12480-12487.

S49. Hansen, B.G., Kliebenstein, D.J. and Halkier, B.A. (2007) Identification of a flavin-monooxygenase as the *S*-oxygenating enzyme in aliphatic glucosinolate biosynthesis in Arabidopsis. *Plant J.*, **50**, 902-910.

S50. Rutherford, S. and Moore, I. (2002) The *Arabidopsis* Rab GTPase family: another enigma variation. *Curr. Opin. Plant Biol.*, **5**, 518-528.

S51. Speth, E.B., Imboden, L., Hauck, P. and He, S.Y. (2009) Subcellular localization and functional analysis of the Arabidopsis GTPase RabE. *Plant Physiol.*, **149**, 1824-1837.

S52. Geng, J., Nair, U., Yasumura-Yorimitsu, K. and Klionsky, D.J. (2010) Post-Golgi Sec proteins are required for autophagy in *Saccharomyces cerevisiae*. *Mol. Biol. Cell*, **21**, 2257-2269.

S53. Lepore, D., Spassibojko, O., Pinto, G. and Collins, R.N. (2016) Cell cycle–dependent phosphorylation of Sec4p controls membrane deposition during cytokinesis. *J. Cell Biol.*, **214**, 691-703.

S54. Khan, M., Imran, Q.M., Shahid, M., Mun, B.G., Lee, S.U., Khan, M.A., Hussain, A., Lee, I.J. and Yun, B.W. (2019) Nitric oxide- induced *AtAO3* differentially regulates plant defense and drought tolerance in *Arabidopsis thaliana*. *BMC Plant Biol.*, **19**, 602.

S55. Seo, M., Aoki, H., Koiwai, H., Kamiya, Y., Nambara, E. and Koshiba, T. (2004) Comparative studies on the *Arabidopsis* aldehyde oxidase (*AAO*) gene family revealed a major role of *AAO3* in ABA biosynthesis in seeds. *Plant Cell Physiol.*, **45**, 1694-1703.

S56. Cheshmazar, N., Dastmalchi, S., Terao, M., Garattini, E. and Hamzeh-Mivehroud, M. (2019) Aldehyde oxidase at the crossroad of metabolism and preclinical screening. *Drug Metab. Rev.*, **51**, 428-452.
